# A synthetic peptide mimic kills *Candida albicans* and synergistically prevents infection

**DOI:** 10.1101/2023.09.25.559234

**Authors:** Sebastian Schaefer, Raghav Vij, Jakob L. Sprague, Sophie Austermeier, Hue Dinh, Peter R. Judzewitsch, Eric Seemann, Britta Qualmann, Amy K. Cain, Nathaniel Corrigan, Mark S. Gresnigt, Cyrille Boyer, Megan D. Lenardon, Sascha Brunke

## Abstract

More than two million people worldwide are affected by life-threatening, invasive fungal infections annually. *Candida* species are the most common cause of nosocomical, invasive fungal infections and are associated with mortality rates above 40%. Despite the increasing incidence of drug-resistance, the development of novel antifungal formulations has been limited. Here we investigate the antifungal mode of action and therapeutic potential of positively charged, synthetic peptide mimics to combat infections by *Candida albicans*. These synthetic polymers cause stress to the endoplasmic reticulum and affect protein glycosylation, a distinct mode of action compared to currently approved antifungal drugs. The most promising polymer composition caused damage to the mannan layer of the cell wall, with additional membrane-disrupting activity. The synergistic combination of the polymer with caspofungin prevented infection of human epithelial cells *in vitro*, improved fungal clearance by human macrophages, and significantly increased host survival in a *Galleria mellonella* model of systemic candidiasis. Additionally, prolonged exposure of *C. albicans* to the synergistic combination of polymer and caspofungin did not lead to the evolution of resistant strains *in vitro*. Together, this work highlights the enormous potential of these synthetic peptide mimics to be used as novel antifungal formulations as well as adjunctive antifungal therapy.

## Introduction

Modern medicine often relies on invasive medical interventions or drugs which can compromise the patient’s immune system. An unfortunate consequence of these undeniably successful treatments for life-threatening diseases like cancer are severe infections caused by opportunistic pathogens.^1, 2^ Among these opportunists are fungal pathogens, including *Candida*, *Aspergillus*, *Cryptococcus*, and *Pneumocystis* species.^1–3^ More recently, increasing numbers of opportunistic fungal infections caused by *Aspergillus*, *Mucorales*, and *Candida* species have been observed in COVID-19 patients with severe respiratory syndromes in intensive care units.^4^ These and other factors result in over 2 million invasive fungal infections annually worldwide, with alarmingly high mortality rates and more than 1.5 million deaths.^3^

*Candida* spp. are the fourth most common cause of hospital-acquired infections, and mortality rates from systemic *Candida* infections exceed 40%, even with antifungal intervention.^3, 5^ Among all *Candida* species, *Candida albicans* accounts for around 50% of *Candida* bloodstream infections.^3, 6^ Novel pathogenic species, such as the multi-drug resistant *Candida auris* have emerged, potentially through adaptations to higher ambient temperature due to global climate change.^7, 8^ Indeed, *C. auris* and *C. albicans* were listed as two of the four critical group pathogens in the World Health Organization’s first-ever fungal priority pathogens list, emphasising the need for new treatment options.^9^

There are currently only four classes of antifungal drugs approved for treatment of invasive *Candida* infections – azoles (*e. g.*, fluconazole), polyenes (*e. g.*, amphotericin B), echinocandins (*e. g.*, caspofungin), and flucytosine.^10^ Their application is limited by undesired drug-drug interactions (azoles), detrimental off-target side-effects (polyenes), and the increasing occurrence of drug resistance (azoles, flucytosine, and echinocandins).^10, 11^ Resistance of *Candida* spp. occurs mainly due to target over-expression, modification of the drug target, or upregulation of drug-efflux pumps.^11^ The urgency to find new treatment options against *Candida* spp. was highlighted by the Centers for Disease Control and Prevention’s (CDC) 2019 classification of *Candida* spp. as a serious threat to human health with equal standing to multidrug-resistant bacteria such as *Pseudomonas aeruginosa*.^12^ However, discovering novel targets for antifungal drugs is complicated by the evolutionary similarity of eukaryotic human and fungal cells, and the antifungal development pipeline is dominated by compounds from established classes, which are likely to result in similar complications.^10, 13^ Exceptions are fosmanogepix and ibrexafungerp which are first-in-class and undergoing clinical trials,^14, 15^ but ultimately the emergence of resistance and clinical success for these new classes remain to be seen.^14^ Combination therapy can decrease the development of resistance or re-sensitise resistant strains by acting on multiple targets.^16–18^ It can also reduce toxicity to the host by decreasing required drug concentrations.^16^

In nature, antifungal peptides (AFPs) prevent and combat fungal infections in all domains of life.^19^ Most interact with the fungal cell membrane, damaging the cell wall or membrane or causing intracellular stress.^19^ Employing those potent natural effectors as a drug is hampered by several issues; the membrane-active AFPs are often toxic to the expression host in biotechnological synthesis, chemical synthesis of peptides is expensive and complicated by their sequence specificity, and susceptibility to host proteases generally limits the applicability of AFPs.^19^ There has been interest in synthetic polymers designed to replicate the antifungal function of AFPs to circumvent the limitations of AFPs.^20–23^

Our group previously synthesised and screened a library of synthetic polyacrylamides inspired by AFP structures for activity against *C. albicans* and biocompatibility.^20^ We identified polymers that outperformed amphotericin B in terms of their therapeutic index against *C. albicans in vitro*.^20^ In the current work, we determined the novel mode of action of our most promising polymer compositions, and investigated the *in vitro* and *in vivo* therapeutic potential of synergistic combinations of our polymers with existing antifungals with a view to enhancing efficacy, minimising toxicity and preventing the emergence of antifungal drug resistance.

## Results and discussion

### Synthesis and characterisation of amphiphilic polyacrylamides that mimic AFPs

Inspired by physicochemical properties of antimicrobial peptides, we previously synthesised random acrylamide copolymers using photo-induced reversible-deactivation radical polymerisation.^20, 24^ The polymer characteristics that conferred the highest activity against *C. albicans* and best biocompatibility with mammalian host cells were short polymers with a degree of polymerisation (*X*_n_) of 20 and an optimal balance of hydrophilic to hydrophobic groups.^20^ Here, we synthesised four ternary polyacrylamides with these characteristics (**Figure 1** **and Table 1**): LP (linear, pentyl), LH (linear, heptyl), CB (cyclic, benzyl), and CX (cyclic, hexyl), named by their distinct hydrophobic features. The successful synthesis and purification were confirmed by ^1^H nuclear magnetic resonance (NMR) spectroscopy and refractive index-based size exclusion chromatography (SEC). We achieved nearly complete monomer conversion of >98% and observed narrow, unimodal molecular weight distributions with a dispersity (*Ð*) between 1.09 and 1.12 (**Supplementary Figures S1-S15**).

**Figure 1.**
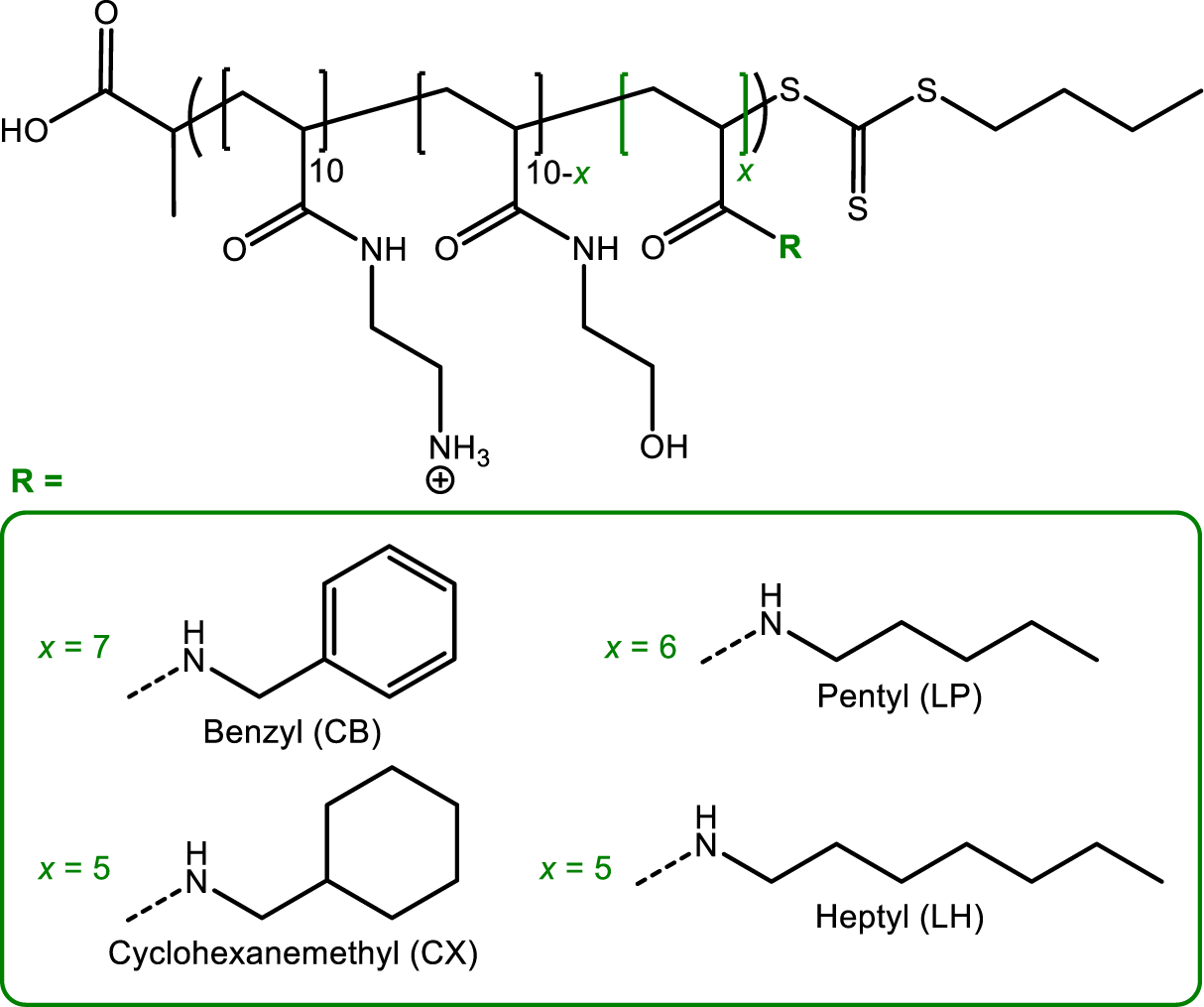
– Chemical structures of polyacrylamides synthesised for this study. R represents the different side chains and *x* indicates the targeted number of hydrophobic residues within the molecule.

**Table 1.**
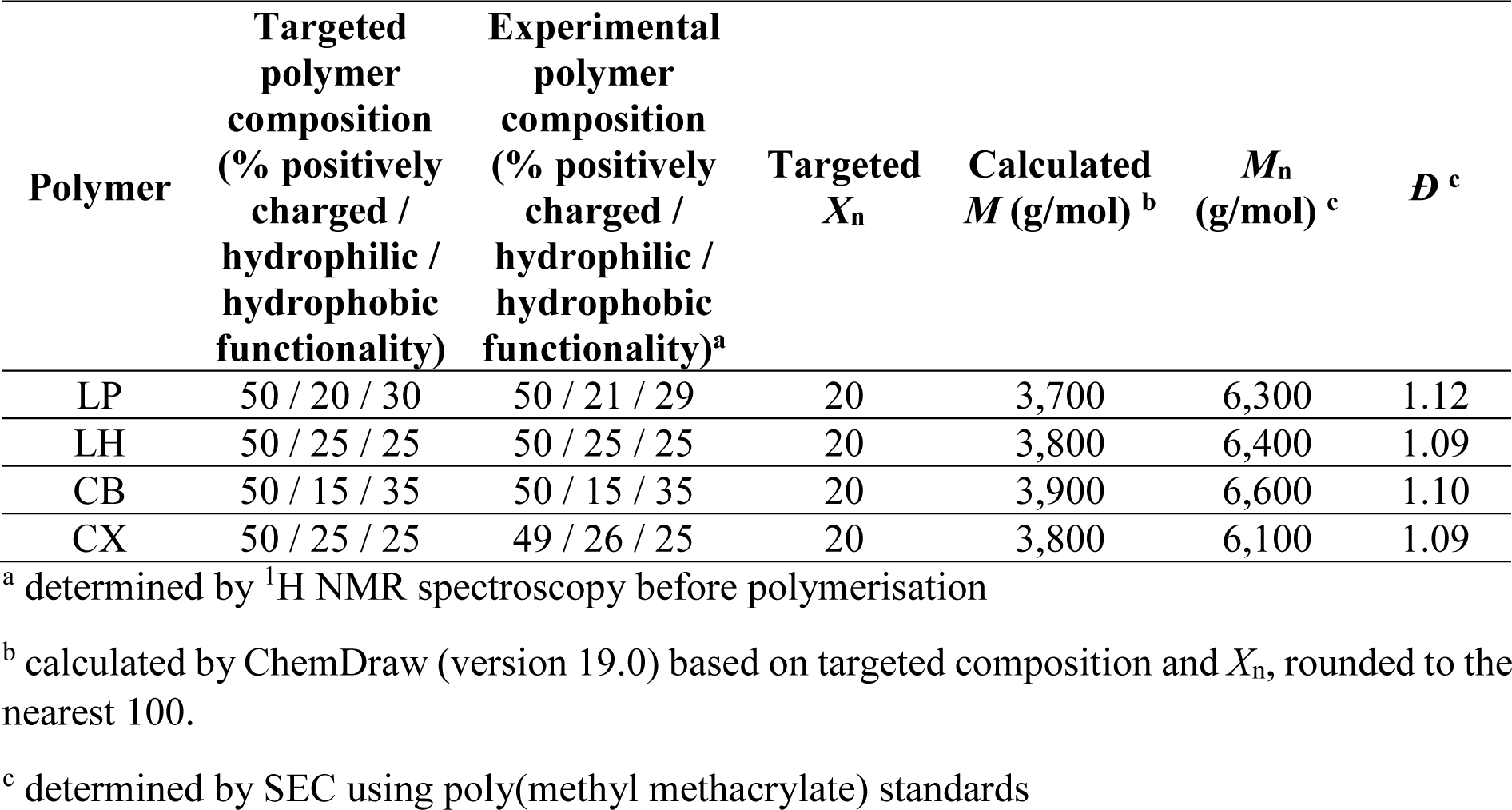
– Composition, targeted degree of polymerisation (*X*_n_), calculated molecular weight (*M*), experimental number-average molecular weight (*M*_n_) and dispersity (*Ð*) of polymers employed in this study.

### Polyacrylamides are active against drug-resistant, clinical *C. albicans* isolates

Our previous work showed that our polymers were active against *C. albicans* and other ascomycetes including *Candida glabrata* (*Nakaseomyces glabrata*), *Candida krusei* (*Pichia kudriavzevii*) and *Saccharomyces cerevisiae*, as well as the basidiomycete *Cryptococcus neoformans*.^20^ Despite a high tolerance of *C. neoformans* towards amphotericin B (AmpB) and fluconazole, and intrinsic resistance of *S. cerevisiae* and *C. glabrata* towards fluconazole, all were susceptible to the four candidate polymers.^20^ This suggested a mode of action that is different to AmpB and fluconazole. To explore this further, the minimum inhibitory concentration (MIC, growth inhibition of >90% at 24 h) of the polymers against antifungal drug-resistant strains of *C. albicans* (**Table 2**) was assessed using. slightly modified Clinical and Laboratory Standards Institute (CLSI) guidelines.^25, 26^

**Table 2.** – Antifungal activity of polymers and selected antifungal drugs against *C. albicans* clinical isolates.

| *C. albicans* strain | Phenotype | Minimum inhibitory concentration<sub>24h, 90%</sub> (µg/mL) |  |  |  |  |  |  |
| --- | --- | --- | --- | --- | --- | --- | --- | --- |
|  |  | Polymers |  |  |  | Antifungal drugs |  |  |
|  |  | LP | LH | CB | CX | AmpB | Flu | Cas |
| SC5314 | Wild-type | 64-128 | 16-32 | 64 | 64 | 0.5-1 | 0.25-0.5 | 0.25-0.5 |
| 110.12 | Gsc1-mutation | 256 | 16 | 128 | 128 | 1-2 | >8 | 4-8 |
| EU0108 | Erg11-/Erg5-mutation, enhanced Cdr efflux | 64 | 8-16 | 16-32 | 32-64 | 2 | >8 | 0.5-1 |
| EU0012 | Erg3-mutation | 128-256 | 16-32 | 64-128 | 64-128 | 2-4 | >8 | 0.5-1 |
| EU1008 | Erg3-/Erg11-mutation | 32-64 | 8-16 | 32 | 16-32 | 2 | >8 | 0.5-1 |
| EU0136 | Erg6-deficient | 256 | 16-32 | 128 | 64-128 | 4 | >8 | 0.5 |
| EU0992 | Enhanced Cdr-efflux | 64-128 | 16 | 64-128 | 64-128 | 1-2 | >8 | 0.5-1 |
| EU0989 | Enhanced Cdr-efflux | 128 | 8-16 | 64 | 64-128 | 1-2 | >8 | 0.5-1 |
| EU0981 | Enhanced Mdr-efflux | 64 | 8-16 | 64 | 32-64 | 1-2 | >8 | 0.25-0.5 |
| EU0999 | Enhanced Mdr-efflux | 64-128 | 8-16 | 64-128 | 32-64 | 1-2 | >8 | 0.25-0.5 |

*Note:* Minimum inhibitory concentrations were determined by the concentration at which the respective compound inhibited >90% of fungal growth compared to an untreated control after 24 h at 35 °C. Red indicates resistance, orange indicates increased tolerance (up to two-fold increase in MIC), and green indicates a decreased MIC compared to wild-type. AmpB: Amphotericin B, Flu: Fluconazole, Cas: Caspofungin.

*C. albicans* strain 110.12 is resistant to azoles and caspofungin, the latter due to a mutation in the echinocandin target Gsc1, an essential β-1,3-glucan synthase subunit.^27^ The MICs for the polymers LP, CB and CX were only slightly increased compared to wild-type, and the clinical isolate was at least as susceptible as the wild-type to polymer LH.

AmpB-tolerant (EU0108, EU1008) or -resistant (EU0012, EU0136) strains have various mutations in enzymes involved in the biosynthesis of ergosterol (Erg3, Erg5, Erg6, Erg11), the target of AmpB.^28, 29^ The strains are also azole resistant, which has been attributed to increased drug efflux at least in strain EU0108.^28, 29^ However, all polymers were active against those clinical isolates with even decreased MICs against the AmpB-tolerant strains. LH was fully active against the AmpB-resistant strains. A slight increase in MIC was observed for LP, CB, and CX.

The fluconazole-resistant *C. albicans* strains (EU0992, EU0989, EU0981, EU0999) have mutations in *CDR* or *MDR* resulting in enhanced drug efflux.^30^ The antifungal activity of the polymers was not affected by those mutations, suggesting that they are not transported out by Cdr- or Mdr-related efflux pumps.

Overall, each antifungal drug-resistant *C. albicans* strain tested was as susceptible to LH, if not more, as the wild-type. This indicates a distinct mode of action of LH. The different activity pattern of the other polymers against drug-resistant strains may indicate slight differences in modes of action between the polymers. We therefore investigated the potentially novel modes of action of our polymers.

### Transcript profiling to investigate the mode of action of the antifungal polymers

We compared the transcriptome of *C. albicans* cells grown for 1 h in the presence of sub-inhibitory concentration (0.5×MIC) of the antifungal polymers to a no treatment control. A non-active polymer, poly(hydroxyethyl acrylamide) (poly-HEA), was also included for comparison (chemical characterisation for poly-HEA in **Supplementary Figures S14-15 and Table S1**).

An overview of the global transcriptomic differences of *C. albicans* in response to the polymers was gained through hierarchical clustering and principal component analyses of normalised gene expression data. These analyses showed that the transcriptome of cells exposed to the antifungal polymers LP, CB, and CX clustered together (**Supplementary Figure S16**). Surprisingly, the hierarchical clustering revealed that the transcriptomic patterns for these three polymers were more similar to the non-toxic poly-HEA (**Supplementary Figure S16A**), while our most active candidate LH clustered separately after applying either of the two statistical methods (**Supplementary Figure S16A and B**).

Next, we looked for biological functions and pathways where there was an over-representation of up- or down-regulated genes associated with the function or pathway in the datasets by performing Gene Ontology (GO) term^31^ and Kyoto Encyclopedia of Genes and Genomes (KEGG) pathway enrichment^32^ analyses (Figure 2). Similar to our clustering analyses, we noted that the non-toxic poly-HEA differed from the four antifungal polymers, and that LH showed the most distinct pattern with the highest number of differentially expressed genes.

**Figure 2.**
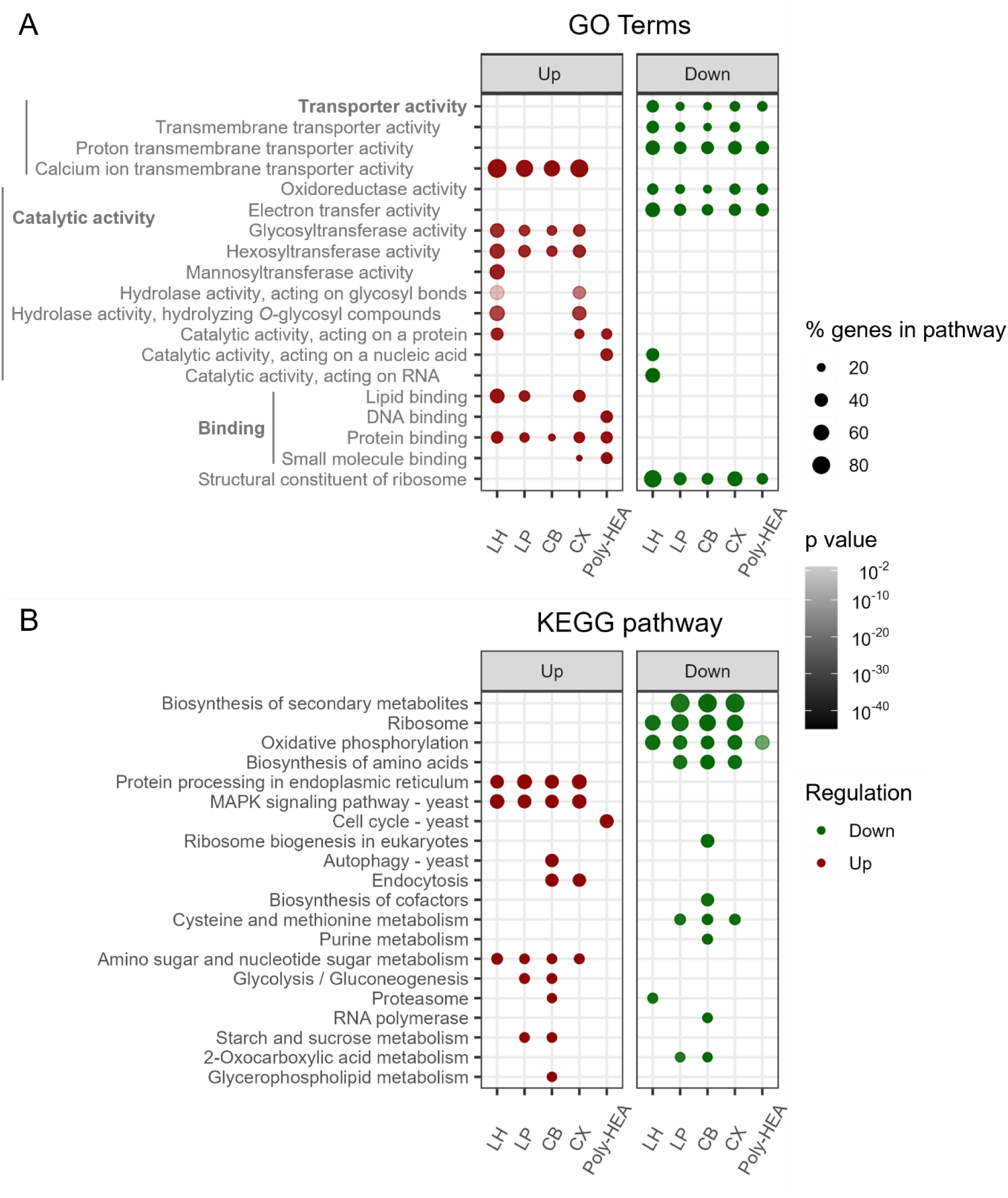
– Antifungal polymers cause transcriptomic responses associated with impaired protein glycosylation, membrane stress, and cell wall damage in *C. albicans*. (A) GO term enrichment analysis, based on molecular function,^31^ and (B) KEGG pathway enrichment analysis^32^ of RNA microarray data after treating *C. albicans* SC5314 for 1 h at 30 °C with sub-inhibitory polymer concentrations (0.5×MIC for LH, LP, CB and CX, 128 µg/mL for poly-HEA). Statistically significantly up- (red) or downregulated (green) gene groups associated with GO terms and KEGG pathways compared to the untreated control are shown. Diameter of circles reflects the percentage of genes differentially regulated in the associated pathway or term and the shading represents the adjusted p-value. GO terms are additionally ordered by their assigned parental processes, displayed by indentation.

Functional GO term enrichment analyses on the sets of differentially expressed genes were performed with GO term finder tool at *Candida* Genome Database^31^ (Figure 2A). GO terms relating to membrane stress were enriched in the upregulated gene set. This included the GO term calcium ion transmembrane transport, suggesting increased Ca^2+^ influx and membrane damage. GO terms associated with transporter activity were enriched in the downregulated gene set, indicating general stress and metabolic arrest of the cells. The GO term oligopeptide transmembrane transporter activity was enriched in the set of genes downregulated following treatment with LH. Together with the overrepresentation of genes involved in lipid and protein binding, this supports the hypothesis of *C. albicans* sensing a toxic, peptide-like structure with amphiphilic properties. Additionally, GO terms associated with glycosylation processes were enriched in the upregulated gene set, specifically hydrolase activity against *O*-glycosyl compounds and hexosyltransferase activity – a parental gene set of mannosyltransferase activity. In contrast, treatment with the non-toxic poly-HEA did not elicit a significant enrichment of these glycosylation-related and membrane damage-indicating GO terms. Instead, we found an enrichment of GO terms associated with DNA, protein, and small molecule binding in the upregulated gene set, which may indicate that *C. albicans* reacts to a small, peptide-like molecule, however, without any effect on glycosylated proteins or the cell membrane.

KEGG pathway enrichment analysis was also used to associate gene expression data with specific pathways.^32, 33^ One KEGG pathway, which was significantly enriched in upregulated genes under all polymer treatments (except poly-HEA), was protein processing in endoplasmic reticulum (ER) (Figure 2B). A more detailed look (Figure 3A) revealed that *C. albicans* cells treated with LH strongly upregulated genes associated with glycosylation and ER-associated degradation of misfolded protein (ERAD), suggesting a disruption in the correct glycosylation of proteins. In support of this hypothesis, we also observed upregulated genes in the *N*-glycan biosynthesis pathway (Figure 3B). KEGG pathway enrichment analysis further revealed a transcriptional signature expected from damage to the cell wall and membrane (Figure 2B), particularly with polymer LH. This includes an overrepresentation of the KEGG pathways glycosylphosphatidylinositol (GPI)-anchor biosynthesis and glycerolipid biosynthesis (shown for LH-treatment in **Supplementary Figure S17**) in the upregulated gene set, and steroid biosynthesis (**Supplementary Figure S18**) in the downregulated gene set.

**Figure 3.**
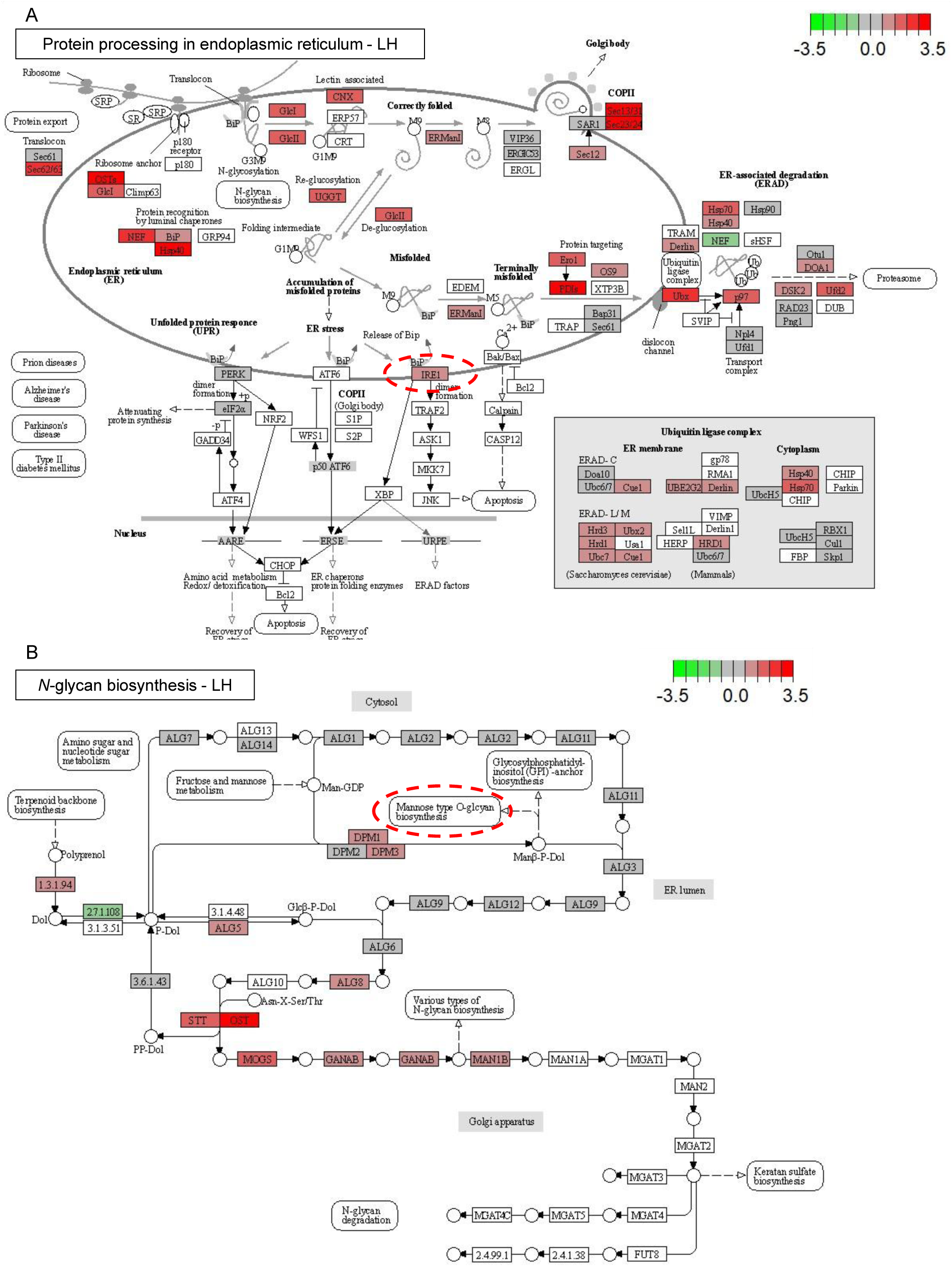
– KEGG pathway maps^32, 33^ for (A) protein processing in the endoplasmic reticulum and (B) *N*-glycan biosynthesis after treating *C. albicans* SC5314 for 1 h at 30 °C with 0.5×MIC for LH. Boxes represent genes and their regulation within the specific pathway, where red shading indicates upregulated, green downregulated, and grey non-regulated in the presence of LH, while white indicates no data. The scale bars represent the log2 fold change in gene expression compared to untreated cells. Red circles highlight the gene and sub-pathway that have been found to be crucial for *C. albicans* to respond to and survive LH-induced stress in the deletion mutant screen.

The mitogen-activated protein kinase (MAPK) stress response signalling pathway was enriched in upregulated genes upon treatment with all antifungal polymers (Figure 2B, LH-treatment in **Supplementary Figure S19**).^34, 35^ This indicates that the antifungal polymers cause responses typical for cell wall and osmotic stress or starvation. Upregulation of genes in these pathways leads to cell cycle arrest and is associated with filamentation, cell wall remodelling, and osmotic stress response.^35^ Treatment with antifungal polymers also led to enrichment of the KEGG pathways protein synthesis (*e. g.*, ribosome, shown for LH in **Supplementary Figure S20**), biosynthesis of amino acids, and oxidative phosphorylation in the downregulated gene set. Together with MAPK signalling, this suggests metabolic arrest of the *C. albicans* cells by environmental stress. Upon treatment with the non-toxic poly-HEA, we found very few pathways with an enrichment of up- or downregulated genes. This highlights the specificity of the pathway responses by protein processing in ER and MAPK signalling for antifungal activity, rather than presence of the polymers alone. In addition, the gene expression profiles of *C. albicans* exposed to our polymers are very distinct compared to exposure to polyenes, azoles or echinocandins.^36^ This again suggests a unique mode of action of these synthetic polymers.

Currently approved antifungals have specific cellular targets, and mutations in these enzymes are one of the main reasons for development of resistance.^10^ In contrast, antimicrobial peptides have multiple modes of action, decreasing the likelihood for the development of resistance.^37^ For example, a *Musca domestica* (housefly) AFP triggers responses in *C. albicans* that include reaction to oxidative stress, cell wall and membrane maintenance, protein synthesis, and energy metabolism.^38^ A bacterium-derived, membranolytic antifungal lipopeptide (jagaricin) similarly induces a broad transcriptional response, comprising the upregulation of cell wall organisation/biogenesis and calcium ion transmembrane transport genes, and downregulation of transmembrane transport for substances such as oligopeptides.^30^ Our antifungal polymers induced a similar transcriptional response in *C. albicans*. Altogether, these data led us to hypothesise that the antifungal polymers likely target multiple processes in *C. albicans* – they damage the cell wall and permeabilise the cell membrane, and they target protein glycosylation and thereby induce ER stress.

### *O*-mannosylation, calcineurin, MAPK signalling, and a phosphoinositide regulator are required for *C. albicans* to survive polymer exposure

To gain further insights into the target of the polymers, we screened selected deletion mutants of *C. albicans* for their growth at sub-inhibitory polymer concentrations (LP, LH, CB, CX). Mutants were chosen to represent suspected target and resistance pathways, including cell wall organisation and stress response, membrane composition and inositol signalling, protein glycosylation (*O*- and *N*-linked mannosylation), calcineurin pathway, osmotic and oxidative stress response (MAPK signalling), drug efflux, and polyamine uptake (**Supplementary Table S2**). Growth curves were compared based on the time to reach half-maximal absorption (**Supplementary Table S2**). This growth speed index is positive for beneficial mutations (green shading in **Supplementary Table S2**) and negative for detrimental ones (red shading in **Supplementary Table S2**).

Deletion of genes important for cell wall organisation and stress response had little effect on growth in the presence of polymers (**Supplementary Table S2**), with the exception of *ire1*, which showed no growth in the presence of LH. *IRE1* encodes a protein kinase of the unfolded protein response and cell wall organisation.^39^ Interestingly, *IRE1* was also upregulated in the presence of LH (Figure 3A).

Among the mutants for genes relevant for membrane composition and inositol signalling (**Supplementary Table S2**), *inp51* – lacking a phosphatase involved in maintenance of phosphoinositide levels and thus cell wall and membrane integrity^40^ – exhibited no growth in the presence of all polymers. *INP51* deletion has been shown to increase susceptibility to cell wall-active compounds.^40^ Deletion of *ERG5*, involved in ergosterol biosynthesis, did not change the susceptibility to the polymers. This agrees well with the susceptibility of clinical AmpB-tolerant isolates towards the polymers (**Table 2**) and indicates that ergosterol biosynthesis likely does not affect the polymers’ mode of action.

Deletion of genes contributing to *O*-linked protein mannosylation (*PMT1*, *PMT3*, *PMT4*, *PMT5*)^41^ consistently increased susceptibility of *C. albicans* (**Supplementary Table S2**), suggesting that these activities are involved in the response to all antifungal polymers. In contrast, gene deletions affecting *N*-linked mannosylation (*MNN13*, *MNN14*, *MNN15*, *MNN22*, *MNN4*, *MNN9*, *MNS1*, *MNT4*, *OCH1*)^41^ resulted in no change in susceptibilities.

Calcineurin and MAPK signalling are crucial in fungal development and response to environmental stress.^42^ Two calcineurin pathway deletion mutants (*crz1*, *mid1*) showed no growth with normally sub-inhibitory polymer concentrations (**Supplementary Table S2**), and their genes’ expression was also upregulated in *C. albicans* exposed to the antifungal polymers, but not poly-HEA. Two MAPK signalling deletion mutants (*hog1*, *pbs2*) also showed no growth in the presence of polymers. *HOG1* similarly has shown upregulation after exposure to the antifungal polymers, most prominently for LH and CX (Figure 2B**, Supplementary Figure S19**). Hence, the antifungal polymers seem to cause stress in *C. albicans*, and both calcineurin and MAPK pathway appear to be essential for fungal survival.

In agreement with the clinical drug-efflux mutants (**Table 2**), neither gain-of-function (in *MRR1* and *TAC1*) nor deletion (of *MRR1*, *SNQ2*, *TAC1*) of genes involved in drug efflux had a consistent impact on susceptibility.

The AFP histatin 5 is actively transported across the fungal membrane by polyamine transporters like Dur31 in *C. albicans* to exert its intracellular antifungal effect.^43^ Our synthetic polymers share characteristics like cationic charge with both histatin 5 and polyamines, and their presence led to upregulation of *DUR31*. However, deletion of *DUR31* or the related *DUR35*^44^ resulted in a slightly increased (*dur31*) or unchanged susceptibility (*dur35*) (**Supplementary Table S2**), suggesting that the polymers are not primarily taken up by those transporters.

We also investigated selected *C. glabrata* (*Nakaseomyces glabrata*) mutants (marked blue in **Supplementary Table S2**). While none of the tested *C. albicans* mutations were beneficial, the deletion of *C. glabrata ERG5* and *INO2* increased tolerance to all antifungal polymers. Both genes are associated with membrane composition and inositol signalling. *C. glabrata cdr1*, lacking a drug efflux pump-encoding gene, also showed increased growth with polymers, except LH. In contrast, deletion of *MNN2*, coding for an *N*-linked mannosyltransferase, had the highest benefit for LH-treated *C. glabrata* cells. Despite these slight differences, our data still suggests a very similar mode of action against *C. glabrata*. Like in *C. albicans*, deletion of genes encoding proteins involved in membrane composition and inositol signalling (*INP51*, *INP53*, *ISC1*), *O*-linked protein mannosylation (*PMT1*, *PMT2*) and MAPK signalling (*PBS2*) were more susceptible to the polymers, while the deletion of genes coding for proteins in the calcineurin pathway (*CNA1*, *CRZ1*, *MID1*) resulted in no growth in the presence of antifungal polymers.

In sum, the mutant screening was in agreement with our analyses of the transcriptome and suggests a mode of action connected to protein glycosylation and general stress with some similarities (involvement of MAPK and calcineurin signalling) to a previous *C. albicans* mutant screen with the antifungal lipopeptide jagaricin, for which disruption of membrane integrity was suggested as primary mode of action.^30^ This suggests a similar effect by the synthetic polymers, especially as our gene expression analyses also showed possible interference with the plasma membrane (Figure 2).

### Polymer LH lyses *C. albicans* cell membranes

The polymers are inspired by antimicrobial peptides, which lyse bacterial and fungal membranes.^45^ Membrane damage by the synthetic polymers was also suggested by gene expression analyses (Figure 2) and mutant screening (**Supplementary Table S2**). We therefore investigated the membranolytic potential of polymer LH with a *C. albicans* strain constitutively expressing GFP in the cytosol (Figure 4). Two antifungal compounds were included – AmpB, which has membranolytic activity,^46, 47^ and tunicamycin, which inhibits *N*-glycosylation of proteins, leading to cell death without membrane lysis.^48^

**Figure 4.**
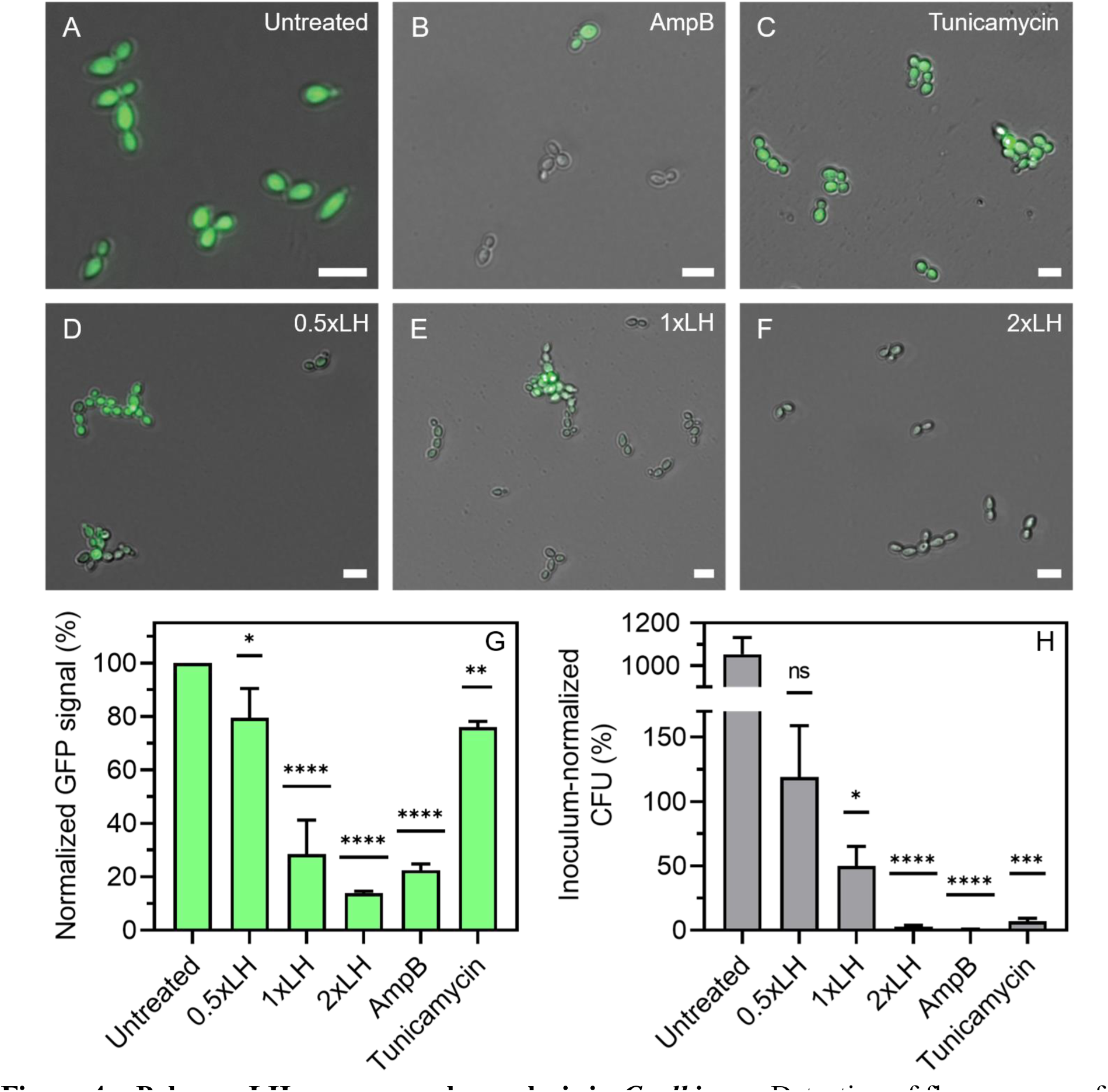
– Polymer LH causes membrane lysis in *C. albicans*. Detection of fluorescence of *C. albicans* expressing cytoplasmic GFP (*ADH1*-GFP) after 6 h at MIC assay conditions for (A) untreated *C. albicans* cells, (B) cells treated with 1×MIC AmpB, (C) 1×MIC tunicamycin, (D) 0.5×MIC polymer LH, (E) 1×MIC polymer LH, and (F) 2×MIC polymer LH to investigate lytic activity of the compounds. (G) The GFP signal was quantified from at least 50 *C. albicans*-GFP cells per replicate (n=3), and then averaged and normalised to the respective untreated control. (H) Colony forming units (CFU) were determined by backplating and normalised to the inoculum. Error bars in G and H represent the standard deviation. Statistical significance in G and H was determined by Dunnett’s ordinary one-way ANOVA multiple comparisons analysis (compared to untreated (100%) in G or inoculum (100%) in H, * p<0.05, ** p<0.005, *** p<0.0005, **** p<0.0001) with ns indicating non-significant. Scale bars represent 10 µm.

Untreated *C. albicans*-GFP cells showed a prominent intracellular GFP signal (Figure 4A), which was absent in AmpB-treated cells (Figure 4B and G) where only 0.3% were viable as determined *via* backplating (Figure 4H). The loss of GFP signal was due to membrane lysis, and not cell death *per se*, as tunicamycin-treated cells showed a less prominent loss of intracellular GFP (Figure 4C), even though only 3% of cells were viable. The polymer LH (Figure 4D-F) led to membranolytic activity, as seen by GFP signal loss, in a concentration-dependent manner with a proportional decrease in cell viability (36% at 1×MIC, 3% at 2×MIC). Interestingly, at MIC and below, we observed cell aggregates, which suggest a change in the surface properties of *C. albicans* cells.

### LH damages mannans attached to cell wall proteins, enhances phagocytosis, and affects yeast-to-hypha transition

Mutant screening and gene expression analyses suggested that glycosylated proteins might be primarily affected by the polymers. A major group of glycosylated proteins in *C. albicans* are the cell wall proteins. These are both *O*- and *N*-mannosylated as they pass through the ER and Golgi on their way to the wall. Once attached to the wall through GPI anchors or Pir linkages, the short *O*-mannan chains remain buried in the inner cell wall layer, whereas the long *N*-mannan chains protrude out from the cell surface forming an outer fibrillar layer of the cell wall (Figure 5A).^49^ To investigate the effects of LH on cell wall structure of *C. albicans*, fungal cells were incubated at sub-inhibitory concentrations for 6 h and analysed by transmission electron microscopy (TEM, Figure 5). Compared to the no-treatment control (Figure 5A), treatment with the non-toxic poly-HEA resulted in no major ultrastructural changes to the cell wall (Figure 5B). A reduction of the outer cell wall *N-*mannan layer was observed after treatment with tunicamycin, an inhibitor of *N*-glycosylation (Figure 5C).^48^ Treatment with a sub-inhibitory concentration of LH (Figure 5D) disrupted the arrangement of the *N*-mannan fibrils, supporting our transcriptome and mutant screen data. At these sub-inhibitory concentrations, LH did not cause any obvious disruption of the membrane.

**Figure 5.**
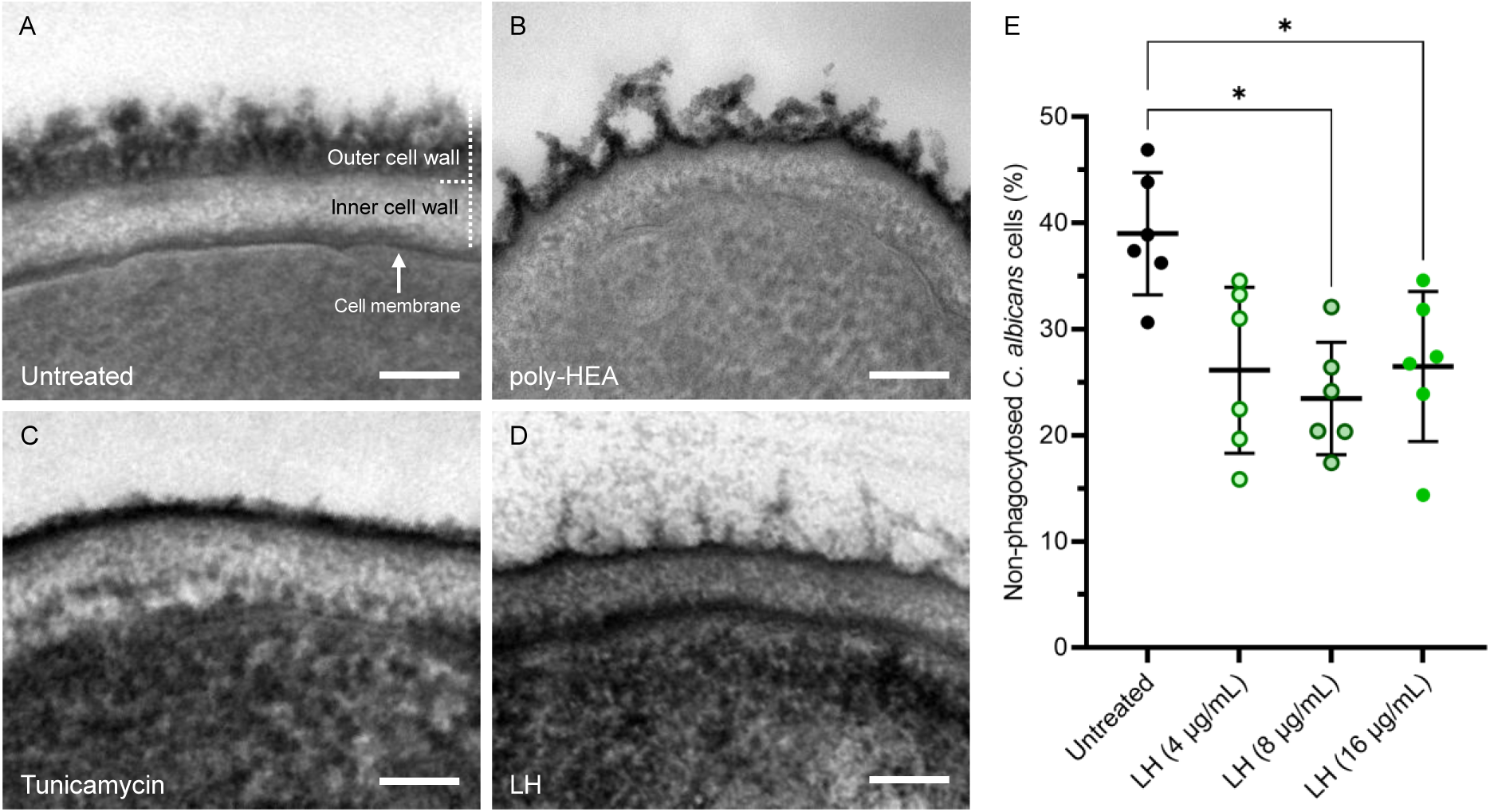
– Cell wall changes in *C. albicans* and immune cell response to LH-treated *C. albicans*. TEM micrographs of the *C. albicans* cell wall after 6 h incubation in SD medium at 30 °C with (A) no additives, (B) with poly-HEA, (C) with tunicamycin, or (D) antifungal polymer LH, at sub-inhibitory concentrations of the antifungal compounds. The scale bars in A-D represent 100 nm. (E) Proportion of *C. albicans* cells not taken up over 15-30 min by human monocyte-derived macrophages. *C. albicans* was pre-treated with LH for one hour before putting it into contact with the macrophages. Statistical significance in E was determined by Dunnett’s repeated measures ANOVA multiple comparisons analysis (*p<0.05).

Components of the *C. albicans* cell wall are also recognised by innate immune cells.^50^ We hypothesised that the LH-induced changes to the cell wall structure (Figure 5D) could also impact immune recognition and clearance by human macrophages. To test this, we challenged *C. albicans* cells that were preincubated with or without polymer LH at sub-inhibitory concentrations with primary human monocyte-derived macrophages (hMDMs). Primary immune cells such as hMDMs show donor-dependent differences in their uptake efficiency. We therefore chose the time point (15 or 30 min) for each donor when 30-50% of untreated *C. albicans* cells were not yet phagocytosed by the specific macrophages. We found that LH pre-treatment significantly increased clearance of *C. albicans* by primary hMDMs, even at sub-inhibitory concentrations (Figure 5E). Concentrations above MIC, however, resulted in decreased clearance (**Supplementary Figure S21A**), most likely due to toxic effects to the hMDMs (**Supplementary Figure S21B**).

It has been previously reported that *C. albicans* mutants with *N*- or *O*-linked glycosylation defects are more efficiently phagocytosed than the wild-type.^51, 52^ With our TEM and transcriptome data, this suggests that LH disrupts the mannan layer and induces cell wall remodelling which increases phagocytosis by macrophages. Thus, *in vivo* application of polymer LH could potentiate fungal clearance by innate immune cells.

The cytokine release by primary human peripheral blood mononuclear cells (PBMCs) to pre-treated *C. albicans* was characterised for the pro-inflammatory cytokines IL-1β, IL-6, and TNF (**Supplementary Figure S22**). The increased clearance by hMDMs (Figure 5E), was reflected by significantly decreased IL-1β, IL-6, and TNF responses of PBMCs at a LH concentration of 4 µg/mL (**Supplementary Figures S22A-C**). Higher LH concentrations resulted in no significant changes in release of IL-1β and TNF and a significant increase of IL-6. The decrease in TNF is in agreement with previous observations on *C. albicans N*-mannosylation defective mutants with impaired recognition by immune cells.^53, 54^ As mannoproteins in *C. albicans* cause a pro-inflammatory immune response,^41, 50^ these data support our notion that polymer LH can act on protein glycosylation.

The cell wall of *C. albicans* is adaptive and constantly remodels, including when *C. albicans*’ transitions from yeast to hyphae, a process that is commonly associated with virulence.^50^ Hence, we tested *C. albicans* under hypha-inducing conditions (37 °C and 5% CO_2_) in the presence of antifungal polymers at sub-inhibitory concentrations over 4, 6 (to measure hypha length), and 24 h (to measure microcolony diameter) (**Supplementary Figure S23**). Treatment with each polymer, especially LH, reduced the hyphae’s length and the diameter of microcolonies. These results underline the therapeutic potential of these polyacrylamides, since the formation of hyphae is a mechanism that drives cell invasion and infection of human epithelial cells by *C. albicans*.^55^

### Polymer LH prevents *in vitro* infection of human epithelial cells by *C. albicans* synergistically with caspofungin or fluconazole

During systemic infection, *C. albicans* hyphae invade the human epithelial barrier, and allow it to spread *via* the bloodstream to distal organs.^56^ The yeast-to-hypha transition is also important for common superficial infection, such as those of the vaginal mucosa, which affect approximately 75% of women worldwide at least once in their lifetime.^57, 58^ To investigate the therapeutic potential of polymer LH, we used an *in vitro* human epithelial cell model (HECM). Monolayers of vaginal epithelial cells (A-431) were infected with *C. albicans*, after addition of different drug dilutions with minimum preincubation time. After 24 h, epithelial damage was assayed by lactate dehydrogenase (LDH) release.

The polymer LH did not prevent damage to the vaginal epithelial cells at any concentration tested (**Supplementary Figure S24**). To check biocompatibility of our polymer, we also incubated A-431 cells with polymer LH alone. At concentrations above 128 µg/mL (4-8 times MIC) we observed more than 50% damage (**Supplementary Figure S24**). In our previous study with red blood cells and murine fibroblasts,^20^ more than 50% metabolic activity was observed in fibroblasts up to an LH concentration of 128 µg/mL. However, even close to these concentrations, the polymer did not reduce the damage by *C. albicans* to the vaginal epithelial cells (**Supplementary Figure S24**).

One possibility for the unexpected failure of LH to prevent damage in the HECM could be the bioavailability. To test this hypothesis, we pre-treated *C. albicans* cells with the polymer for 1 h at concentrations between 16 and 512 µg/mL before infection. Damage by *C. albicans* was strongly reduced with this altered protocol in a concentration-dependent manner up to 256 µg/mL (34% of untreated infection control at 256 µg/mL, **Supplementary Figure S25A**). This supports the hypothesis that poor bioavailability in the HECM reduces the antifungal properties of the polymer. Further supporting this, supernatants taken after co-incubation from uninfected vaginal epithelial cells were unable to inhibit *C. albicans* growth *in vitro*, even when the initial concentration of polymer LH exceeded the MIC (**Supplementary Figure S25B**). This suggests that the presence of human cells reduces the concentration of polymer in the supernatant. In contrast, the common antifungal drugs AmpB, caspofungin, and fluconazole (among others) successfully reduced damage by *C. albicans* to the epithelial cells in the HECM (**Supplementary Table S3**).

Since the polymer LH alone unexpectedly did not inhibit damage by *C. albicans* in the HECM, we studied combinations of LH with established antifungal compounds, first at MIC assay conditions and without human cells (**Supplementary Figure S26** and **Table S4**). Synergy was defined as a minimum two-fold decrease in MIC in the presence of the other drug (fractional inhibitory concentration (FIC) index ≤ 0.5). Antagonistic drug combinations result in an FIC index of at least 4, *i. e.*, a MIC increase of at least two-fold. Indifferent (FIC index = 1), and additive (FIC index between 0.5 and 1) effects were also considered.

The compounds selected for our synergy studies differ in their targets: cell wall (Calcofluor White, Congo Red, caspofungin, nikkomycin Z); cell membrane (AmpB, cetyltrimethylammonium bromide (CTAB), dodecyltrimethylammonium bromide (DTAB), sodium dodecyl sulfate (SDS)); or intracellular processes (fluconazole, cycloheximide, tunicamycin, FK506, geldanamycin), which in turn could again affect cell membrane or wall composition. Seven out of these thirteen antifungal compounds showed a synergistic or strong additive effect (FIC index below 0.6) with LH *in vitro*, and were therefore tested in the HECM. In addition to the LDH release assay, propidium iodide was used to visualise dead epithelial cells after 24 h of infection (Figure 6**, Supplementary Figure S27**).

**Figure 6.**
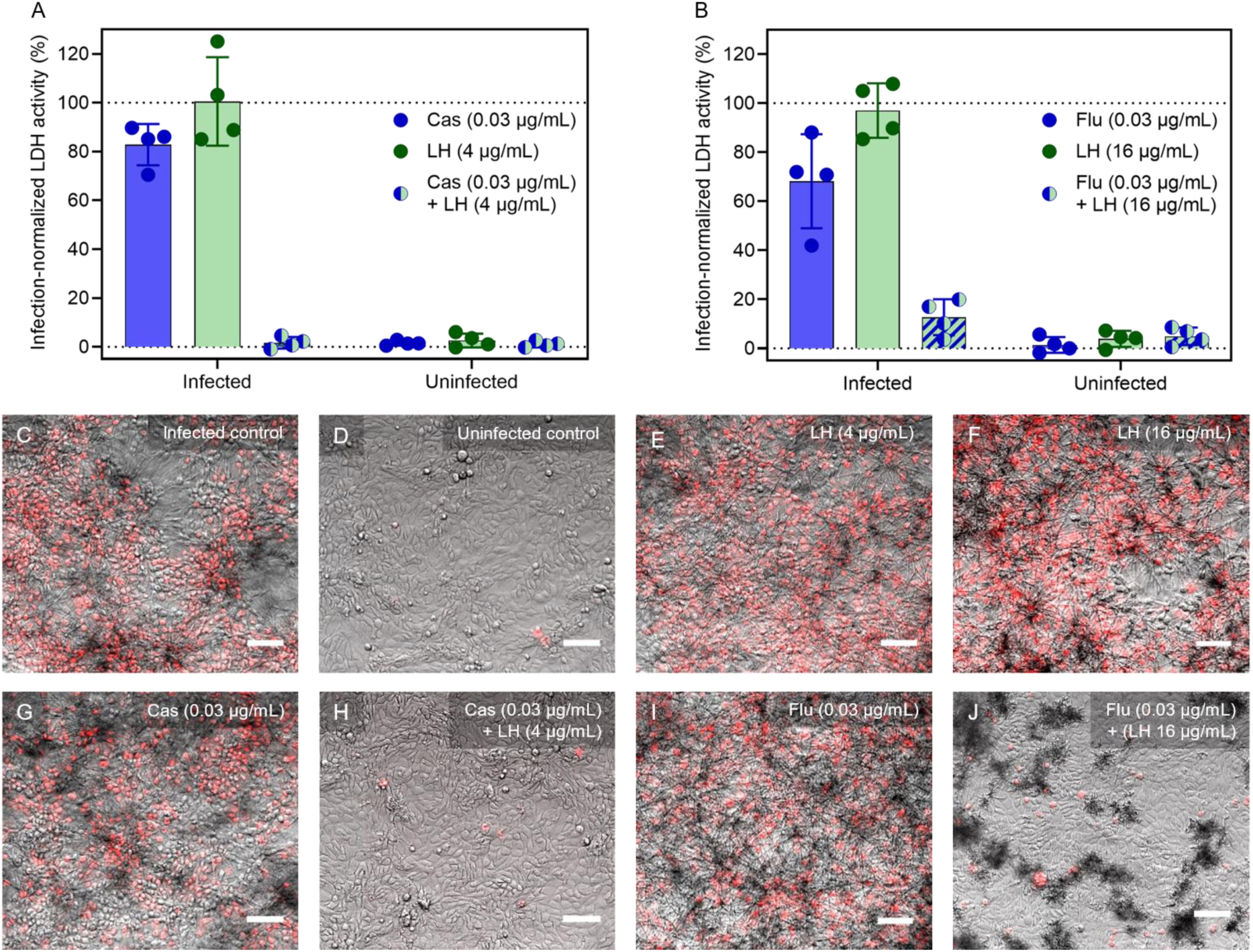
– Synergistic effects of LH in combination with selected antifungal drugs against *C. albicans* SC5314 during vaginal epithelial cell infections. Damage to vaginal epithelial (A-431) cells (infected by *C. albicans* and uninfected) after treatment with polymer LH and (A) caspofungin (Cas) or (B) fluconazole (Flu) and their respective combinations was measured by LDH release. Damage to A-431 cells was normalised to an untreated infection control (for infected samples) or a Triton-X-treated 100% lysis control (uninfected samples). C to J show fluorescence microscopy images of the scenarios represented in A and B to visualise morphological changes and the viability of vaginal epithelial cells by staining with 1 µg/mL propidium iodide. Scale bars represent 100 µm.

Although LH alone did not prevent damage by *C. albicans* (**Supplementary Figure S24**), it did so in combination with the antifungal drugs caspofungin or fluconazole, cells even at normally sub-inhibitory concentrations (Figure 6). Strikingly, only 0.03 µg/mL of caspofungin (*i. e.*, 8× less than its MIC of 0.25 µg/mL in the HECM) combined with 4 µg/mL LH was required to reduce host cell damage to 2% with no visible cytotoxicity (Figure 6A). Similarly, a low dose of 0.03 µg/mL fluconazole (again 8× less than its MIC in the HECM) combined with 16 µg/mL LH reduced vaginal epithelial cell damage to 13%, while remaining biocompatible (Figure 6B). Without LH, these low antifungal drug concentrations did not prevent infection with over 50% epithelial cell damage. This finding was supported by fluorescence microscopy (Figure 6C-J), where the same combinations resulted in healthy vaginal epithelial cells and low doses of antifungal drugs did not significantly protect the epithelial cells from damage at up to 0.25 µg/mL. Combination with LH therefore reduced the MIC for established antifungal drugs up to eight-fold, showing its potential as a synergistic agent for antifungal applications.

The other antifungal compounds with strong additive or synergistic behaviour with LH in the *in vitro* pre-screen (nikkomycin Z, cycloheximide, tunicamycin, FK506, geldanamycin), did not significantly reduce damage to epithelial cells (**Supplementary Figure S27**). Some of those compounds are cytotoxic at elevated concentrations, and indeed, the combination of LH with cycloheximide or geldanamycin caused damage to uninfected human cells.

### Combination of polymer LH and caspofungin prolongs survival in an invertebrate model of *C. albicans* infection

Next, we assessed whether the synergism of polymer LH with caspofungin protected against fungal infection *in vivo*. For this, we used the well-established *Galleria mellonella* (greater wax moth) model of systemic candidiasis by injecting larvae with an infectious dose of *C. albicans* followed by treatment with LH and caspofungin.^59–62^

We first determined the acute toxicity of polymer LH, caspofungin, and AmpB in larvae. Lethal effects of LH to larvae were observed at 500 mg/kg or higher (**Supplementary Figure S28A**). No toxicity was observed for caspofungin at doses up to 100 mg/kg (**Supplementary Figure S28B**, blue line), AmpB showed toxicity at 100 mg/kg after 4 d (**Supplementary Figure S28B**, purple line). Therefore, both LH and caspofungin outperformed AmpB in terms of toxicity, and we determined that doses of LH up to 250 mg/kg and caspofungin up to 100 mg/kg could be used to treat larvae. The synergistic combination of LH with fluconazole was not tested, because it was not active against *C. albicans* in the presence of 50% (v/v) or more of foetal bovine serum in MIC assays (**Supplementary Table S5**).

To simulate systemic candidiasis, we infected each larva with 1×10^5^ *C. albicans* cells in one proleg, treated them in another proleg, and monitored survival over 14 days (Figure 7). All uninfected, water-treated *G. mellonella* larvae survived for 14 d. In contrast, untreated and water-mock treated *C. albicans*-infected larvae died after 2 d (difference not significant at p=0.674).

**Figure 7.**
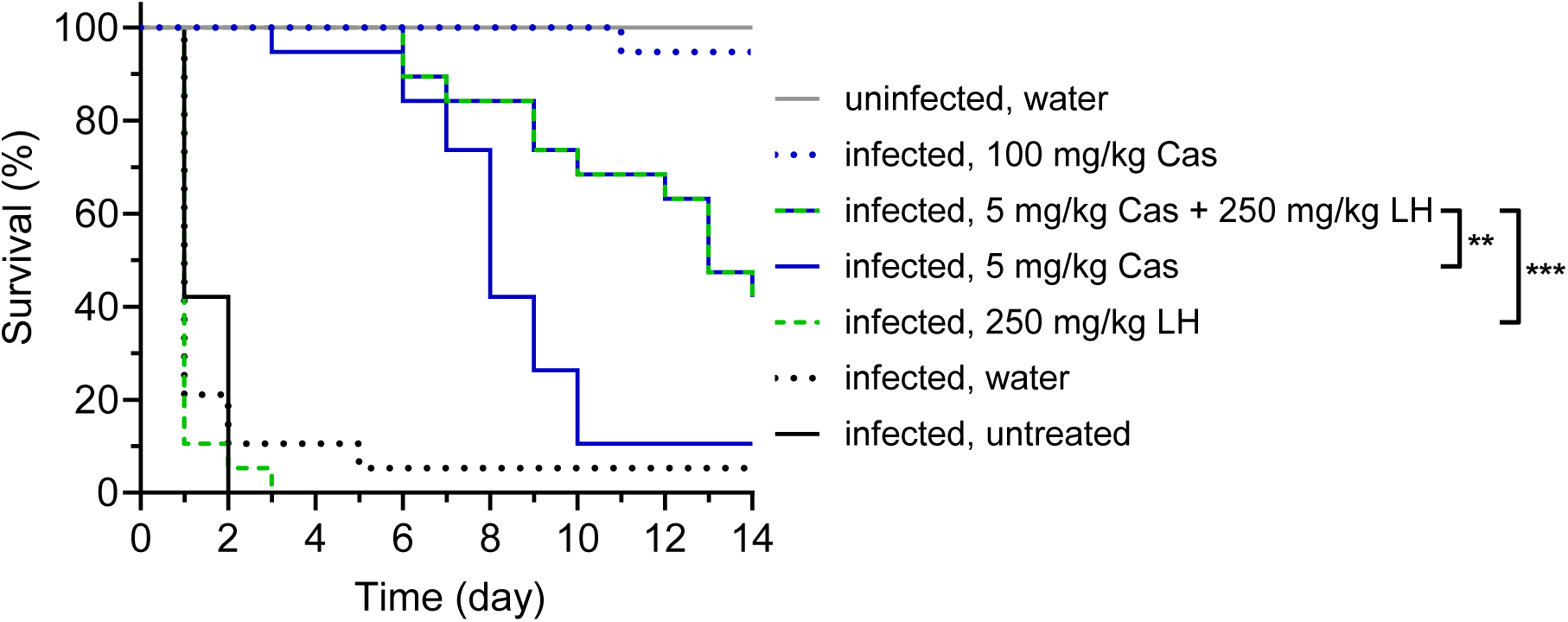
– Treatment with polymer LH in combination with caspofungin (Cas) prolongs survival of *C. albicans*-infected *Galleria mellonella* larvae. *G. mellonella* larvae were infected with 1×10^5^ *C. albicans* cells (except uninfected control, grey) and treated after 2 h with caspofungin (Cas; 5 mg/kg: blue; 100 mg/kg: blue dotted), polymer LH (250 mg/kg, green dashed), and the combination of 5 mg/kg Cas and 250 mg/kg LH (green and blue dashed). Untreated, infection controls are shown in black (*C. albicans* only) or black dashed (*C. albicans*, injected with water). Survival of *G. mellonella* was monitored over 14 d. Nineteen larvae per condition were tested. Statistical significance was determined for infected larvae treated with the combination (5 mg/kg Cas + 250 mg/kg LH) compared to infected larvae treated with the respective dose of single drug by Log Rank (Mantel-Cox) pairwise comparison (** p<0.01, *** p<0.001).

Treatment with 100 mg/kg caspofungin alone protected 94.7% of the larvae from death, whereas only 10.5% of larvae survived to 14 d at 5 mg/kg (Figure 7). All infected larvae treated with polymer LH alone at 250 mg/kg succumbed by day 3, however, the same dose in combination with low-dose caspofungin (5 mg/kg) significantly increased the survival of *C. albicans*-infected larvae (5 mg/kg caspofungin vs. combination: p=0.003; 250 mg/kg LH vs. combination: p<0.001; water vs. combination: p<0.001). Compared to low-dose caspofungin treatment, the survival after 14 d increased four-fold to 42.1%. Notably, 100% of infected larvae treated with the synergistic combination survived until day 6 post-infection. This therefore demonstrates *in vivo* synergy between polymer LH and caspofungin and emphasises the *in vivo* potential of the synthetic polymer LH in combination therapy.

### *In vitro* evolution leads to tolerance of *C. albicans* to LH, but not to the combinations of LH with caspofungin or fluconazole

A major obstacle in the development of antifungal drugs is the emergence of resistance. We used an *in vitro* evolution assay (Figure 8A) to determine whether prolonged exposure to polymer LH, caspofungin, fluconazole or their synergistic combinations results in the emergence of tolerant *C. albicans* variants. Growth at 1×MIC over 14 d (Figure 8B) showed an increasing tolerance for the single drug treatments after 9-10 d incubation. A less pronounced tolerance developed for the combination of fluconazole and LH, while essentially no change in growth was seen for the combination of caspofungin and LH.

**Figure 8.**
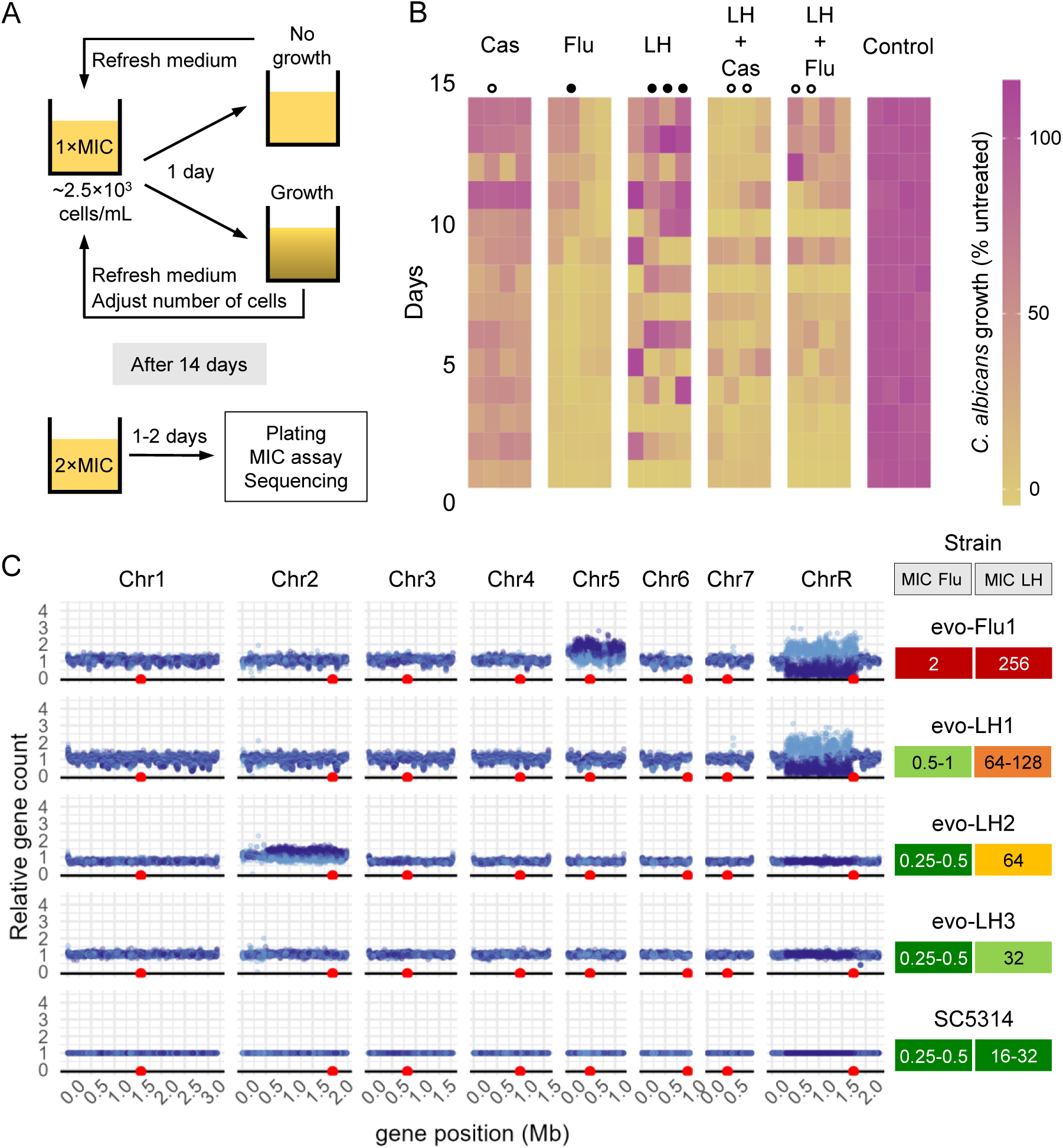
– *In vitro* evolution experiment of *C. albicans* challenged with 1×MIC of caspofungin (Cas), fluconazole (Flu), polymer LH, and the synergistic combinations. (A) Experimental setup of the *in vitro* evolution experiment. (B) Growth of *C. albicans* was monitored by absorbance over 14 d and normalised to the untreated controls. Filled circles highlight strains that were selected for whole genome sequencing and empty circles highlight strains additionally analysed for their MIC against antifungal compounds in **Supplementary Table S6**. (C) Genomes of the isolated strains with the highest tolerance to antifungal drugs were sequenced and analysed for their relative gene count for Flu- and LH-evolved strains. Each point in C represents the relative count compared to wild-type *C. albicans* strain SC5314 at t=0 for a gene (Y-axis) on its chromosome position (X-axis), colour-coded by allele. Positions of the centromeres are indicated by red circles. The MICs for Flu and LH are indicated on the right, where MIC in SC5314 is depicted in green and higher values scale to red.

After 14 d, an aliquot of the cells was incubated at 2×MIC for 24-48 h. If growth was observed, the sample was plated, single colonies were isolated and their MIC to the antifungal polymers (LP, LH, CB, and CX) and antifungal drugs (AmpB, fluconazole, caspofungin, tunicamycin) was determined (**Supplementary Table S6**). Isolates with an increased MIC were selected for further analysis (Figure 8B, filled circles – increased tolerance to LH, empty circles – no increase in LH tolerance). In agreement their growth pattern over 14 d, we did not find any stably tolerant isolates after treatment with combinations of LH and caspofungin or fluconazole (MICs in **Supplementary Table S6**).

To identify the genetic basis for LH tolerance, we sequenced the genomes of three independent LH-evolved strains with high MIC (evo-LH1, evo-LH2, and evo-LH3, filled black circles in Figure 8B), and a fluconazole-evolved strain (evo-Flu1) as control. Evo-LH1 showed an increase in MIC against LH to 64-128 µg/mL, and was also more tolerant to the polymers LP, CB, and CX. The MICs towards established antifungal drugs (AmpB, caspofungin, fluconazole, tunicamycin) remained unchanged. Interestingly, evo-Flu1 not only showed increased tolerance to fluconazole (MIC 2 µg/mL), but also to caspofungin and the polymers. The caspofungin-evolved strain (evo-Cas1) developed an increased MIC for caspofungin (2 µg/mL), but we did not observe any cross-resistance.

To investigate the genomic mechanisms of *C. albicans* adaption to LH or fluconazole, we examined the gene counts for the evolved strains compared to wild-type *C. albicans* SC5314 (Figure 8C, **Supplementary Figure S29**). In all strains, including the parental wild-type SC5314, loss of heterozygosity was detected in the left arm of chromosome 2 (**Supplementary Figure S29**). Analysis of the evolved strains revealed major ploidy changes in some cases: In evo-Flu1, we observed aneuploidy of chromosome R and trisomy of chromosome 5. Both are associated with fluconazole resistance,^63, 64^ and the chromosome 5 trisomy has been shown to be driven by amplification of the *ERG11* and *TAC1* genes.^65^ For evo-LH1 and evo-LH2, we found aneuploidy in chromosome R, and trisomy in chromosome 2, respectively. Notably, chromosome 2 trisomy and tetrasomy have been linked to adaption to the ER stressor tunicamycin.^66^ For the strain evo-LH3, which showed only a minor change in MIC, we found no large-scale ploidy changes.

We did not find mutations or copy number variations on the gene level that could explain the drug tolerance phenotype. We conclude that major chromosomal aberrations probably drive tolerance to polymer LH in *C. albicans*. Together with our transcriptomics and chemical-genetic screening evidence, we therefore postulate that polymer LH has multiple targets, which differ from those of known antifungal drugs.

Importantly, in our *in vitro* evolution experiment we found no development of genetically stable tolerance by combinatorial treatment of LH with caspofungin or fluconazole, in further support of our hypothesis of distinct targets. The strong synergistic action with caspofungin against *C. albicans* on human epithelial cells *in vitro* and *G. mellonella* larvae *in vivo*, together with the reduced tolerance development suggests that the amphiphilic polymer LH is a promising antifungal lead for combination therapy with well-established drugs.

## Conclusions

In this work, we investigated four synthetic polymers, which were inspired by amphiphilic antimicrobial peptides and which kill drug-resistant clinical *C. albicans* isolates. Our findings reveal that the most promising polymer, LH, exerts its activity on *C. albicans* by a putatively novel mode of action. We found evidence that it targets protein glycosylation, and also interferes with the fungal membrane, which together leads to fungal cell death. The combination of LH with caspofungin is particularly promising in its therapeutic potential since it inhibits infection of human epithelial cells by *C. albicans* at otherwise sub-inhibitory concentrations, and additionally increases fungal uptake by human macrophages. Moreover, the synergistic combination of polymer LH and caspofungin prolonged survival in an *in vivo* model of systemic candidiasis. In addition to these promising synergistic effects, which prevent *C. albicans* infection *in vitro* and protect *in vivo*, the combination of polymer LH and caspofungin did not lead to any resistant *C. albicans* strains after prolonged exposure *in vitro*, highlighting the therapeutic potential of polymer LH as an antifungal lead particularly for combination therapy. In light of the promising performance of polymer LH *in vitro* and *in vivo*, further optimisations are conceivable regarding its antifungal activity as a stand-alone-formulation.

## Materials and methods

### Polymer synthesis

#### Materials

Ethylenediamine (Sigma-Aldrich, ≥99%), *N*-amylamine (Sigma-Aldrich, 99%), *N*-benzylamine (Sigma-Aldrich, 99%), *N*-heptylamine (Sigma-Aldrich, 99%), *N*-cyclohexanemethylamine (Sigma-Aldrich, 98%), di-tert-butyl dicarbonate (Sigma-Aldrich, 99%), *N*-hydroxyethyl acrylamide (Sigma-Aldrich, 97%), triethylamine (TEA) (Scharlau, 99%), trifluoroacetic acid (TFA) (Sigma-Aldrich, 99%), chloroform (Merck), dichloromethane (DCM) (Merck), tetrahydrofuran (THF) (Merck), diethyl ether (Merck), hexane (Merck), dimethyl sulfoxide (DMSO) (Merck), dimethylacetamide (DMAc) (Sigma-Aldrich), thionyl chloride (Sigma-Aldrich, 99%), acrylic acid (Sigma-Aldrich), deuterated DMSO (Cambridge Isotope Laboratories, Inc.), 2-(butylthiocarbonothioylthio)propanoic acid (BTPA, Boron Molecular) and 5,10,15,20-tetraphenyl-21*H*,23*H*-porphine zinc (ZnTPP) (Sigma-Aldrich) were used as received.

#### Acryloyl chloride synthesis

Acryloyl chloride was synthesised according to the previously reported procedure,^67^ with slight changes.^20^ Briefly, acrylic acid (41.2 mL, 1.2 equiv) was added dropwise to 36.3 mL of thionyl chloride at 0 °C over 45 min under nitrogen. The mixture was stirred for 12 h at 40 °C. The product was collected by *in situ* distillation under atmospheric pressure.

### Synthesis of monomers

#### Cationic monomer - tert-butyl (2-acrylamidoethyl)carbamate

*tert*-Butyl (2-acrylamidoethyl)carbamate was prepared according to the previously reported procedure.^20, 68, 69^ Ethylenediamine (0.33 mol) was dissolved in chloroform (400 mL). Di-*tert*-butyl dicarbonate (0.03 mol) was dissolved in 100 mL of chloroform and was added dropwise to the ethylenediamine solution over 4 h at 0 °C while stirring and continued over night at room temperature. After filtering the white precipitate, the organic phase was washed with 200 mL of Milli-Q water six times and then dried using MgSO4. Solids were separated by filtration, and chloroform was evaporated, resulting in a pale-yellow oil. THF (100 mL) was added to dissolve the obtained oil. TEA (1.2 equiv) and acryloyl chloride (1.1 equiv) were added dropwise to the solution at 0 °C with N_2_ bubbling. The reaction mixture was stirred at room temperature for 2 h. Afterward, THF was removed by rotary evaporation. The crude product was dissolved in chloroform (150 mL) and washed with 0.1 M HCl solution (1×75 mL), saturated NaHCO3 (1×75 mL), brine (1×75 mL), and water (1×75 mL). The organic phase was dried using MgSO4 and filtered, and the remaining solvent was removed by rotary evaporation. The product was further purified by repeated precipitation steps in hexane to yield the Boc-protected monomer as a fine white powder, which was dried *in vacuo*.

#### Synthesis of hydrophobic monomers

A standard procedure, as previously reported,^20, 69^ was employed for the synthesis of four hydrophobic monomers (*N*-pentylacrylamide, *N-*heptylacrylamide, *N-* cyclohexanemethyl)acrylamide, and *N-*benzylacrylamide) from their corresponding amines (*N*-amylamine, *N*-heptylamine, *N*-cyclohexanemethylamine, or *N*-benzylamine) using acryloyl chloride.

### Random copolymerisation by photoinduced electron/energy transfer-reversible addition-fragmentation chain transfer (PET-RAFT) polymerisation

The linear, random copolymers were synthesised using a slight modification of the general one-pot protocol reported previously.^70^ Briefly, stock solutions of the monomers were prepared with a concentration of 33% (w/w) in DMSO. ZnTPP was dissolved in DMSO at a concentration of 1 mg/mL. The RAFT agent BTPA was added to a 4 mL glass vial in an amount corresponding to the targeted *X*_n_ of 20 and dissolved in DMSO. Monomer stock solutions were added into the vial to a final monomer concentration of 25% (w/w) in DMSO, corresponding to the targeted ratios. The photocatalyst (ZnTPP) was added at 100 ppm relative to the monomers. The vial was sealed with a rubber septum, and the headspace was degassed with N_2_ for 10 min. The vial was then placed under a green light-emitting diode light (*λ* = 530 nm) for 20 h to produce the Boc-protected copolymers. The copolymers were analysed by size-exclusion chromatography (SEC) and ^1^H nuclear magnetic resonance (NMR) to examine the monomer conversion, polymer composition, and molecular weight distribution. Then, the polymer was purified by precipitating in a diethyl ether/hexane mixture (3:7), followed by centrifugation (9000 rpm for 5 min, 0 °C). The precipitate was dissolved in acetone or methanol and reprecipitated twice more. The polymer was then dried *in vacuo* prior to Boc-group removal.

#### Polymer deprotection

TFA was used to remove Boc-protecting groups based on our group’s previously reported protocol.^70^ Briefly, the polymer was dissolved in DCM (∼7% (w/w) polymer), followed by the addition of TFA (20 mol equivalent with respect to Boc groups). The mixture was stirred at room temperature for 3 h and precipitated into diethyl ether. The precipitate was isolated by centrifugation, dissolved in acetone, and reprecipitated twice more. The polymer was then dried *in vacuo*, and ^1^H NMR analysis was used to determine the removal of Boc-protective groups and to examine the targeted *X*_n_.

#### Polymer characterisation

^1^H NMR spectra were obtained using a Bruker AVANCE III spectrometer (300 MHz, 5 mm BBFO probe) or a Bruker AVANCE III 400 spectrometer (400 MHz, 5 mm BBFO probe). Deuterated DMSO was used as a solvent to determine the polymer composition and conversion at concentrations of ∼10–20 mg/mL. All experiments were run with a gas flow across the probes at 535 L/h with sample spinning and at a temperature of 25 °C. All chemical shifts were stated in parts per million (ppm) relative to tetramethylsilane.

SEC analysis was performed using a Shimadzu liquid chromatography system equipped with a Shimadzu refractive index detector and three MIX C columns operating at 50 °C. DMAc (containing 0.03% (w/v) LiBr and 0.05% (w/v) 2,6-dibutyl-4-methylphenol) was used as the eluent at a flow rate of 1 mL/min. The system was calibrated using narrow poly(methyl methacrylate) (PMMA) standards with molecular weights from 200 to 10^6^ g/mol.

### Biological experiments

#### Media and buffers

Phosphate-buffered saline (pH 7.4)

A 10×phosphate-buffered saline (PBS) stock (1.37 mol/L sodium chloride, 0.027 mol/L potassium chloride, 0.08 mol/L disodium hydrogen phosphate, 0.02 mol/L potassium dihydrogen phosphate) was prepared by dissolving 80.06 g sodium chloride, 2.01 g potassium chloride, 11.36 g disodium hydrogen phosphate, and 2.72 g potassium dihydrogen phosphate in 900 mL of double-distilled water and the pH was adjusted to 7.4, before adding double-distilled water up to a final volume of 1 L. The solution was autoclaved for sterilisation. A 1×PBS solution was obtained by dissolving 100 mL of 10×PBS in 900 mL sterile double-distilled water.

Yeast extract peptone dextrose medium

The yeast extract peptone dextrose (YEPD; 1% (w/v) yeast extract, 2% (w/v) mycological peptone, and 2% (w/v) D-glucose) broth was prepared by dissolving 4 g yeast extract and 8 g mycological peptone in double-distilled water up to a total volume of 360 mL. After autoclaving, 40 mL of filter-sterilised 20% (w/v) D-glucose was added. For YEPD plates, 8 g agar was added to the solution before autoclaving.

Synthetic defined medium

Synthetic defined (SD) medium was prepared by dissolving 6.7 g yeast nitrogen base (YNB) without amino acids, and 0.395 g complete supplement mixture were dissolved in 900 mL double-distilled water, adjusted pH to 6.0 with HCl and NaOH, and autoclaved. Afterwards, 100 mL of filter-sterilised 20% (w/v) D-glucose was added. When required, 5 mL of a filter-sterilised 5 mg/mL uridine solution was added, too.

Modified Roswell Park Memorial Institute (RPMI)-1640 medium for *C. albicans* studies To 1 L of RPMI-1640 medium (with L-glutamine, without bicarbonate), 18 g D-glucose and 34.53 g 3-(*N*-morpholino)propane-1-sulfonic acid (MOPS) were added. Afterward, the pH was adjusted with HCl and NaOH to 4.0 and filter-sterilised.

### Polymer and antifungal stock solutions

Polymer stock solutions were prepared at a concentration of 10 mg/mL in sterile distilled water and stored at 4 °C. Before use, the stock solutions were sonicated for approx. 3 min.

Antifungal drug stocks were prepared at different stock solutions in sterile distilled water or DMSO, as summarised in **Table 3**.

**Table 3.** Concentrations, medium and storage temperature of antifungal stocks.

| Antifungal compound | Stock concentration (mg/mL) | Medium | Storage temperature |
| --- | --- | --- | --- |
| Amphotericin B (AmpB) | 1 | DMSO | 4 °C |
| Calcufluor white | 50 | DMSO | -20 °C |
| Caspofungin | 1 | Distilled water | -20 °C |
| Cetyltrimethylammonium bromide (CTAB) | 10 | Distilled water | Room temperature |
| Congo red | 6 | Distilled water | 4 °C |
| Cycloheximide | 5 | Distilled water | 4 °C |
| Dodecyltrimethylammonium bromide (DTAB) | 10 | Distilled water | Room temperature |
| FK506 | 5 | DMSO | -20 °C |
| Fluconazole | 1 | DMSO | 4 °C |
| Geldanamycin | 5 | DMSO | -20 °C |
| Nikkomycin Z | 5 | Distilled water | 4 °C |
| Sodium dodecyl sulphate (SDS) | 10 | Distilled water | Room temperature |
| Tunicamycin | 5 | DMSO | 4 °C |

### Culture conditions of fungal strains

The yeasts were routinely streaked on YEPD agar and incubated for 1-2 days at 30 °C (37 °C for clinical isolates). Cultures on agar plates were stored for up to two weeks at 4 °C. Long-term stocks were stored at -80 °C in 50% (v/v) sterile glycerol from an overnight culture. Overnight cultures were prepared by inoculating colonies from a YEPD plate in YEPD broth and shaking overnight at 30 °C at 180 rpm (37 °C for clinical isolates).

### Fungal strains

A complete list of used fungal strains is attached in the **Supplementary information**. For most assays, the *C. albicans* reference strain SC5314 was used, unless otherwise indicated.

### Minimum inhibitory concentration (MIC) assay and evaluation of drug interactions

The MICs of polymers against different strains of *C. albicans* were determined *via* the broth microdilution method according to Clinical and Laboratory Standards Institute (CLSI) guidelines for fungal susceptibility testing, with slight modifications.^25, 26^ Briefly, the *C. albicans* strains were grown on YEPD plates for 48 h at 30 °C (37 °C for clinical isolates). One colony was emulsified in 1 mL of sterile Milli-Q water. Cells were counted using a haemocytometer and adjusted to 2-5×10^6^ cells/mL. The cell suspension was diluted 1:1000 in the modified RPMI-1640 medium (supplemented with glucose and MOPS, pH 4.0) to obtain the 2×concentrated stock suspension. A two-fold dilution series of the 100 μL polymer solution was added into 96-well microplates (final concentration between 4 and 512 μg/mL), followed by the addition of 100 μL of fungal cell suspension. The 96-well plates were incubated for 24 h at 35 °C in a humidified chamber, wells were resuspended, and the absorbance was measured at 405 nm with a microtiter plate reader. Additionally, AmpB, fluconazole, and caspofungin were tested at final concentrations between 0.125–16 (AmpB) and 0.06–8 μg/mL (fluconazole and caspofungin). DMSO controls at the respectively used final concentrations, no-polymer and no-cell controls were included in all experiments. The MIC value was defined as the lowest concentration of the respective polymer that showed growth inhibition of >90% compared to the untreated control. Three independent biological replicates were carried out (unless otherwise indicated).

To evaluate interactions of the polymer LH with selected antifungal compounds (see **Table 3**), *C. albicans* cells were treated and prepared as described above. The drug-dilution plates were prepared separately for each drug before combining them. For that, a four-fold dilution series of the respective antifungal drug was added into a 96-well microplate along the rows. The same was performed with polymer LH, diluting it in a 96-well microplate along the columns. Then, 50 µL of antifungal were combined with 50 µL of polymer LH, resulting in a two-fold dilution series of the respective compounds. Thereby, one row and one column acted as controls only containing one compound. Drug interactions were classified according to their fractional inhibitory concentration (FIC) indices. The FIC index was calculated as described in the following equation, where c_A/B_ are the concentrations of compounds A or B, respectively, in combination resulting in growth inhibition >90%, and MIC_A_ and MIC_B_ are the MICs of compound A or B, respectively, alone:

A combination was called synergistic if the FIC index was below 0.5, antagonistic for an FIC index above 4 and values in between as additive (between 0.5 and 1.0) or indifferent (between 1 and 4). Three independent biological replicates were carried out.

### RNA isolation, microarray and KEGG pathway analysis

For RNA isolation, *C. albicans* SC5314 was grown in YEPD broth overnight, diluted 1:50 in YEPD broth and subcultured for 4 h (30 °C, 180 rpm). The cells were washed three times in PBS (2,500×g, 1 min) and adjusted to approx. 5×10^7^ cells/mL in SD broth. The cell suspension was added to a final concentration of approx. 5×10^6^ cells/mL into sterile glass flasks containing 6 mL SD medium supplemented with no polymer (for the untreated control) or 32 (LH), 64 (CB and CX), or 128 µg/mL (LP and poly-HEA) polymer. The flasks were incubated for 1 h at 30 °C and 180 rpm. Three biological replicates were performed for each condition. Cell viability was ascertained by backplating on YEPD agar. Fungal cells were harvested for subsequent RNA isolation (2,500×g, 2 min, 4 °C) and handled on ice from that step onwards. RNA was then isolated using a RNeasy mini kit (QIAGEN) by mechanical disruption with acid-washed glass beads following instructions in the manual. Concentration and quality of RNA were checked by Nanodrop ND-1000 (ThermoScientific) and Bioanalyzer 2100 (Agilent). The Quick Amp Gene Expression Labeling Kit (Agilent) was used to synthesise Cy5-labelled cRNA. A common reference (RNA from a mid-log-phase-grown *C. albicans* SC5314 ^71^) was labelled with Cy3 following the same procedure. The dye incorporation was assured by spectrophotometric measurement using a NanoDrop ND-1000. Samples and common reference were cohybridised on Agilent arrays (AMADID 026869) containing 15,744 *Candida albicans* probes corresponding to 6,105 genes. The arrays were then scanned in a GenePix 4000B (Molecular Devices) with GenePix Pro 6.1 (AutoPMT; pixel size of 5 µm) and analysed with GeneSpring 14.8 (Agilent).

Data was log2-transformed and normalised to untreated wild-type SC5314. A heatmap with dendrogram depicting hierarchal clustering (based on Euclidean distance) was generated with base R. PCA and k-means clustering (K=3) was performed and plotted on R using packages stats and factoextra. We performed KEGG pathway enrichment analysis on 1.65× up- or downregulated genes using clusterProfiler package on R ^72, 73^. For the gene set enrichment analysis, the parameters used were: minimum gene set size of 10, adjusted p-value of 0.05. For GO term enrichment analysis on at least two-fold differentially regulated genes, the GO term finder online tool ^31^ on *Candida* Genome Database was used and assessed on 18/01/2023 with the following parameters: species and background set – *Candida albicans*; ontology – molecular function; adjusted maximum p-value of 0.1. The microarray data are available in the ArrayExpress database (https://www.ebi.ac.uk/biostudies/arrayexpress) under accession number E-MTAB-13294.

### Mutant screening

A complete list of the tested mutants and reference strains is shown in the **supplementary information**. The experiments were generally performed in SD broth, supplemented with uridine if necessary.

Yeast cultures of the mutants were grown overnight at 30 °C, 180 rpm. The cultures were washed twice in PBS (1 min, 5,000×g). After counting with a haemocytometer, cell concentration was adjusted to approx. 10^6^ cells/mL in SD broth and diluted 1:10. In a 96 well plate, 100 µL of yeast suspension was mixed with 100 µL of SD broth (untreated control) or SD broth supplemented with 2× concentrated antifungal polymers to reach final concentrations of 16 (LH), 32 (CB and CX), or 64 µg/mL (LP). Reference strains relevant for the tested mutants were incubated with and without treatment alongside. All samples were prepared in technical duplicates and performed independently in triplicates. The 96-well plate was covered with sterile sealing foil. Growth curves were recorded with the infinite 200 or infinite 200Pro microplate reader (Tecan, i-control software) over 3 days at 30 °C by measuring absorbance at 600 nm every 15 min after orbital shaking for 10 s. To compare the growth curves of various mutants to their reference strains, a growth speed index based on time until half-maximum growth was used. The growth speed index was calculated as follows:

Here, M is time until half-maximum growth was reached for mutants, treated or untreated, and equally WT corresponds to the respective wild-type or reference strain. Thus, negative values display reduced growth speed in the presence of polymer and positive values reflect beneficial growth of the mutant compared to the reference strain in the presence of polymer.

Two- or four-fold increase of antifungal polymers consistently led to no growth of the yeast. In case of unexpected fast growth or late-onset growth at those elevated concentrations, the experiment was excluded and repeated.

### Cell lysis assay *via* fluorescence microscopy

A *C. albicans* SC5314 mutant, expressing GFP intracellularly under the regulation of the constitutive *ADH1* promoter (*adh1*::P*_ADH1_*-GFP CaSAT1) was grown overnight in YEPD broth. The cells were subcultured 1:50 in YEPD broth for 4 h (30 °C, 180 rpm), washed three times in PBS (2,500×g, 1 min) and diluted to 2×10^5^/mL in modified RPMI (with glucose and MOPS, pH 4.0). Of this, 100 µL were added to 100 µL of 2× concentrated antifungal stock solutions in modified RPMI in technical duplicates in a 96-well plate to final concentrations of 16, 32, and 64 µg/mL polymer LH (0.5×, 1×, and 2×MIC), 2 µg/mL AmpB (1×MIC), or 8 µg/mL tunicamycin (1×MIC) with a medium-only control. Inoculation counts were checked by backplating on YEPD. After 6 h incubation at 35 °C in a humidified chamber, samples from each well were backplated in duplicates on YEPD to check viability compared to the inoculum control and final cell counts. The 96-well plate was then centrifuged at 250×g for 10 min, the supernatant removed carefully, and cells were fixed in 75 µL ROTI®Histofix (Carl Roth, Germany) for 30 min at room temperature. The plates were centrifuged again, supernatants discarded and 200 µL PBS added for microscopical analysis. A ZEISS Celldiscoverer 7 (20× plan-apochromat objective, 2×magnification, equipped with an Axiocam 506) was used for acquiring brightfield images (5-20 ms exposure) and detection of fluorescence (LED excitation at 470 nm for 500 ms, 501-547 nm emission filter). The ZEN software (blue edition, ZEISS) was used to normalise fluorescence of treated samples to untreated control. Fluorescence intensities were quantified by region-of-interest measurements in ImageJ^74^ of >50 *C. albicans* cells relative to untreated cells (100%). The experiment was repeated three times independently.

### Transmission electron microscopy of *C. albicans*’ cell wall

For transmission electron microscopy (TEM), a *C. albicans* overnight culture (YEPD broth, 30 °C, 180 rpm) was diluted 1:50 in YEPD and incubated for 4 h at 30 °C and 180 rpm. After centrifugation (2,000×g, 1 min), the pellet was washed in PBS and the cell concentration was adjusted to 10^7^/mL in SD broth. Of this, 200 µL were added to 9.8 mL SD broth (untreated control), SD broth plus LH (128 µg/mL final concentration), SD broth plus tunicamycin (4 µg/mL), or SD broth plus poly-HEA (128 µg/mL) and incubated in glass flasks for 6 h at 180 rpm and 30 °C. Cell viability was checked by backplating on YEPD. The samples were centrifuged for 5 min at 4 °C and 1,000×g, the cell pellets were kept on ice and inactivated by ultraviolet light using a Vilber-Lourmat BLX312 UV crosslinker. For TEM preparation, samples were mixed with 20% (w/v) BSA and high-pressure-frozen with a HPM 010 (BAL-TEC) in 200 µm HPF carriers (Baltic Präparation). Freeze substitution (FS) was performed in acetone containing 1% (w/v) osmium tetroxide and 0.1% (w/v) uranyl acetate (12 h at -90 °C, 8 h at -60 °C, 8 h at -30 °C) using the FSU 010 (BAL-TEC). After FS, samples were infiltrated and embedded in Lowicryl HM20 (Polysciences) at -4 °C. Polymerisation was carried out with UV light. For ultrathin sectioning (60 nm), an ultramicrotome (Leica Ultracut E; Leica Biosystems) was used. After mounting on filmed copper grids and post-staining with lead citrate, the sections were studied in a transmission electron microscope (EM 902 A; ZEISS) at 80 kV. Images were acquired with a 1k FastScan CCD camera (TVIPS).

### Hypha formation assay

To study formation of hyphae and microcolonies by *C. albicans*, the cell concentration of a PBS washed overnight culture was adjusted to 6×10^4^/mL (4 and 6 h timepoints) and 400/mL (24 h timepoint) in RPMI-1640 (with L-glutamine, without phenol red). In a 24-well plate, 500 µL of RPMI-1640 medium plus LP (64 µg/mL final concentration), LH (16 µg/mL for 4 and 6 h timepoint, 4 µg/mL for 24 h timepoint), CB (32 µg/mL), or CX (32 µg/mL) were prepared. To this, 500 µL of diluted *C. albicans* suspension was added and the plates were incubated for 2, 4, or 24 h at 37 °C with 5% CO_2_ in a humidified chamber to induce formation of hyphae and microcolonies. After 2, 4, or 24 h, cells were fixed with formaldehyde (approx. 10% (v/v) final concentration from a 37% stock). The plates were then stored until imaging at 4 °C. Fixed cells were imaged using a ZEISS Axio Vert A1 microscope, equipped with a plan-apochromat 10× objective, AxioCam ICc1, and a 0.63× camera adapter. The internal ZEN blue software was used to measure the length of 50 hyphae and diameters of 50 microcolonies per sample. The experiment was performed independently in triplicates.

### Fungal clearance by macrophages

Human peripheral blood mononuclear cells (PBMCs) and monocytes were isolated from buffycoats donated by healthy volunteers after obtaining written informed consent. The donation procedure was approved by the Jena institutional ethics committee (Ethik-Kommission des Universitätsklinikums Jena, Permission No 2207–01/08). Isolation of PBMCs was performed by density gradient centrifugation over Lymphocytes Separation Medium (Capricorn Scientific) in a sterile 50 mL tube. CD14+ monocytes were selected from the PBMC fraction using magnetic beads and automated cell sorting (autoMACs; MiltenyiBiotec). To differentiate monocytes into monocyte-derived macrophages (MDMs), 1×10^7^ cells were seeded into 175 cm^2^ cell culture flasks in RPMI-1640 media with 2 mM L-glutamine (Thermo Fisher Scientific) containing 10% heat-inactivated foetal bovine serum (FBS; Bio&SELL) and 50 ng/mL recombinant human macrophage colony-stimulating factor (ImmunoTools) and incubated for seven days at 37 °C with 5% CO_2_. Adherent MDMs were detached with 50 mM EDTA in PBS, seeded in 96-well plates at a concentration of 4×10^4^ cells/well and incubated for 24 h in RPMI-1640 at 37 °C with 5% CO_2_. Drug dilution plates (120 µL per well) were prepared on the day of the experiment at two-times final concentration of polymer LH, caspofungin, fluconazole and their respective combinations in RPMI-1640. An overnight culture of *C. albicans ADH1*-GFP (derived from SC5314), expressing GFP intracellularly, was centrifuged at 2,500×g for 1 min and the pellet was washed in PBS twice. GFP-expressing *C. albicans* has been chosen for this assay to facilitate distinction of yeast and macrophages. The washed *C. albicans* cell culture was adjusted to a concentration of 1.2×10^6^ cells/mL in RPMI-1640. 120 µL of the adjusted and washed cell suspension was added to each well of the pre-warmed drug dilution plate and incubated for 1 h at 30 °C in a humidified chamber. Samples for each donor were prepared in duplicates. In addition to the infected samples, uninfected samples were prepared in the same manner by replacing the *C. albicans* solution with RPMI-1640 medium. Afterward, the supernatant of the 96-well plates harbouring the macrophages was aspirated, and 200 µL of the pre-treated *C. albicans* cells in RPMI-1640, supplemented with the respective antifungal compounds, was added to the macrophages. The 96-well plates were centrifuged for 5 min at 200×g to settle the cells for subsequent imaging. Following centrifugation, the 96-well plate was immediately transferred into a ZEISS Celldiscoverer 7 and incubated for 45 min at 37 °C and 5% CO_2_. Images were acquired every 15 min using a 20× plan-apochromat objective at 2×magnification and fluorescence signal of GFP-expressing *C. albicans* was detected (LED excitation at 470 nm for 500 ms, 501-547 nm emission filter). ZEN software (version 3.1, blue edition) was used to analyse the images. Non-phagocytosed cells were counted at time 0 (set to 100%) and timepoint 15 min or 30 min (displayed relative to the count at time 0). The analysed timepoint was chosen based on when in the untreated controls non-phagocytosed *C. albicans* cells decreased to 30-50%, compared to time 0. A total of six donors were investigated on three different days.

### Response of human peripheral blood mononuclear cells (PBMCs)

Human PBMCs were isolated from fresh human blood as described above and seeded in 96-well plates at 1×10^6^ cells/well in 100 µL. The drug dilutions were prepared in RPMI-1640 medium at two-fold final concentration (150 µL each well for 2 donors) in a separate 96-well plate and prewarmed at 30 °C. A washed *C. albicans* overnight culture was diluted to 10^7^ cells/mL in RPMI-1640. 150 µL of the adjusted *C. albicans* suspension was added to the wells of the drug dilution plate and incubated for 1 h. In parallel, uninfected samples were prepared in the same manner by replacing *C. albicans* in RPMI-1640 by medium only to check putative responses of the PBMCs to the antifungal compounds only. The seeded PBMCs were stimulated with 100 µL of the LH and RPMI pre-treated *C. albicans* samples and incubated for 24 h at 37 °C with 5% CO_2_ in a humidified chamber. Afterwards, the 96-well plates were centrifuged at 250×g for 10 min and 150 µL supernatant was collected and stored at -20 °C. Cytokine levels were detected by enzyme-linked immunosorbent assays (ELISA, R&D Systems) and were performed according to the protocol supplied by the manufacturer. Cytokine release was displayed relative to the untreated *C. albicans* control (set to 1) as fold changes. A total of 11 (IL-6 and IL-1β) or 8 donors (TNF) were analysed on 3 or 4 different days.

### *In vitro* human epithelial cell infection model

To mimic human epithelial cells, A-431 epithelial cells (ACC 91)^75^ were used. A-431 cells were derived from a vulva epidermoid carcinoma and are routinely used to simulate the vaginal mucosa.^75–78^ The cells were cultivated in RPMI-1640 medium supplemented with 10% FBS in a humidified incubator at 37 °C with 5% CO_2_ and regularly tested for *Mycoplasma* contamination. For seeding in 96-well plates, 200 µL of a 10^5^ cells/mL suspension was added per well in RPMI-1640 medium, supplemented with 10% FBS. The plates were incubated for 48 h at 37 °C with 5% CO_2_ in a humidified chamber to let the cells attach to the surface. On the day of infection, a washed *C. albicans* SC5314 overnight culture was diluted to 3.64×10^5^ cells/mL in prewarmed RPMI-1640 medium (without FBS). A drug-dilution plate containing 110 µL 2× concentrated antifungal compound(s) was prepared as described in the MIC assay and prewarmed. The supernatants were aspirated from the seeded vaginal epithelial cells and 100 µL of the antifungal-*C. albicans* solutions in RPMI-1640 and 100 µL of the antifungal drug in RPMI-1640 were added with minimum preincubation time to infect the vaginal epithelial cells. As a control, an uninfected plate was prepared as described above without the addition of *C. albicans* (replaced by 100 µL of RPMI-1640). Infected and uninfected samples were incubated for 24 h at 37 °C and 5% CO_2_ in a humidified chamber. Cells were stained with propidium iodide (1 µg/mL final concentration, 1 mg/mL stock) for 15 min at 37 °C with 5% CO_2_ to detect damaged vaginal epithelial cells. The stained samples were imaged using a ZEISS Celldiscoverer 7 (5× plan-apochromat objective, 2×magnification, equipped with an Axiocam 506, prewarmed to 37 °C with 5% CO_2_) for brightfield image acquisition (automatic exposure time and focus) and detection of fluorescence (LED excitation at 545 nm, 583-601 nm emission filter). To quantify damage caused by *C. albicans* to the vaginal epithelial cells, lactate dehydrogenase (LDH), released by the epithelial cells, was measured in the supernatant. For that, Triton X-100 was added to at least two wells containing vaginal epithelial cells in medium at a final concentration of 0.1% from a 5-10% stock and incubated for 5 min at 37 °C with 5% CO_2_, reflecting 100% damage to uninfected epithelial cells. Plates were centrifuged at 250×g for 10 min, and 20 µL from the supernatants were taken up in 80 µL PBS and an LDH detection assay was performed using the Cytotoxicity Detection Kit (LDH) (Roche) following the manual’s instructions. LDH release data is presented relative to the respective controls, *i. e.*, *C. albicans* infected vaginal epithelial cells served as a control for the infected samples (set to 100%), and uninfected vaginal epithelial cells, treated with Triton-X-100, were set to 100% for all uninfected samples. Uninfected, untreated vaginal epithelial cells were used as background controls and were subtracted from all measured values. All experiments were carried out independently at least in triplicates.

### G. mellonella in vivo toxicity assay

The acute toxicity of AmpB, caspofungin, and polymer LH was tested *in vivo* in a *G. mellonella* model using previously described methods with slight modifications.^62, 79^ Briefly, *G. mellonella* larvae were originally obtained from Westmead hospital (Sydney, Australia) and maintained in an environmentally controlled room at Macquarie University (Sydney, Australia) at 30 °C and 65% humidity with a 12 h light/dark cycle. Larvae (200-250 mg) were individually injected with 10 µL of the respective compound into the last right proleg using a 100 µL syringe (Hamilton Ltd.), reaching final concentrations of 100 mg/kg (AmpB, caspofungin) or 250-500 mg/kg (polymer LH). For each condition, four larvae were treated, including a 1% (v/v) DMSO control (as the AmpB stock used DMSO as a solvent) or water (solvent for all dilutions of the compounds). Following injection, the larvae were incubated at 37 °C and monitored every 24 h for 10 days. Larval performance was assessed according to the *G. mellonella* Health Index Scoring System.^80^

### G. mellonella-C. albicans infection model

Before infection, *C. albicans* SC5314 was cultured on YEPD agar plates and grown overnight in YEPD broth at 30 °C and 200 rpm. An aliquot of the culture was then washed twice in PBS and counted using a haemocytometer. Fungal cell viability was ascertained by backplating.

To test the *in vivo* efficacy of caspofungin, polymer LH and the combination of both to prevent infection by *C. albicans*, *Galleria* larvae (200-250 mg) were infected with 1×10^5^ *C. albicans* cells/larva suspended in 10 µL of water in the last right proleg and incubated at 37 °C. After 2 h, 10 µL of the tested compounds or compound combinations were injected in the last left proleg. Three groups of larvae were included as controls: infected larvae treated with 100 mg/kg caspofungin as a positive treatment control, infected larvae injected with water only as a negative treatment control, and uninfected larvae injected with water to control for injection injury. Larval survival was monitored daily over 14 days. The experiment was performed in three biological replicates, whereby each condition was tested in 6-7 larvae per replicate (n = 19 total). Statistical analyses were performed using IBM SPSS Statistics 29.0.

### Evolution assay

For the *in vitro* evolution assay, *C. albicans* SC5314 cells were incubated and prepared as in the MIC assays in the presence of polymer LH (32 µg/mL), the synergistic combinations of LH (4 µg/mL) with caspofungin (0.07 µg/mL) or fluconazole (0.13 µg/mL), and the antifungal drugs fluconazole (0.5 µg/mL) and caspofungin (0.5 µg/mL) alone as controls. All samples, including an untreated control, were prepared in four replicates in a 96-well plate. After 24 h incubation at 35 °C, fungal growth was measured photometrically (absorbance measurement at 405 nm with a Tecan infinite M200 plate reader) and compared to the untreated control. After the first day, growth was inhibited in all samples except the untreated control, resulting in an absorption <0.1, and continued cell viability was assured *via* backplating on YEPD plates. In case of a measured increase in absorbance indicating fungal growth after day 1, the samples were diluted based on their absorption: absorption >0.5 (indicating >5×10^7^ cells/mL) samples were diluted 1:1000; absorption between 0.2 and 0.5 (>10^7^ cells/mL) samples were diluted 1:500; absorption between 0.1 and 0.2 (>10^6^ cells/mL) samples were diluted 1:100. Absorptions below 0.1 were defined as “no visual growth” and further used undiluted. A sample of 20 µL of diluted or undiluted cell cultures were added to 180 µL of fresh medium, plus respective antifungal. This procedure was repeated for 14 consecutive days.

Afterwards, 50 µL were incubated in 450 µL modified RPMI medium containing the antifungal at a concentration of 2×MIC in Eppendorf tubes under gentle rolling at 35 °C. If growth was observed after 24 h or 48 h, a sample was plated on YEPD and 500 µL of 50% (v/v) glycerol was added to the more tolerant fungal cells and frozen at -80 °C. If no growth was observed after 48 h, the sample was considered not more tolerant to the respective antifungal. A *C. albicans* MIC assay for the polymers LP, LH, CB and CX, and the antifungal compounds AmpB, fluconazole, caspofungin and tunicamycin, was performed with the more tolerant strains.

The most tolerant strains were selected for whole genome sequencing. Genomic DNA was isolated from overnight cultures of evolved strains by phenol/chloroform extraction as described in ^81^. Whole genome sequencing of the evolved strains was performed on an Illumina Novaseq, 2×150 bp sequencing and 10 million raw paired-end reads per sample by Azenta life sciences. The sequences were trimmed with Trimmomatic and aligned to *C. albicans* assembly 22 using Bowtie2 (v2.4) with standard settings. The lab strain (SC5314) used for the evolution experiment was also sequenced and assembled. Genomic variants unique to an evolved strain were called with SnpEff.^82^ Changes in ploidy, loss of heterozygosity and copy number variation was determined by YMAP^83^, and also visualised by R. Integrative genomics viewer^84^ and R (package Gviz^85^) were used to visualise genome coverage for genomic and copy number variants. The whole genome sequencing data are available in the European Nucleotide Archive database (https://www.ebi.ac.uk/ena/browser/home) under accession number PRJEB65085.

## Supporting information

Supplementary Table - Mutant strains

## Acknowledgement

SB is supported by the Priority Programm SPP2225 “Exit strategies of intracellular pathogens” of the German Research Foundation (Deutsche Forschungsgemeinschaft – DFG, project 446404928). RV has received funding from the AReST grant of the German Bundesministerium für Bildung und Forschung (BMBF, project 16GW0238K). SA was supported by funding from the European Union’s Horizon 2020 research and innovation program under grant agreement No 847507 (HDM-FUN). MSG is supported by the Deutsche Forschungsgemeinschaft (DFG) Emmy Noether Program (project no. 434385622 / GR 5617/1-1). JLS and MSG were supported by the German Research Foundation (Deutsche Forschungsgemeinschaft, DFG) within the Collaborative Research Centre (CRC)/Transregio (TRR) 124 “FungiNet” project C1 (DFG project number 210879364). ES would like to thank Electron Microscopy Centre in Jena for the support by providing the equipment for TEM preparation and evaluation. BQ would like to acknowledge the German Research Foundation (Deutsche Forschungsgemeinschaft – DFG, project QU116/9-1 for funding. AKC was supported by ARC Future Fellowship FT220100152. AKC and HD acknowledge Macquarie University Research Infrastructure Scheme (MQRIS) grants for funding the *Galleria* Research Facility. PRJ acknowledges the receipt of an Australian Government Research Training Program Scholarship. SS is supported by a University International Postgraduate Award from the University of New South Wales (UNSW). CB would like to acknowledge ARC Laureate Fellowship (FL220100016) for funding.

The authors would like to thank the NMR facility within the Mark Wainwright Analytical Centre (MWAC) at UNSW for providing and maintenance of the necessary instruments; Eh Hau Pan for technical assistance at the School of Chemical Engineering at UNSW; Bernhard Hube (HKI Jena) for supporting SS’s research stay at HKI Jena and his valuable feedback on the project; Maximilian Himmel, Julia Mantke, and Stephanie Wisgott for technical assistance at the HKI Jena; Steffi Feller for assistance with organising SS’s research stay at the HKI in Jena; Osama Elshafee (HKI Jena) for providing the *ADH1*-GFP *C. albicans* strain.

## References

1. Clark, C. & Drummond, R.A. The hidden cost of modern medical interventions: How medical advances have shaped the prevalence of human fungal disease. Pathogens 8, 45 (2019).

2. Perfect, J.R., Hachem, R. & Wingard, J.R. Update on epidemiology of and preventive strategies for invasive fungal infections in cancer patients. Clinical Infectious Diseases 59, S352–S355 (2014).

3. Brown, G.D. et al. Hidden killers: Human fungal infections. Science Translational Medicine 4, 165rv113–165rv113 (2012).

4. Hoenigl, M. et al. COVID-19-associated fungal infections. Nature Microbiology 7, 1127–1140 (2022).

5. Mazi, P.B. et al. Attributable mortality of *Candida* bloodstream infections in the modern era: A propensity score analysis Clinical Infectious Diseases 75, 1031–1036 (2022).

6. Pfaller, M.A., Diekema, D.J., Turnidge, J.D., Castanheira, M. & Jones, R.N. Twenty years of the SENTRY antifungal surveillance program: Results for *Candida* species from 1997-2016. Open Forum Infectious Diseases 6, S79–S94 (2019).

7. Nnadi, N.E. & Carter, D.A. Climate change and the emergence of fungal pathogens. PLOS Pathogens 17, e1009503 (2021).

8. Casadevall, A. Climate change brings the specter of new infectious diseases. The Journal of Clinical Investigation 130, 553–555 (2020).

9. WHO Fungal priority pathogens list to guide research, development and public health action. World Health Organization (WHO) (2022).

10. Perfect, J.R. The antifungal pipeline: a reality check. Nature Reviews Drug Discovery 16, 603–616 (2017).

11. Revie, N.M., Iyer, K.R., Robbins, N. & Cowen, L.E. Antifungal drug resistance: Evolution, mechanisms and impact. Current Opinion in Microbiology 45, 70–76 (2018).

12. CDC Antibiotic resistance threats in the United States, 2019. U. S. Department of Health and Human Services, CDC (2019).

13. Gintjee, T.J., Donnelley, M.A. & Thompson, G.R. Aspiring antifungals: Review of current antifungal pipeline developments. Journal of Fungi 6, 28 (2020).

14. Hoenigl, M. et al. The antifungal pipeline: Fosmanogepix, ibrexafungerp, olorofim, opelconazole, and rezafungin. Drugs 81, 1703–1729 (2021).

15. Pappas, P.G. et al. Clinical safety and efficacy of novel antifungal, fosmanogepix, for the treatment of candidaemia: Results from a Phase 2 trial. Journal of Antimicrobial Chemotherapy, dkad256 (2023).

16. Spitzer, M., Robbins, N. & Wright, G.D. Combinatorial strategies for combating invasive fungal infections. Virulence 8, 169–185 (2017).

17. Revie, N.M. et al. Targeting fungal membrane homeostasis with imidazopyrazoindoles impairs azole resistance and biofilm formation. Nature Communications 13, 3634 (2022).

18. dos Reis, T.F., et al. A host defense peptide mimetic, brilacidin, potentiates caspofungin antifungal activity against human pathogenic fungi. Nature Communications 14, 2052 (2023).

19. Fernández de Ullivarri, M., Arbulu, S., Garcia-Gutierrez, E. & Cotter, P.D. Antifungal peptides as therapeutic agents. Frontiers in Cellular and Infection Microbiology 10 (2020).

20. Schaefer, S. et al. Rational design of an antifungal polyacrylamide library with reduced host-cell toxicity. ACS Applied Materials & Interfaces 13, 27430–27444 (2021).

21. Liu, R. et al. Nylon-3 polymers with selective antifungal activity. Journal of the American Chemical Society 135, 5270–5273 (2013).

22. Jiang, W. et al. Short guanidinium-functionalized poly(2-oxazoline)s displaying potent therapeutic efficacy on drug-resistant fungal infections. Angewandte Chemie International Edition 61, e202200778 (2022).

23. Ng, V.W.L. et al. Antimicrobial polycarbonates: Investigating the impact of nitrogen-containing heterocycles as quaternizing agents. Macromolecules 47, 1285–1291 (2014).

24. Corrigan, N. et al. Reversible-deactivation radical polymerization (Controlled/living radical polymerization): From discovery to materials design and applications. Progress in Polymer Science 111, 101311 (2020).

25. CLSI Performance standards for antifungal susceptibility testing of yeasts, Edn. 1st. (Clinical and Laboratory Standards Institute, Wayne, PA; 2017).

26. CLSI Reference method for broth dilution antifungal susceptibility testing of yeasts, Edn. 4th. (Clinical and Laboratory Standards Institute, Wayne, PA; 2017).

27. Lackner, M. et al. Positions and numbers of *FKS* mutations in *Candida albicans* selectively influence *in vitro* and *in vivo* susceptibilities to echinocandin treatment. Antimicrobial Agents and Chemotherapy 58, 3626–3635 (2014).

28. Martel, C.M. et al. A clinical isolate of *Candida albicans* with mutations in *ERG11* (encoding sterol 14α-demethylase) and *ERG5* (encoding C22 desaturase) is cross resistant to azoles and amphotericin B. Antimicrobial Agents and Chemotherapy 54, 3578–3583 (2010).

29. Martel, C.M. et al. Identification and characterization of four azole-resistant *erg3* mutants of *Candida albicans*. Antimicrobial Agents and Chemotherapy 54, 4527–4533 (2010).

30. Fischer, D. et al. Disruption of membrane integrity by the bacterium-derived antifungal jagaricin. Antimicrobial Agents and Chemotherapy 63, e00707–00719 (2019).

31. Boyle, E.I. et al. GO::TermFinder—open source software for accessing Gene Ontology information and finding significantly enriched Gene Ontology terms associated with a list of genes. Bioinformatics 20, 3710–3715 (2004).

32. Kanehisa, M. & Goto, S. KEGG: Kyoto Encyclopedia of Genes and Genomes. Nucleic Acids Research 28, 27–30 (2000).

33. Kanehisa, M., Furumichi, M., Sato, Y., Kawashima, M. & Ishiguro-Watanabe, M. KEGG for taxonomy-based analysis of pathways and genomes. Nucleic Acids Research 51, D587–D592 (2022).

34. Schaeffer, H.J. & Weber, M.J. Mitogen-activated protein kinases: Specific messages from ubiquitous messengers. Molecular and Cellular Biology 19, 2435–2444 (1999).

35. Monge, R.A., Román, E., Nombela, C. & Pla, J. The MAP kinase signal transduction network in *Candida albicans*. Microbiology 152, 905–912 (2006).

36. Liu, T.T. et al. Genome-wide expression profiling of the response to azole, polyene, echinocandin, and pyrimidine antifungal agents in *Candida albicans*. Antimicrobial Agents and Chemotherapy 49, 2226–2236 (2005).

37. Rautenbach, M., Troskie, A.M. & Vosloo, J.A. Antifungal peptides: To be or not to be membrane active. Biochimie 130, 132–145 (2016).

38. Wang, T. et al. Transcriptional responses of *Candida albicans* to antimicrobial peptide MAF-1A. Frontiers in Microbiology 8 (2017).

39. Sircaik, S. et al. The protein kinase Ire1 impacts pathogenicity of *Candida albicans* by regulating homeostatic adaptation to endoplasmic reticulum stress. Cellular Microbiology 23, e13307 (2021).

40. Badrane, H. et al. The *Candida albicans* phosphatase Inp51p interacts with the EH domain protein Irs4p, regulates phosphatidylinositol-4,5-bisphosphate levels and influences hyphal formation, the cell integrity pathway and virulence. Microbiology 154, 3296–3308 (2008).

41. Hall, R.A. & Gow, N.A.R. Mannosylation in *Candida albicans*: Role in cell wall function and immune recognition. Molecular Microbiology 90, 1147–1161 (2013).

42. Rispail, N. et al. Comparative genomics of MAP kinase and calcium–calcineurin signalling components in plant and human pathogenic fungi. Fungal Genetics and Biology 46, 287–298 (2009).

43. Mayer, F.L. et al. The novel *Candida albicans* transporter Dur31 is a multi-stage pathogenicity factor. PLOS Pathogens 8, e1002592 (2012).

44. Kumar, R. et al. Histatin 5 uptake by *Candida albicans* utilizes polyamine transporters Dur3 and Dur31 proteins. Journal of Biological Chemistry 286, 43748–43758 (2011).

45. Cesare, G.B.D., Cristy, S.A., Garsin, D.A., Lorenz, M.C. & Alspaugh, J.A. Antimicrobial peptides: A new frontier in antifungal therapy. mBio 11, e02123–02120 (2020).

46. Gray, K.C. et al. Amphotericin primarily kills yeast by simply binding ergosterol. Proceedings of the National Academy of Sciences 109, 2234–2239 (2012).

47. Anderson, T.M. et al. Amphotericin forms an extramembranous and fungicidal sterol sponge. Nature Chemical Biology 10, 400–406 (2014).

48. Kuo, S.C. & Lampen, J.O. Tunicamycin — An inhibitor of yeast glycoprotein synthesis. Biochemical and Biophysical Research Communications 58, 287–295 (1974).

49. Lenardon, M.D., Sood, P., Dorfmueller, H.C., Brown, A.J.P. & Gow, N.A.R. Scalar nanostructure of the *Candida albicans* cell wall; a molecular, cellular and ultrastructural analysis and interpretation. The Cell Surface 6, 100047 (2020).

50. Gow, N.A.R. & Hube, B. Importance of the *Candida albicans* cell wall during commensalism and infection. Current Opinion in Microbiology 15, 406–412 (2012).

51. McKenzie, C.G. et al. Contribution of *Candida albicans* cell wall components to recognition by and escape from murine macrophages. Infect Immun 78, 1650–1658 (2010).

52. Bain, J.M. et al. *Candida albicans* hypha formation and mannan masking of β-glucan inhibit macrophage phagosome maturation. mBio 5, e01874–01814 (2014).

53. Yadav, B. et al. Differences in fungal immune recognition by monocytes and macrophages: N-mannan can be a shield or activator of immune recognition. The Cell Surface 6, 100042 (2020).

54. Jiang, H.-H. et al. Cell wall mannoprotein of *Candida albicans* polarizes macrophages and affects proliferation and apoptosis through activation of the Akt signal pathway. International Immunopharmacology 72, 308–321 (2019).

55. Gow, N.A.R., Brown, A.J.P. & Odds, F.C. Fungal morphogenesis and host invasion. Current Opinion in Microbiology 5, 366–371 (2002).

56. Sprague, J.L., Kasper, L. & Hube, B. From intestinal colonization to systemic infections: *Candida albicans* translocation and dissemination. Gut Microbes 14, 2154548 (2022).

57. Denning, D.W., Kneale, M., Sobel, J.D. & Rautemaa-Richardson, R. Global burden of recurrent vulvovaginal candidiasis: A systematic review. The Lancet Infectious Diseases 18, e339–e347 (2018).

58. Sobel, J.D. Vulvovaginal candidosis. The Lancet 369, 1961–1971 (2007).

59. Jacobsen, I.D. *Galleria mellonella* as a model host to study virulence of *Candida*. Virulence 5, 237–239 (2014).

60. Gu, W., Yu, Q., Yu, C. & Sun, S. *In vivo* activity of fluconazole/tetracycline combinations in *Galleria mellonella* with resistant *Candida albicans* infection. Journal of Global Antimicrobial Resistance 13, 74–80 (2018).

61. Li, D.-D. et al. Using *Galleria mellonella*-*Candida albicans* infection model to evaluate antifungal agents. Biological and Pharmaceutical Bulletin 36, 1482–1487 (2013).

62. Frei, A. et al. Metal complexes as antifungals? From a crowd-sourced compound library to the first *in vivo* experiments. JACS Au 2, 2277–2294 (2022).

63. Selmecki, A., Forche, A. & Berman, J. Aneuploidy and isochromosome formation in drug-resistant *Candida albicans*. Science 313, 367–370 (2006).

64. Yang, F. et al. The fitness costs and benefits of trisomy of each *Candida albicans* chromosome. Genetics 218 (2021).

65. Selmecki, A., Gerami-Nejad, M., Paulson, C., Forche, A. & Berman, J. An isochromosome confers drug resistance *in vivo* by amplification of two genes, *ERG11* and *TAC1*. Molecular Microbiology 68, 624–641 (2008).

66. Yang, F. et al. Tunicamycin potentiates antifungal drug tolerance *via* aneuploidy in *Candida albicans*. mBio 12, e02272–02221 (2021).

67. Wei, M.-X. et al. Enantioselective synthesis of *Amaryllidaceae* alkaloids (+)-vittatine, (+)-epi-vittatine, and (+)-buphanisine. Chemistry – An Asian Journal 8, 1966–1971 (2013).

68. Nguyen, T.-K. et al. Rational design of single-chain polymeric nanoparticles that kill planktonic and biofilm bacteria. ACS Infectious Diseases 3, 237–248 (2017).

69. Phuong, P.T. et al. Effect of hydrophobic groups on antimicrobial and hemolytic activity: Developing a predictive tool for ternary antimicrobial polymers. Biomacromolecules 21, 5241–5255 (2020).

70. Judzewitsch, P.R., Zhao, L., Wong, E.H.H. & Boyer, C. High-throughput synthesis of antimicrobial copolymers and rapid evaluation of their bioactivity. Macromolecules 52, 3975–3986 (2019).

71. Wächtler, B., Wilson, D., Haedicke, K., Dalle, F. & Hube, B. From attachment to damage: Defined genes of *Candida albicans* mediate adhesion, invasion and damage during interaction with oral epithelial cells. PLOS ONE 6, e17046 (2011).

72. Yu, G.W., Li-Gen Han, Yanyan; He, Qing-Yu clusterProfiler: An R package for comparing biological themes among gene clusters. OMICS: A Journal of Integrative Biology 16, 284–287 (2012).

73. Wu, T. et al. clusterProfiler 4.0: A universal enrichment tool for interpreting omics data. The Innovation 2, 100141 (2021).

74. Schneider, C.A., Rasband, W.S. & Eliceiri, K.W. NIH Image to ImageJ: 25 years of image analysis. Nature Methods 9, 671–675 (2012).

75. Giard, D.J. et al. *In vitro* cultivation of human tumors: Establishment of cell lines derived from a series of solid tumors. Journal of the National Cancer Institute 51, 1417–1423 (1973).

76. Hernandez, R. & Rupp, S. in Host-Pathogen Interactions: Methods and Protocols. (eds. S. Rupp & K. Sohn) 105–123 (Humana Press, Totowa, NJ; 2009).

77. Schaller, M., Zakikhany, K., Naglik, J.R., Weindl, G. & Hube, B. Models of oral and vaginal candidiasis based on *in vitro* reconstituted human epithelia. Nature Protocols 1, 2767–2773 (2006).

78. Pekmezovic, M. et al. *Candida* pathogens induce protective mitochondria-associated type I interferon signalling and a damage-driven response in vaginal epithelial cells. Nature Microbiology 6, 643–657 (2021).

79. Frei, A. et al. Platinum cyclooctadiene complexes with activity against Gram-positive bacteria. ChemMedChem 16, 3165–3171 (2021).

80. Tsai, C.J.-Y., Loh, J.M.S. & Proft, T. *Galleria mellonella* infection models for the study of bacterial diseases and for antimicrobial drug testing. Virulence 7, 214–229 (2016).

81. Siscar-Lewin, S. et al. Transient mitochondria dysfunction confers fungal cross-resistance against phagocytic killing and fluconazole. mBio 12, e01128–01121 (2021).

82. Cingolani, P. et al. A program for annotating and predicting the effects of single nucleotide polymorphisms, SnpEff. Fly 6, 80–92 (2012).

83. Abbey, D.A. et al. YMAP: A pipeline for visualization of copy number variation and loss of heterozygosity in eukaryotic pathogens. Genome Medicine 6, 100 (2014).

84. Robinson, J.T., et al. Integrative genomics viewer. Nature Biotechnology 29, 24–26 (2011).

85. Hahne, F. & Ivanek, R. in Statistical Genomics: Methods and Protocols. (eds. E. Mathé & S. Davis) 335–351 (Springer New York, New York, NY; 2016).

