## Supplementary Material for "A synthetic peptide mimic kills *Candida albicans* and synergistically prevents infection"

### Size exclusion chromatography data of polymers

SEC molecular weight distributions were calculated based on poly(methyl methacrylate) (PMMA) standards and the resulting 3rd order calibration curve.

$$Mn = -0.001287685x^3 + 0.09233964x^2 - 2.345552x + 25.20845$$

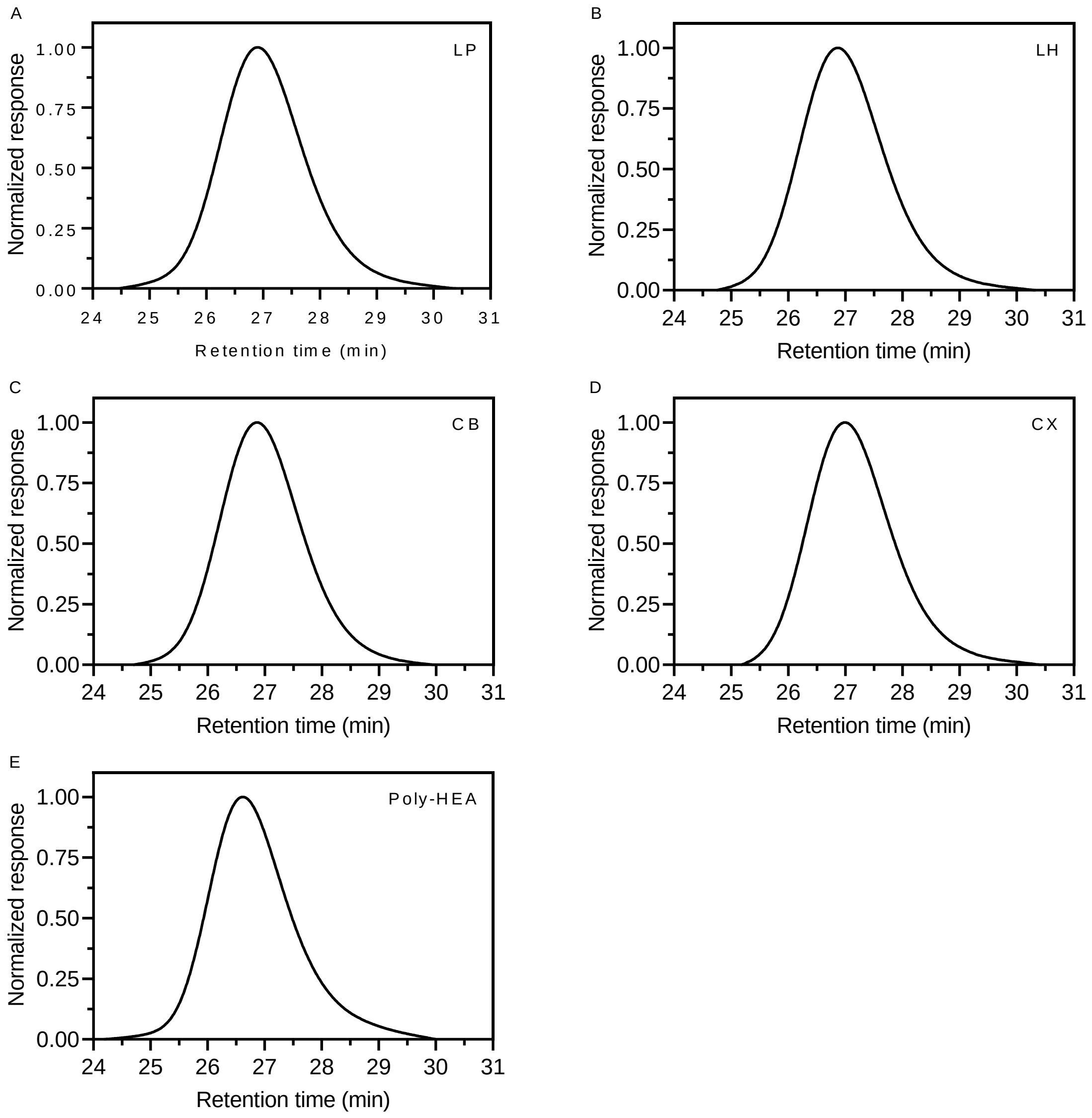

**Figure S1** – SEC spectra of the polymers (A) LP, (B) LH, (C) CB, (D) CX, and (E) poly-HEA.

### $^1$H NMR spectroscopy data of polymer synthesis

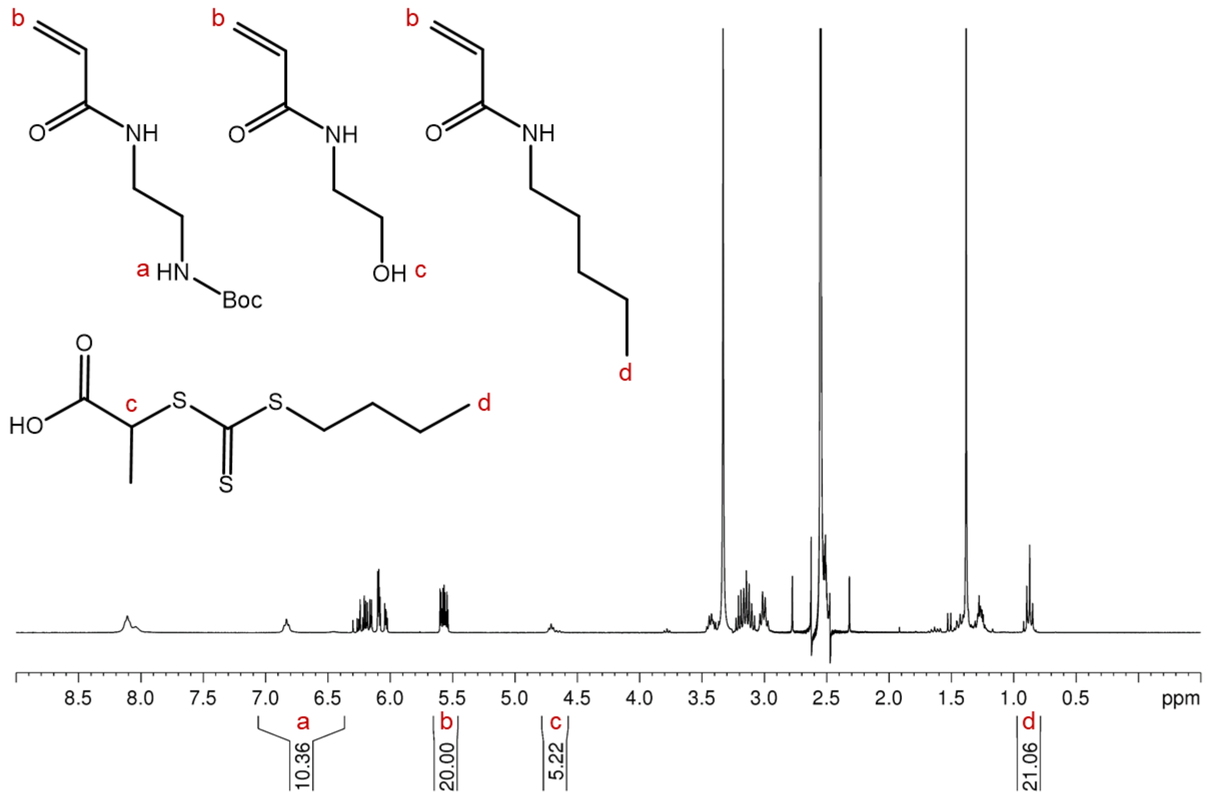

**Figure S2** – $^1$H NMR of reaction mixture before polymerisation to synthesise polymer LP. The composition was calculated as outlined below based on the characteristic signals a, c, and d, which were normalised to b ($X_n$=20). Solvent: DMSO-$d_6$.

$$\%\,Positively\ charged\ monomer = \frac{\int a}{\int a + (c-1) + \left(\frac{d-3}{3}\right)} \approx 50.3\%$$

$$\%\,HEAm = \frac{\int (c-1)}{\int a + (c-1) + \left(\frac{d-3}{3}\right)} \approx 20.5\%$$

$$\%\,Hydrophobic\ monomer = \frac{\int (\frac{(d-3)}{3})}{\int a + (c-1) + \left(\frac{d-3}{3}\right)} \approx 29.2\%$$

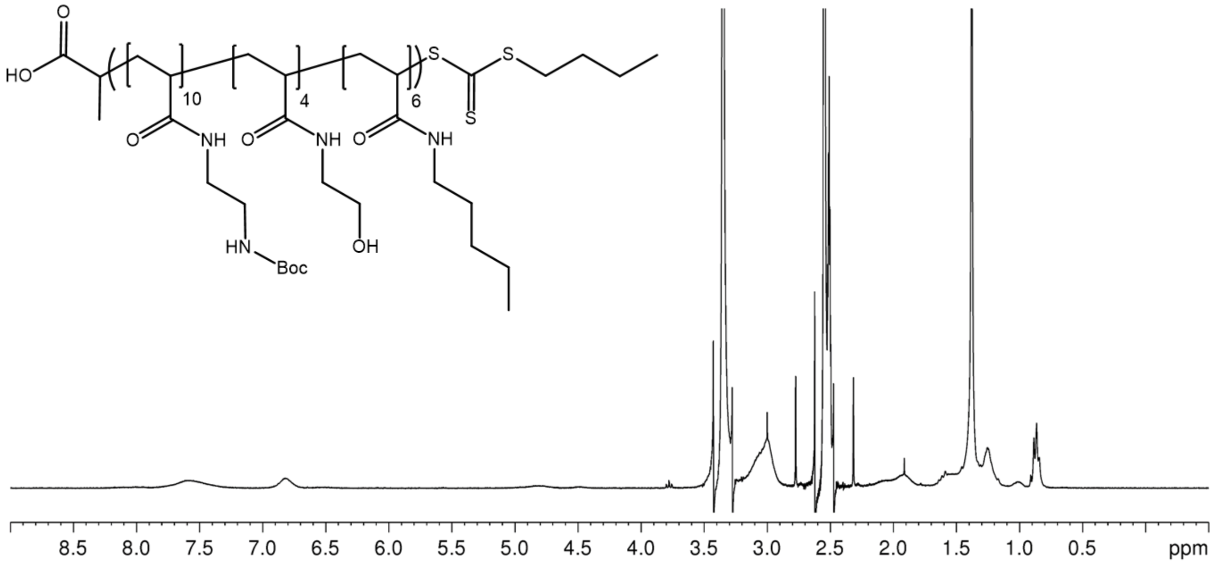

**Figure S3** – $^{1}$H NMR after 15 h polymerisation of LP to check monomer conversion resulting in fading of the signal between 5.5 and 6.35 ppm and appearance of the polymer backbone signal between 1.1 and 2.2 ppm. Solvent: DMSO-$d_6$.

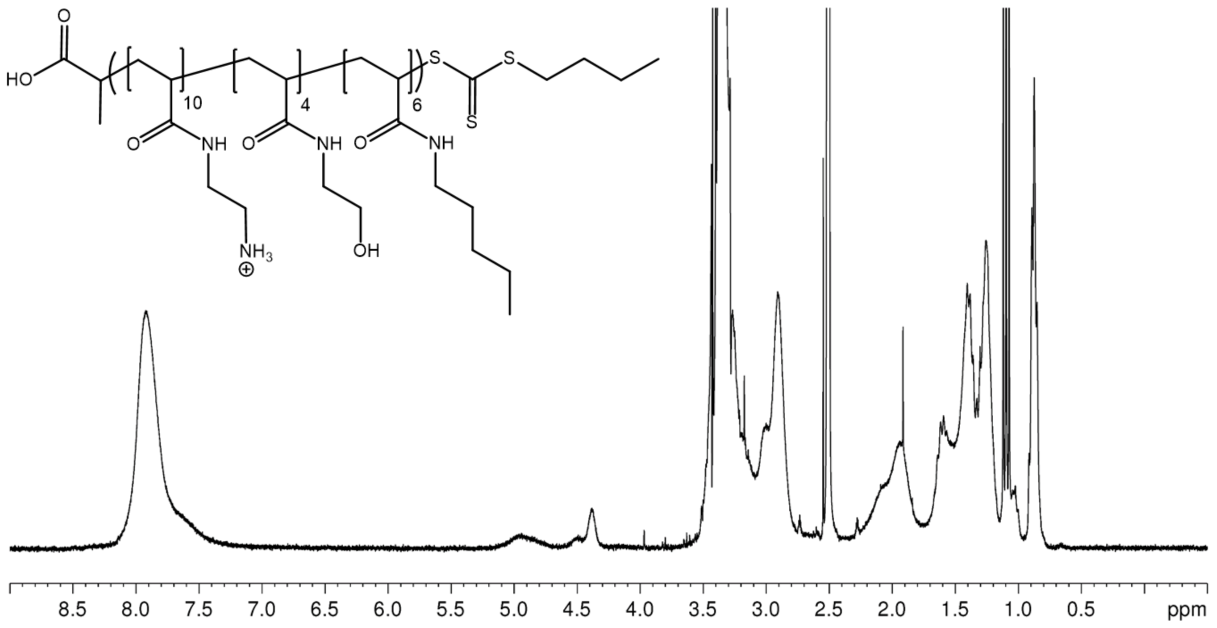

**Figure S4** – $^{1}$H NMR after Boc-deprotection of LP resulting in fading of the signal between 6.35 and 7 ppm and appearance of a signal between 7.7 and 8.4 ppm. Solvent: DMSO-$d_6$.

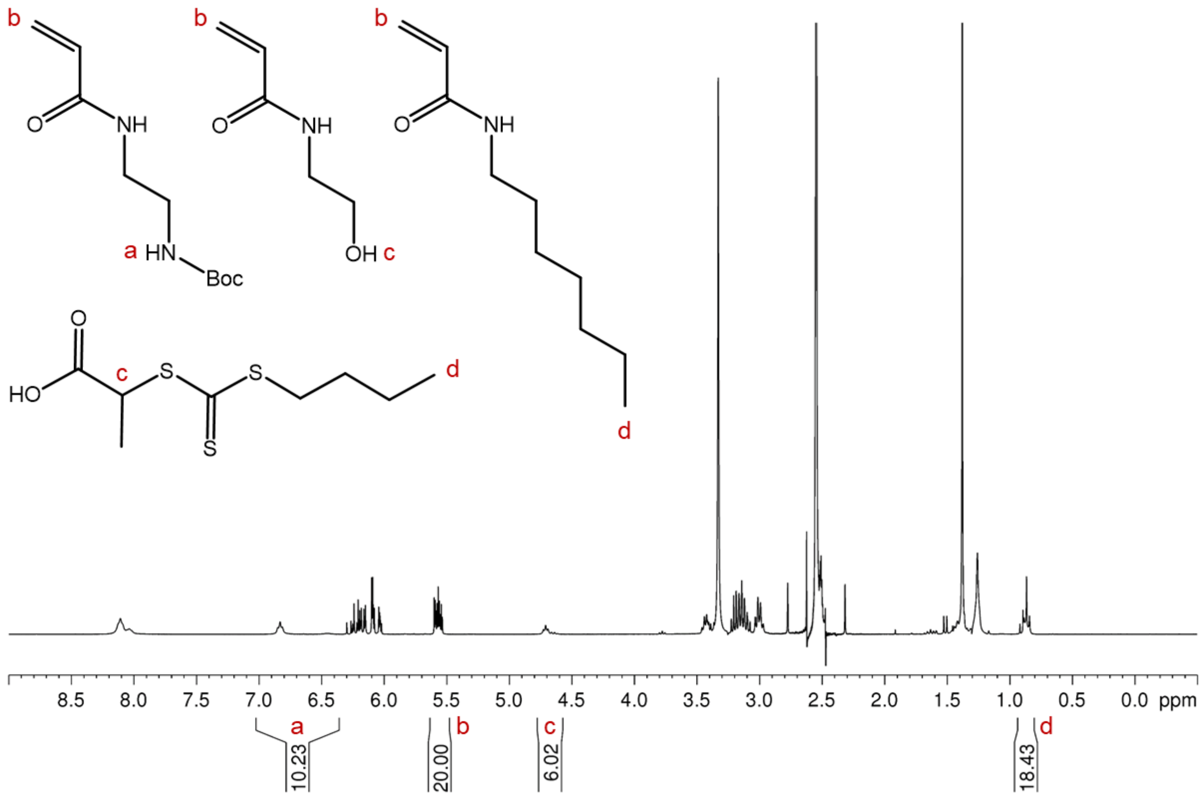

**Figure S5** – [1]H NMR of reaction mixture before polymerisation to synthesise polymer LH. The composition was calculated as outlined below based on the characteristic signals a, c, and d, which were normalised to b ($X_n$=20). Solvent: DMSO-$d_6$.

$$\%\ Positively\ charged\ monomer = \frac{\int a}{\int a + (c-1) + \left(\frac{d-3}{3}\right)} \approx 50.2\%$$

$$\%\ HEAm = \frac{\int (c-1)}{\int a + (c-1) + \left(\frac{d-3}{3}\right)} \approx 24.6\%$$

$$\%\ Hydrophobic\ monomer = \frac{\int (\frac{(d-3)}{3})}{\int a + (c-1) + \left(\frac{d-3}{3}\right)} \approx 25.2\%$$

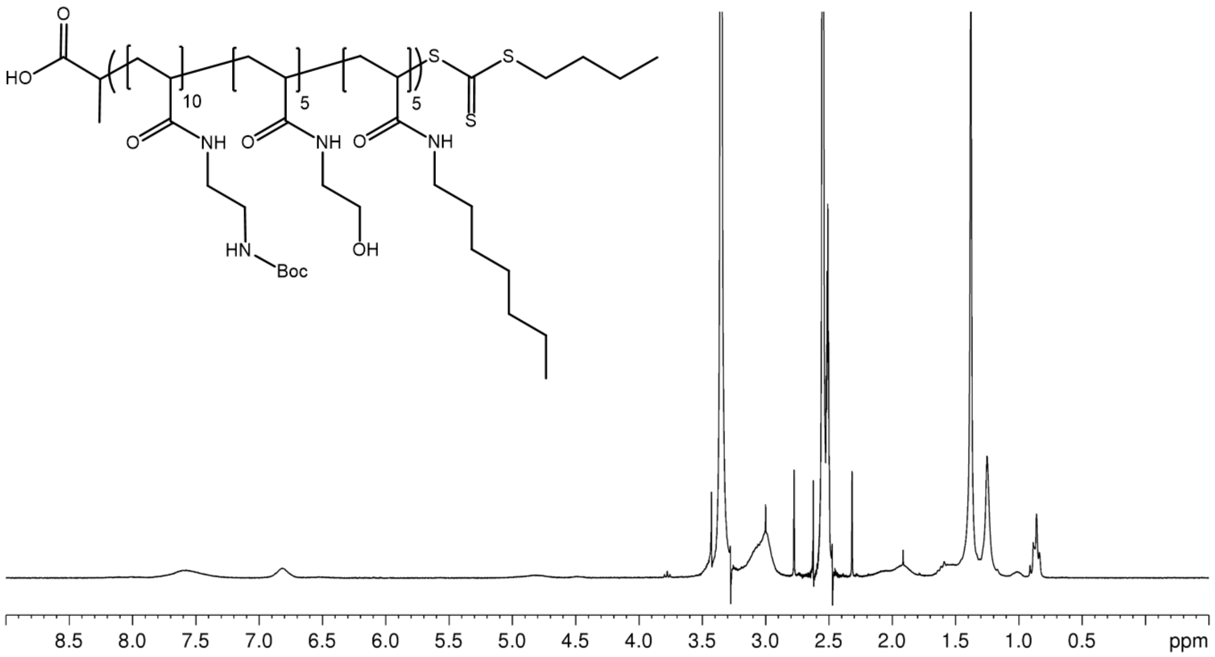

**Figure S6** – [1]H NMR after 15 h polymerisation of LH to check monomer conversion resulting in fading of the signal between 5.5 and 6.35 ppm and appearance of the polymer backbone signal between 1.1 and 2.2 ppm. Solvent: DMSO-d6.

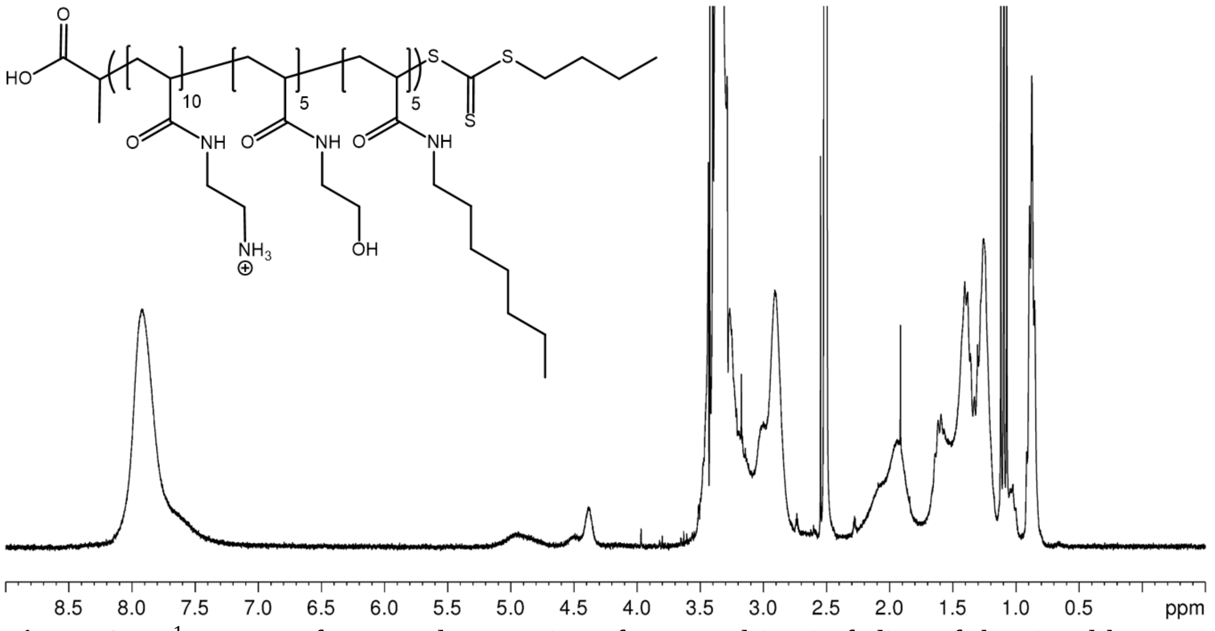

**Figure S7** – [1]H NMR after Boc-deprotection of LH resulting in fading of the signal between 6.35 and 7 ppm and appearance of a signal between 7.7 and 8.4 ppm. Solvent: DMSO-d6.

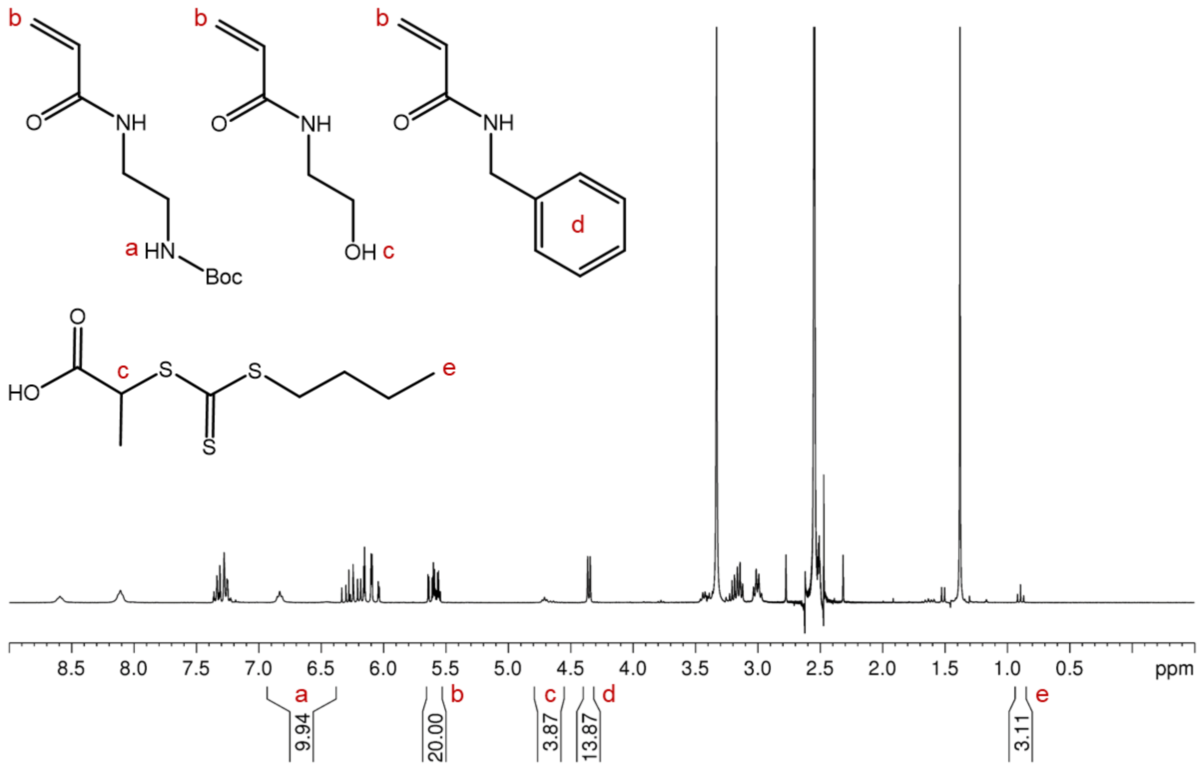

**Figure S8** – [1]H NMR of reaction mixture before polymerisation to synthesise polymer CB. The composition was calculated as outlined below based on the characteristic signals a, c, and d, which were normalised to b ($X_n$=20). The addition of the correct amount of BTPA was confirmed by e. Solvent: DMSO-$d_6$.

$$\%\,Positively\ charged\ monomer = \frac{\int a}{\int a + (c-1) + d/2} \approx 50.3\%$$

$$\%\,HEAm = \frac{\int (c-1)}{\int a + (c-1) + d/2} \approx 14.6\%$$

$$\%\,Hydrophobic\ monomer = \frac{\int d/2}{\int a + (c-1) + d/2} \approx 35.1\%$$

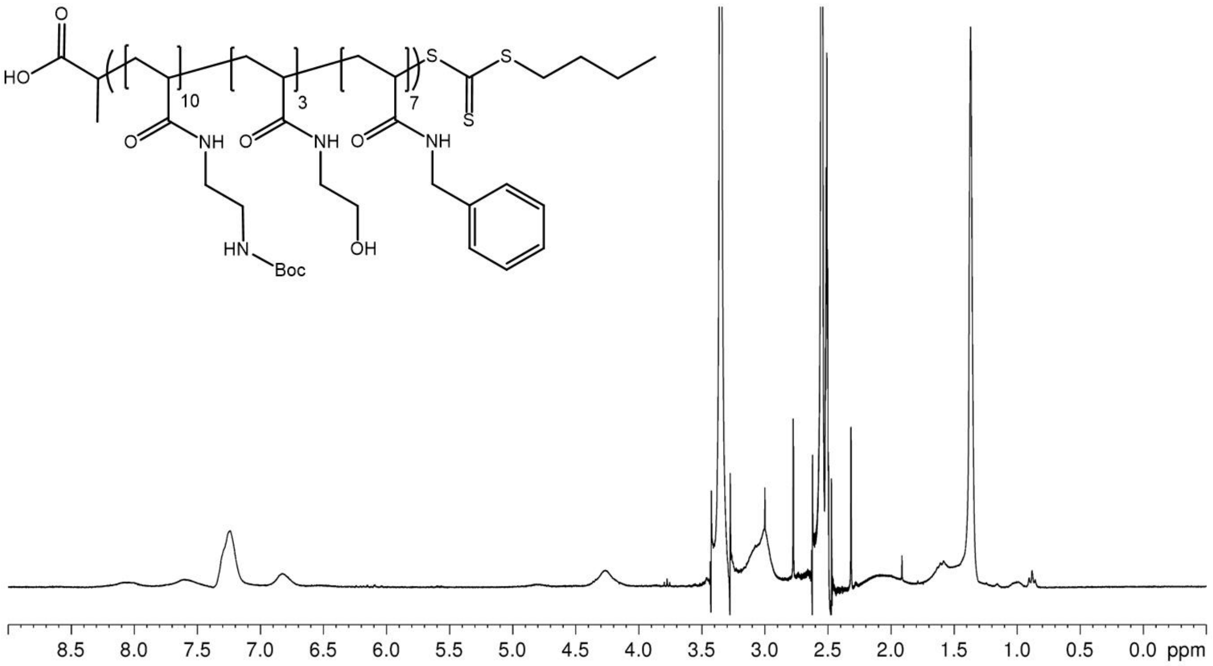

**Figure S9** – $^{1}$H NMR after 15 h polymerisation of CB to check monomer conversion resulting in fading of the signal between 5.5 and 6.35 ppm and appearance of the polymer backbone signal between 1.1 and 2.2 ppm. Solvent: DMSO-$d_6$.

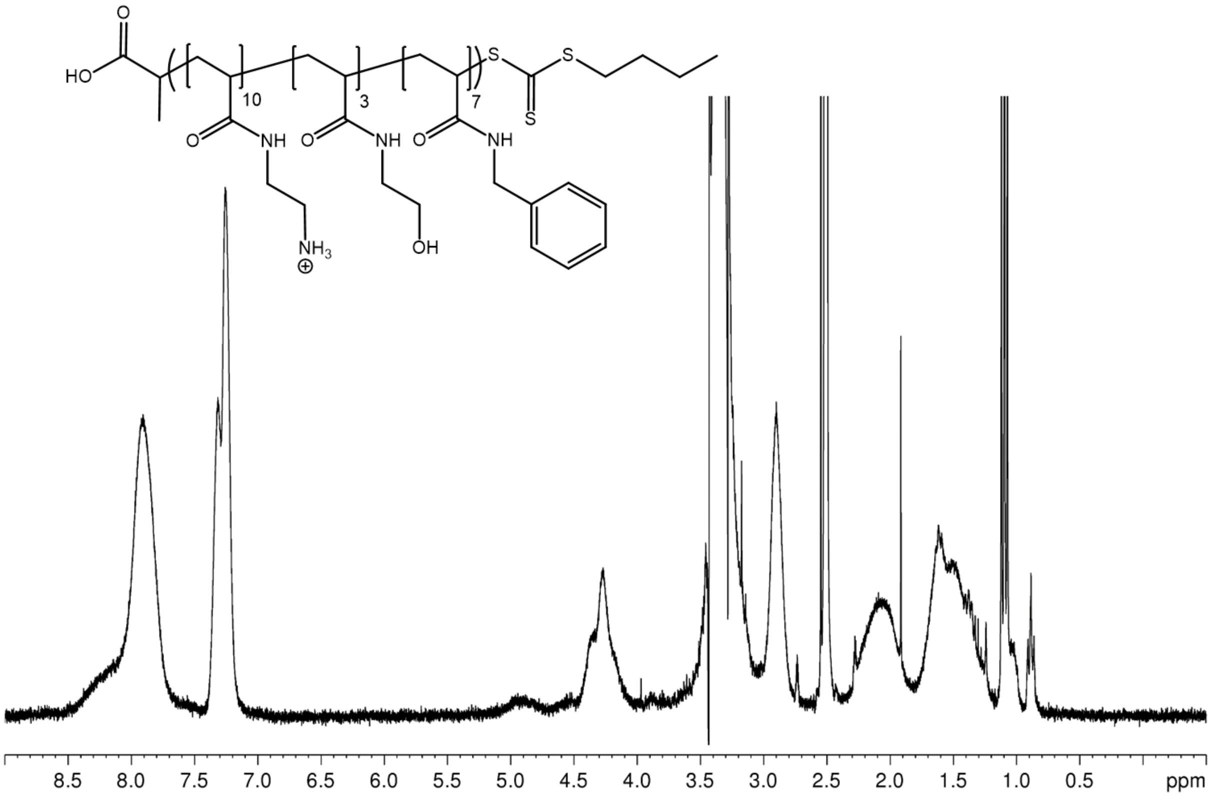

**Figure S10** – $^{1}$H NMR after Boc-deprotection of CB resulting in fading of the signal between 6.35 and 7 ppm and appearance of a signal between 7.7 and 8.4 ppm. Solvent: DMSO-$d_6$.

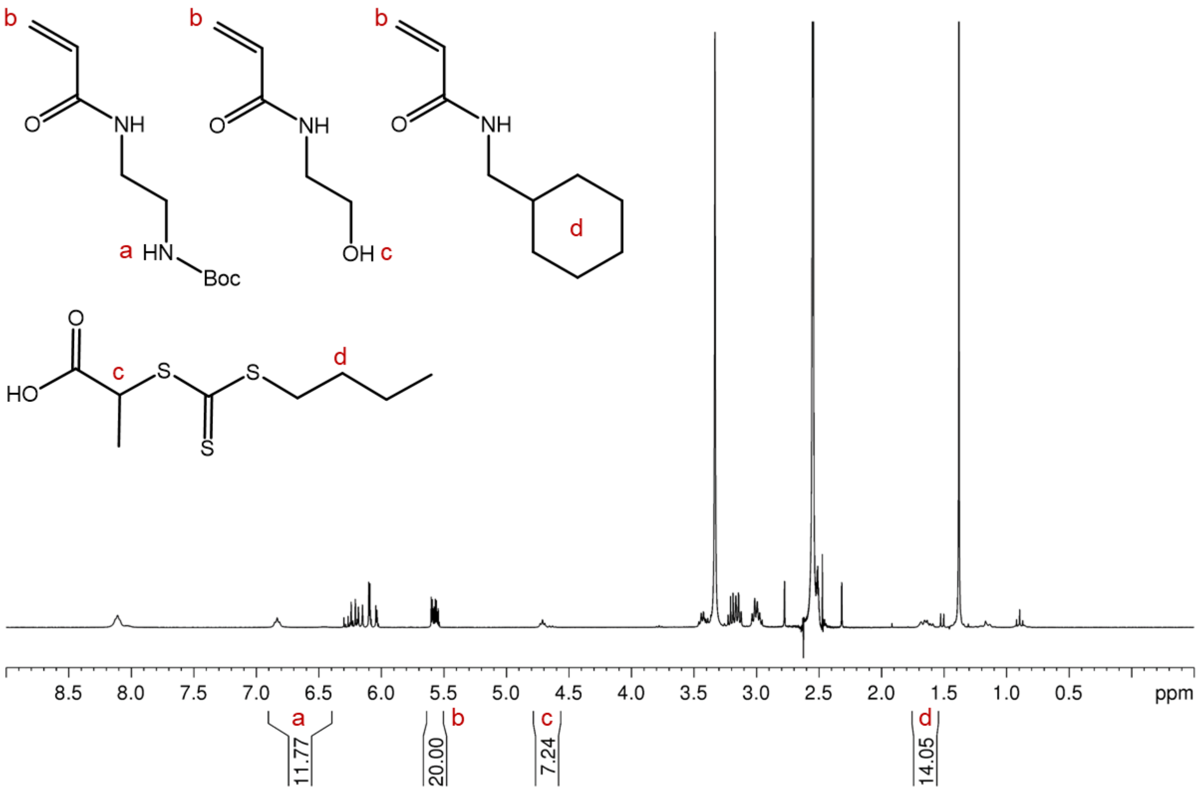

**Figure S11** – [1]H NMR of reaction mixture before polymerisation to synthesise polymer CX. The composition was calculated as outlined below based on the characteristic signals a, c, and d, which were normalised to b ($X_n$=20). Solvent: DMSO-$d_6$.

$$\%\ Positively\ charged\ monomer = \frac{\int a}{\int a + (c-1) + (\frac{d-2}{2})} \approx 49.0\%$$

$$\%\ HEAm = \frac{\int (c-1)}{\int a + (c-1) + (\frac{d-2}{2})} \approx 26.0\%$$

$$\%\ Hydrophobic\ monomer = \frac{\int \frac{d-2}{2}}{\int a + (c-1) + (\frac{d-2}{2})} \approx 25.0\%$$

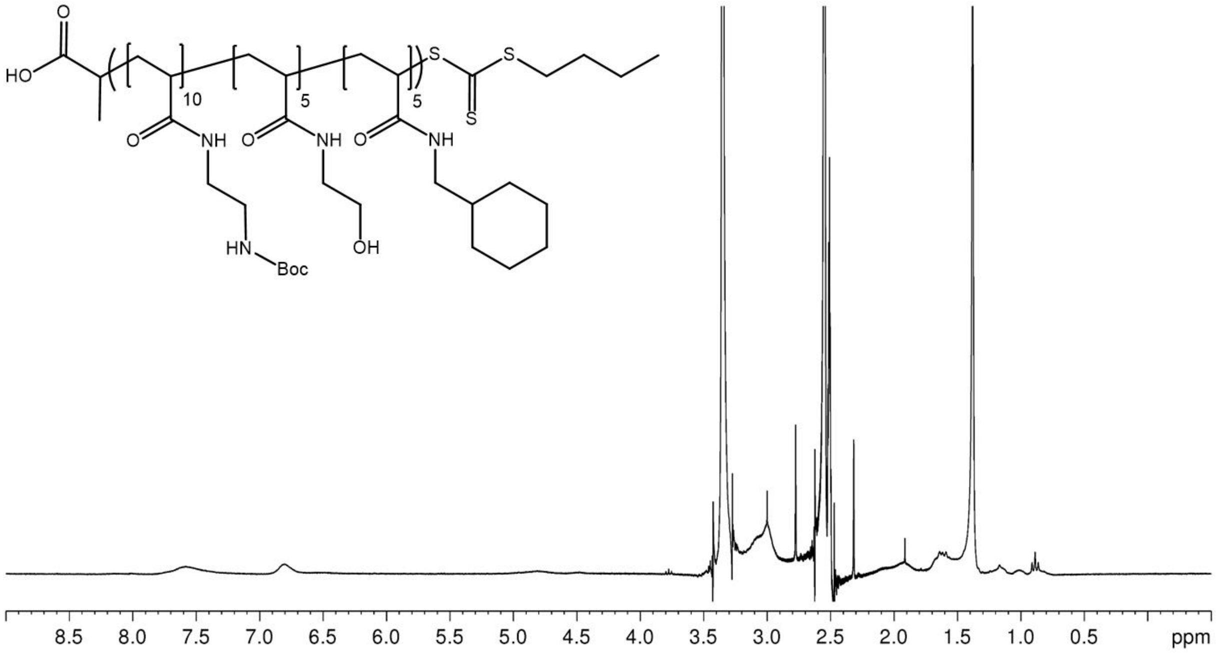

**Figure S12** – $^{1}$H NMR after 15 h polymerisation of CX to check monomer conversion resulting in fading of the signal between 5.5 and 6.35 ppm and appearance of the polymer backbone signal between 1.1 and 2.2 ppm. Solvent: DMSO-$d_6$.

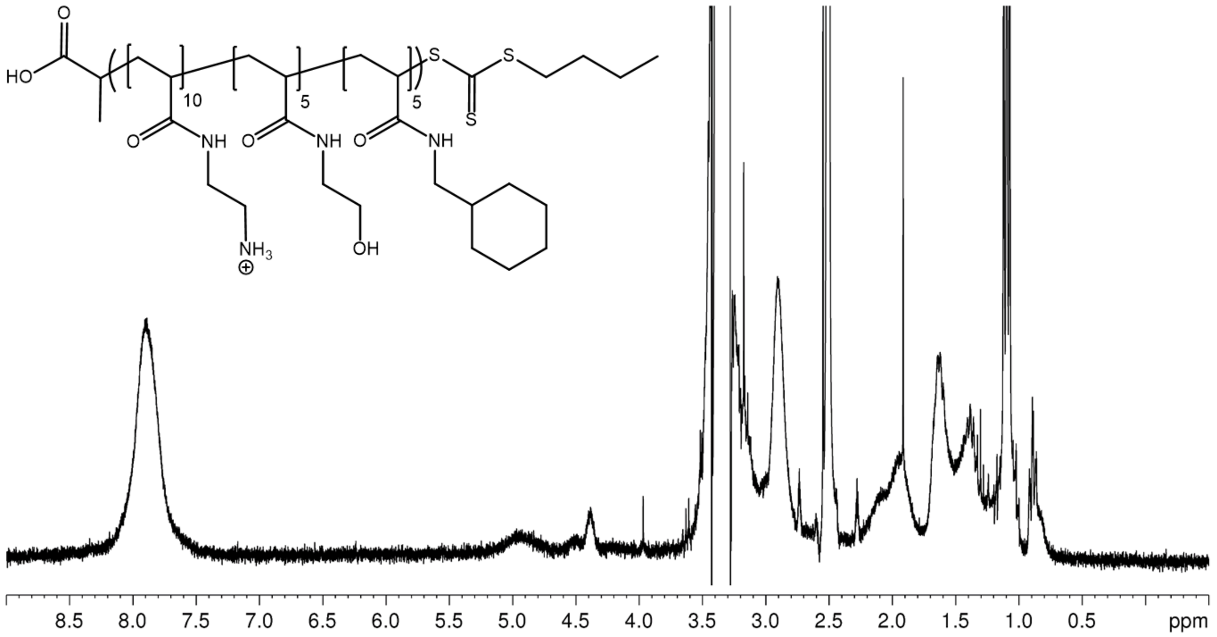

**Figure S13** – $^{1}$H NMR after Boc-deprotection of CX resulting in fading of the signal between 6.35 and 7 ppm and appearance of a signal between 7.7 and 8.4 ppm. Solvent: DMSO-$d_6$.

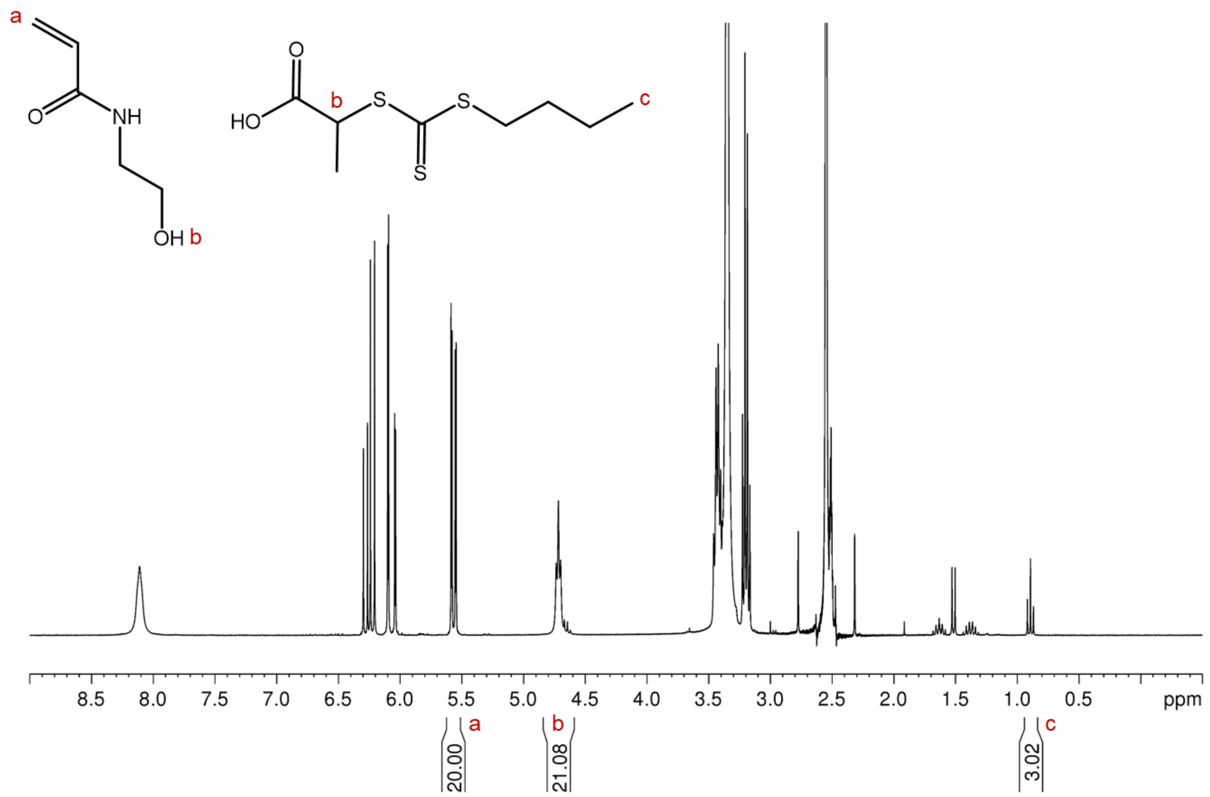

**Figure S14** – [1]H NMR of reaction mixture before polymerisation to synthesise polymer poly-HEA. The composition was calculated as outlined below based on the characteristic signal b which was normalised to b ($X_n$=20). The addition of the correct amount of BTPA was confirmed by c. Solvent: DMSO-$d_6$.

$$\% \, HEAm = \frac{\int(b-1)}{\int a} \approx 100.4\%$$

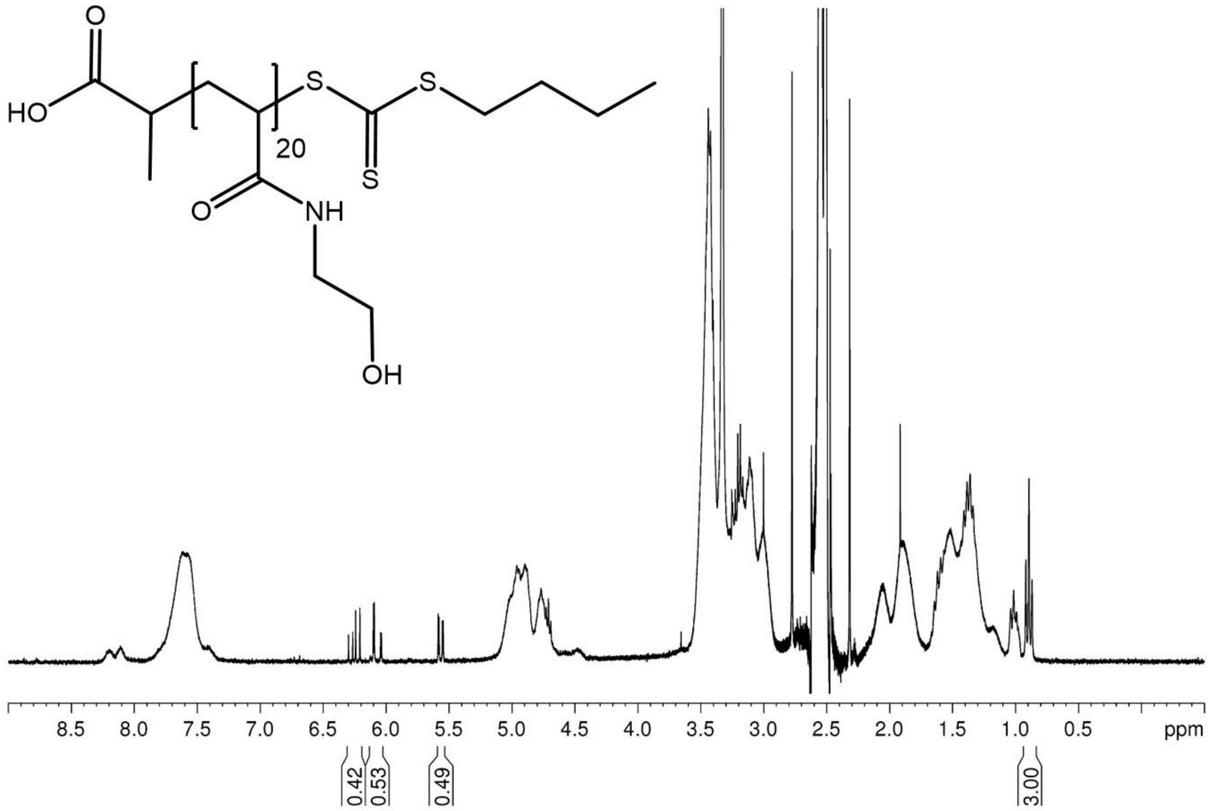

**Figure S15 -** [1]H NMR after 15 h polymerisation of poly-HEA to check monomer conversion resulting in fading of the signal between 5.5 and 6.35 ppm and appearance of the polymer backbone signal between 1.1 and 2.2 ppm. Exemplary monomer conversion calculation is described below. Solvent: DMSO-$d_6$.

$$Average\ of\ remaining\ monomer\ integrals\ between\ 5.5\ and\ 6.35\ ppm: 0.48$$

$$Monomer\ integral\ before\ polymerization\ (see\ \boldsymbol{Figure\ S14}): 20 * \frac{3.02}{3.00} = 20.133$$

$$Monomer\ conversion: 1 - \frac{0.48}{20.133} \approx 97.6\%$$

### Characterisation of the non-toxic polymer poly-HEA

**Table S1** - Composition, molecular weight, and dispersity of the non-toxic control polymer, consisting of the hydrophilic, uncharged hydroxyethyl acrylamide (poly-HEA).

| **Polymer** | **Targeted polymer composition (% positively charged / hydrophilic / hydrophobic functionality)** | **Polymer composition (% positively charged / hydrophilic / hydrophobic functionality)[a]** | **Targeted** $X_n$ | **Theoretical** $M$ **(g/mol)** [b] | $M_n$ **(g/mol)** [c] | $Đ$ [c] |
|---|---|---|---|---|---|---|
| poly-HEA | 0 / 100 / 0 | 0 / 100 / 0 | 20 | 2,600 | 6,900 | 1.08 |

*Note*:
[a] determined by $^1$H NMR spectroscopy before polymerisation
[b] calculated based on targeted composition and $X_n$, rounded to the nearest 100.
[c] determined by SEC using poly(methyl methacrylate) standards

### Transcriptomic response of *C. albicans* to polymers

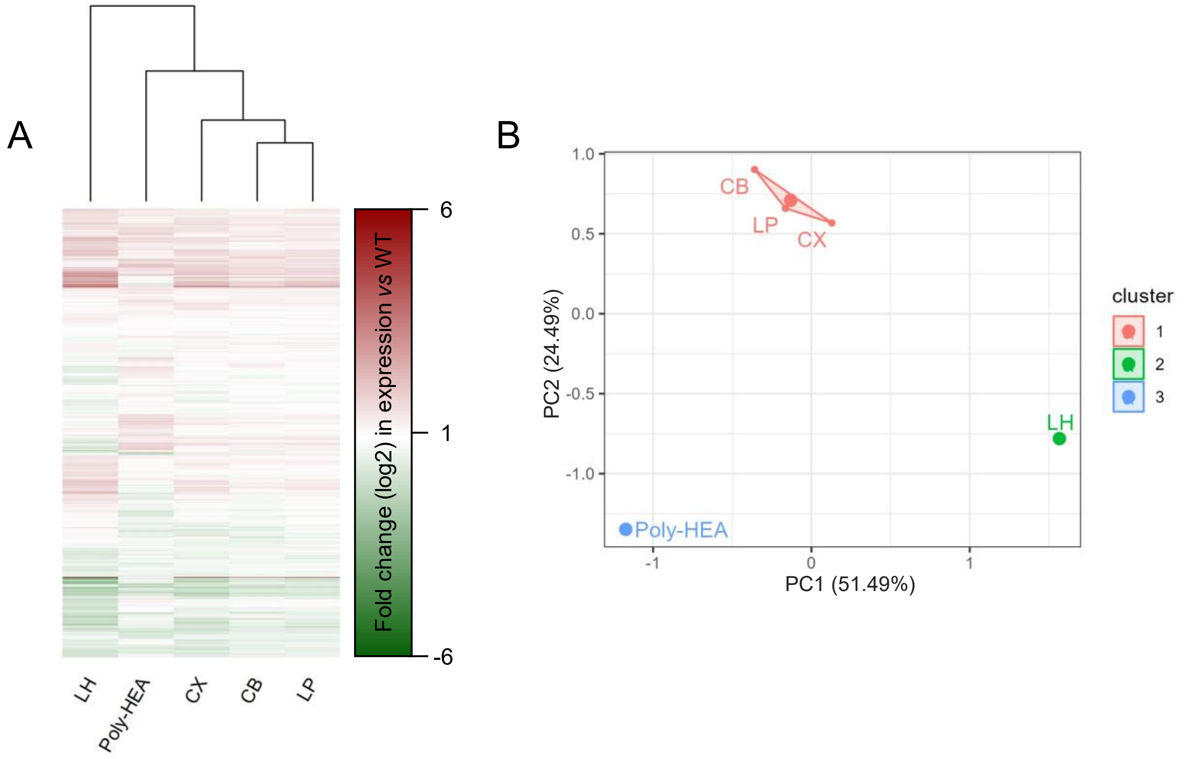

**Figure S16** – Global transcriptomic response of *C. albicans* SC5314 to antifungal polymers visualised (A) as a heatmap and (B) by principal component analysis and k-mean clustering after dimensionality reduction. The tree in the heatmap (A) with polymers as leaves depicts their hierarchical clustering where the height of each node is proportional to the dissimilarity (Euclidian distance) between their transcriptional response signatures.

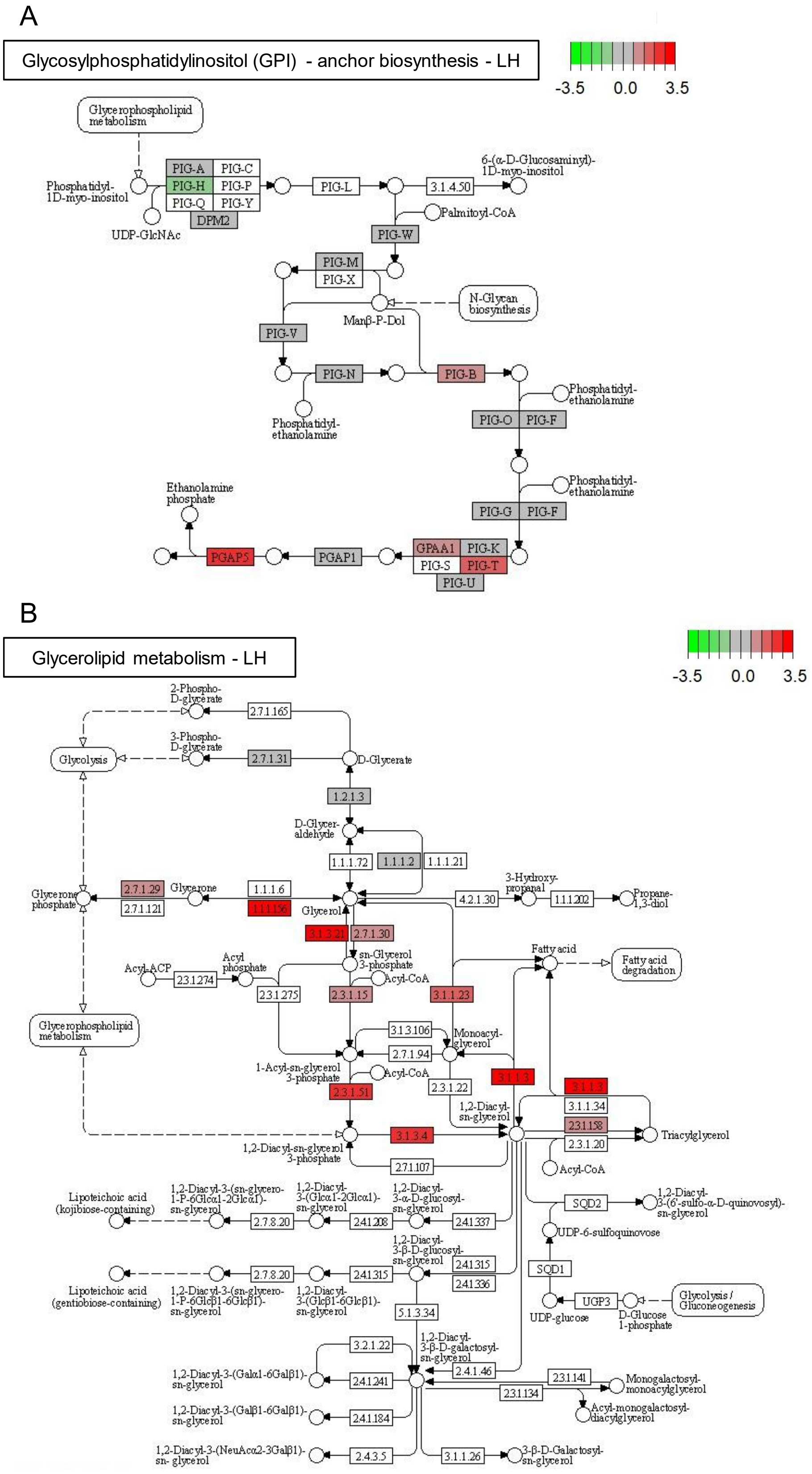

**Figure S17.** KEGG pathway maps for (A) glycosylphosphatidylinositol(GPI)-anchor biosynthesis and (B) glycerolipid metabolism KEGG pathways and respectively up- (red) and downregulated (green) genes in *C. albicans* SC5314 after treatment with polymer LH. The scale bars represent the log2 fold change in gene expression compared to untreated cells.

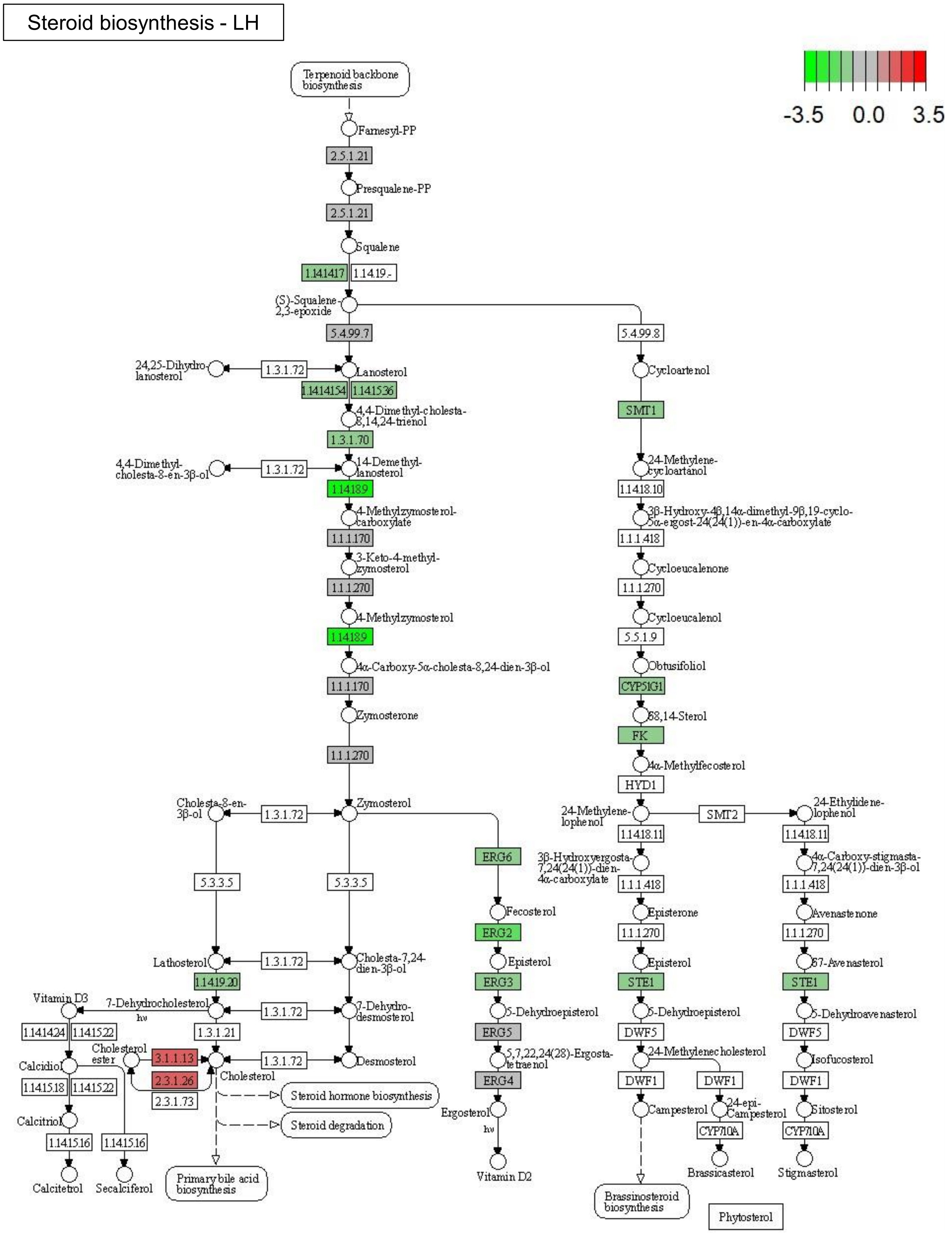

**Figure S18.** KEGG pathway map of steroid biosynthesis and respectively up- (red) and downregulated (green) genes in *C. albicans* SC5314 after treatment with polymer LH. The scale bar represents the log2 fold change in gene expression compared to untreated cells.

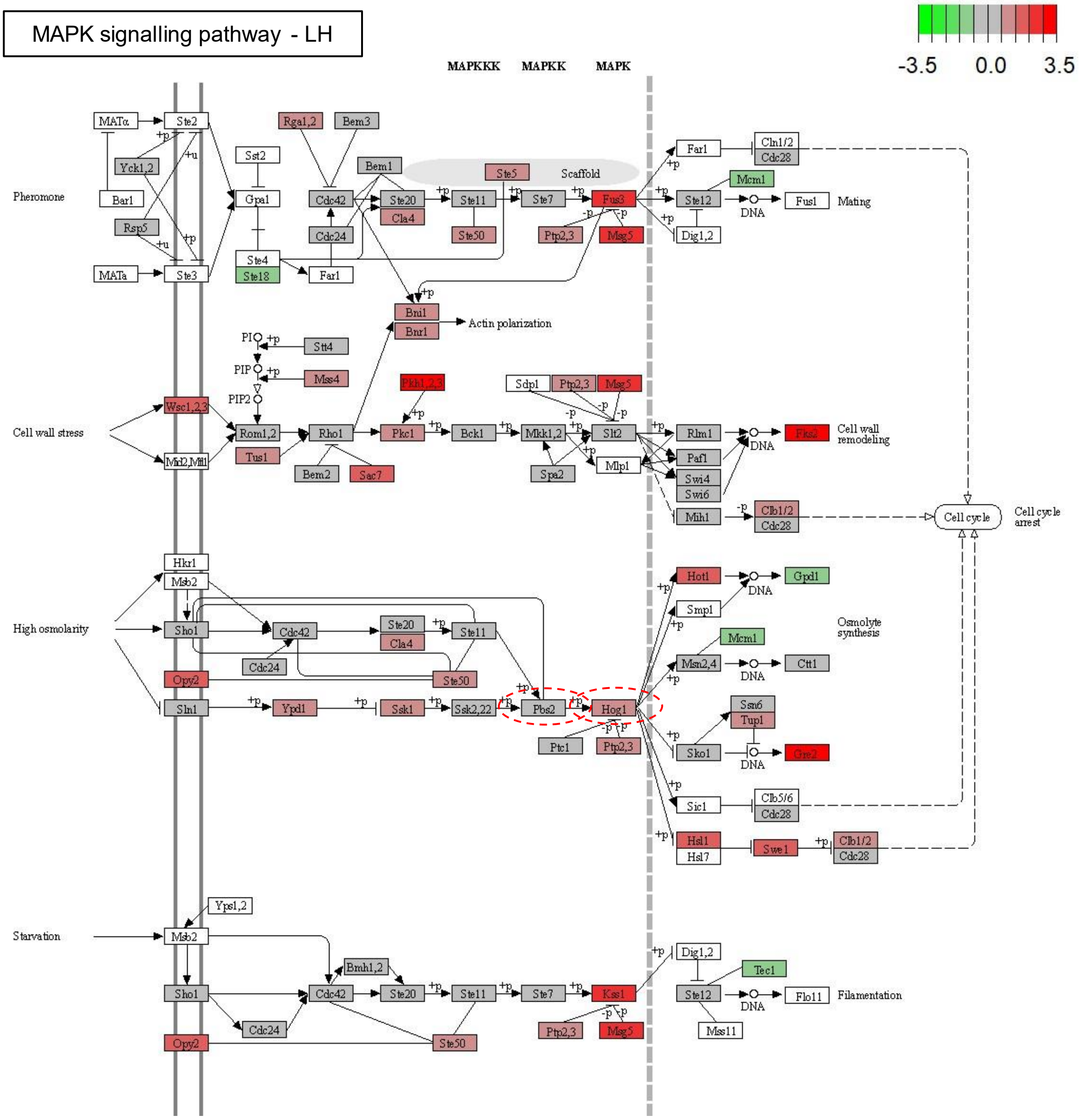

**Figure S19** –KEGG pathway map of MAPK signalling and respectively up- (red) and downregulated (green) genes in *C. albicans* SC5314 after treatment with polymer LH. The red-encircled genes represent genes which have been found to be crucial for *C. albicans* to respond to LH-induced stress in the deletion mutant screen. The scale bar represents the log2 fold change in gene expression compared to untreated cells.

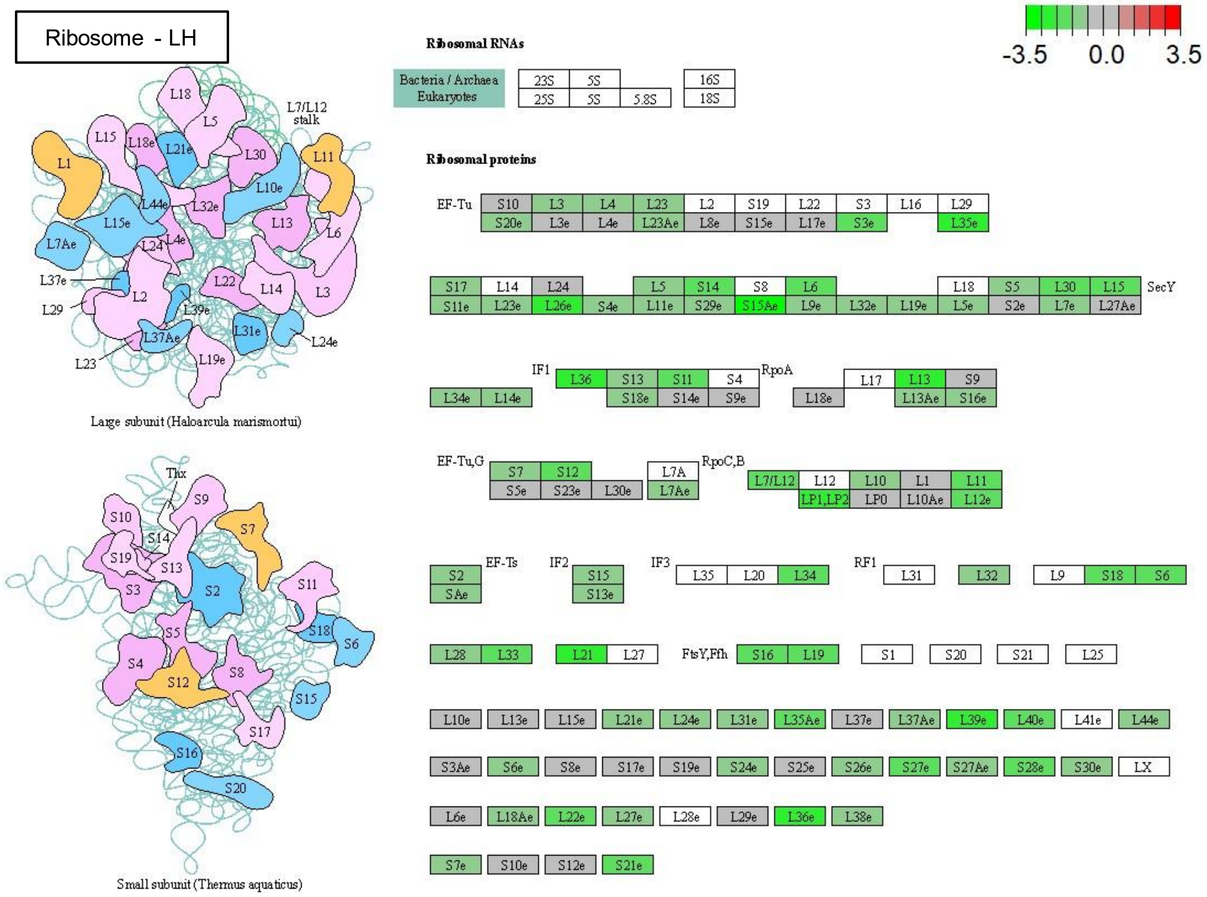

**Figure S20.** KEGG pathway map of ribosome and respectively non-different (grey) or downregulated genes (green) in *C. albicans* SC5314 after treatment with polymer LH.

### Mutant screening and growth speed index

**Table S2** – Growth speed index of selected *C. albicans* (black) and *C. glabrata* (blue) mutants.

| Mutant | ORF/systematic name | LH | LP | CB | CX |
|---|---|---|---|---|---|
| **Cell wall organisation and stress response** | | | | | |
| *chk1* | orf19.896 | 0.03 | -0.04 | 0.06 | -0.05 |
| *chs4* | orf19.7349 | -0.01 | 0.01 | -0.04 | 0 |
| *cis2* | orf19.6053 | -0.02 | 0 | 0.01 | 0.04 |
| *ire1* | orf19.5068 | NG | 0.04 | 0.01 | 0.09 |
| *mkc1* | orf19.7532 | -0.01 | -0.13 | 0.01 | -0.14 |
| *pir1* | orf19.220 | 0.1 | 0.1 | 0.01 | 0.14 |
| *rlm1* | orf19.4662 | 0.16 | -0.04 | 0.04 | -0.02 |
| *sun41* | orf19.3642 | 0.02 | 0 | 0.01 | 0.09 |
| *chs2* | CAGL0J11506g | -0.02 | -0.02 | 0.02 | -0.08 |
| *chs3* | CAGL0B04389g | -0.02 | 0.09 | 0.03 | 0.04 |
| *fks2* | CAGL0K04037g | -0.17 | -0.18 | -0.13 | -0.14 |
| *fks3* | CAGL0M13827g | -0.05 | 0 | 0.02 | -0.08 |
| *hac1* | CAGL0K12540g | -0.06 | 0.22 | 0.2 | -0.03 |
| *ire1* | CAGL0F03245g | 0.18 | -0.02 | -0.04 | -0.09 |
| *mkk2* | CAGL0J03828g | -0.08 | -0.15 | -0.27 | -0.11 |
| *pir3* | CAGL0M08492g | -0.04 | -0.03 | -0.04 | -0.05 |
| *pir4* | CAGL0I06160g | 0.06 | -0.03 | -0.05 | -0.01 |
| *rlm1* | CAGL0H05621g | -0.01 | -0.05 | -0.04 | 0.02 |
| *rom2* | CAGL0G04873g | -0.75 | -0.74 | -0.46 | -0.67 |
| *slt2* | GAGL0J00539g | -0.08 | -0.13 | -0.22 | 0.03 |
| **Membrane composition and inositol signaling** | | | | | |
| *erg5* | orf19.5178 | -0.1 | -0.1 | -0.16 | -0.15 |
| *ino4* | orf19.837.1 | 0.08 | -0.11 | 0.02 | -0.1 |
| *inp51* | orf19.1373 | NG | NG | NG | NG |
| *upc2* gof | orf19.391 | -0.04 | 0.02 | 0.05 | 0.06 |
| *erg5* | CAGL0M07656g | 0.3 | 0.5 | 0.24 | 0.32 |
| *ino2* | CAGL0B01947g | 0.53 | 0.74 | 0.63 | 0.49 |
| *inp51* | CAGL0J02134g | -0.4 | -0.1 | -0.24 | -0.26 |
| *inp53* | CAGL0B04631g | -0.29 | -0.25 | 0.04 | -0.21 |
| *isc1* | CAGL0E06556g | -0.63 | -0.65 | -0.2 | -0.56 |

| | | | | | |
|---|---|---|---|---|---|
| **Protein glycosylation - *O*-linked mannosylation** | | | | | |
| *mnt1* | orf19.1665 | 0 | 0.02 | -0.05 | 0.01 |
| *pmt1* | orf19.5171 | -1.08 | -0.63 | -0.53 | -0.67 |
| *pmt2** | orf19.6812 | -0.07 | 0.02 | 0.16 | 0.02 |
| *pmt3* | orf19.3802 | -0.23 | -0.19 | -0.09 | -0.21 |
| *pmt4* | orf19.4109 | -0.51 | -0.48 | -0.33 | -0.49 |
| *pmt5* | orf19.7549 | -0.52 | -0.43 | -0.31 | -0.48 |
| *pmt1* | CAGL0L07216g | -0.75 | -0.22 | -1.12 | -0.63 |
| *pmt2* | CAGL0J08734g | -1.57 | -0.98 | -1.53 | -1.28 |
| *pmt4* | CAGL0M00220g | 0.09 | 0.1 | 0.07 | 0.07 |
| **Protein glycosylation - N-linked mannosylation** | | | | | |
| *mnn13* | orf19.4270 | 0.02 | 0 | 0 | 0 |
| *mnn14* | orf19.6996 | -0.15 | -0.01 | -0.07 | -0.08 |
| *mnn15* | orf19.753 | 0.06 | 0.03 | 0.01 | 0.03 |
| *mnn22* | orf19.3803 | -0.06 | -0.02 | -0.09 | -0.03 |
| *mnn4* | orf19.2881 | -0.09 | -0.05 | -0.05 | -0.03 |
| *mnn9* | orf19.7383 | -0.08 | 0.01 | -0.02 | -0.01 |
| *mns1* | orf19.1036 | 0.03 | 0.03 | 0.01 | 0.05 |
| *mnt4* | orf19.6313 | -0.02 | 0.02 | 0 | 0 |
| *och1* | orf19.7391 | -0.09 | -0.04 | -0.05 | -0.03 |
| *anp1* | CAGL0L01331g | -0.02 | -0.11 | -0.15 | -0.01 |
| *mnn10* | CAGL0K11231g | 0.3 | 0.06 | -0.12 | 0.24 |
| *mnn11* | CAGL0G07491g | 0.37 | 0.1 | -0.08 | 0.17 |
| *mnn14* | CAGL0C04048g | -0.25 | -0.14 | -0.18 | -0.1 |
| *mnn2* | CAGL0I04532g | 0.66 | 0.25 | 0.13 | 0.27 |
| *mnn41* | CAGL0H09130g | 0.25 | 0.04 | 0.04 | 0.11 |
| *mns1* | CAGL0M00528g | 0.01 | 0.02 | -0.04 | -0.03 |
| *rot2* | CAGL0K06963g | -0.05 | -0.15 | -0.05 | -0.09 |
| **Calcineurin pathway** | | | | | |
| *crz1* | orf19.7359 | NG | NG | NG | NG |
| *mid1* | orf19.3212 | NG | NG | NG | NG |
| *cna1* | CAGL0L11110g | NG | NG | NG | NG |
| *crz1* | CAGL0M06831g | NG | NG | NG | NG |
| *mid1* | CAGL0M03597g | NG | NG | NG | NG |

| | | | | | |
|---|---|---|---|---|---|
| **Osmotic and oxidative stress response (MAPK signalling)** | | | | | |
| *hog1* | orf19.895 | NG | NG | NG | NG |
| *pbs2* | orf19.7388 | NG | NG | NG | NG |
| *skn7* | orf19.971 | 0.13 | -0.03 | 0.1 | -0.06 |
| *sko1* | orf19.1032 | 0.09 | -0.04 | 0.05 | -0.04 |
| *pbs2* | CAGL0L05632g | -1.15 | -1.17 | -1.11 | -1.09 |
| *skn7* | CAGL0F09097g | 0 | 0.07 | 0.02 | -0.04 |
| *ssk2* | CAGL0M10829g | -0.06 | 0.05 | -0.03 | -0.05 |
| **Drug efflux** | | | | | |
| *mrr1* | orf19.7372 | 0.06 | -0.23 | 0.06 | -0.12 |
| *mrr1* gof | orf.19.7372 | -0.14 | -0.1 | -0.05 | -0.05 |
| *snq2* | orf19.5759 | -0.09 | -0.06 | -0.13 | -0.16 |
| *tac1* | orf.19.3188 | 0.21 | 0.01 | 0.14 | -0.04 |
| *tac1* gof | orf.19.3188 | -0.06 | 0 | 0.05 | 0.02 |
| *cdr1* | CAGL0M01760g | 0.13 | 1.13 | 0.79 | 0.99 |
| *pdr1* | CAGL0A00451g | 0.1 | 0.11 | 0.03 | 0.06 |
| *snq2* | CAGL0I04862g | 0.02 | -0.03 | -0.14 | -0.12 |
| **Polyamine uptake** | | | | | |
| *dur31* | orf19.6656 | -0.29 | -0.16 | -0.12 | -0.22 |
| *dur35* | orf19.5915 | -0.06 | -0.02 | -0.05 | -0.03 |

*Note:* Red-shading indicates poor growth of the mutant compared to the wildtype, green indicates better growth of the mutant compared to the wildtype. NG displays no growth of the mutant in the presence of polymer.

*Heterozygous mutation

The growth speed index was based on the following equation:

$$Growth\ speed\ index = -\log\left(\frac{M(treated)-M(untreated)}{WT(treated)-WT(untreated)}\right)$$

Values for time until half maximum absorption was reached were inserted into the above equation (M – deletion mutant, WT – respective wildtype)

### Immune cell response to polymer LH

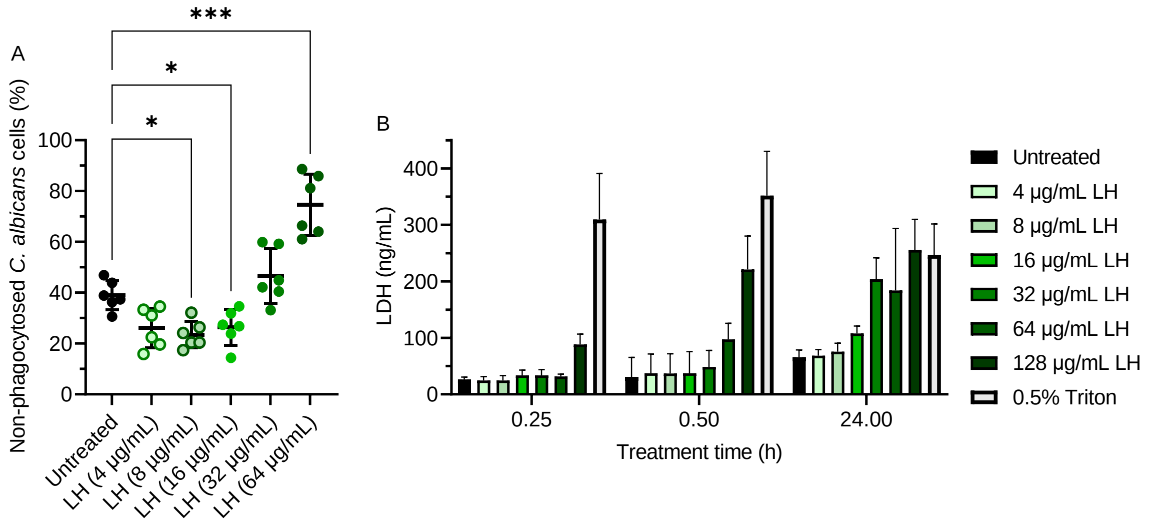

**Figure S21.** (A) Phagocytosis of *C. albicans*, pretreated with LH, by human monocyte-derived macrophages after 15-30 mins and (B) damage to uninfected macrophages measured by LDH release after 15 and 30 min and 24 h. Statistical analysis in A was performed by Dunnett's repeated measures ANOVA multiple comparison analysis. *p<0.05, **p<0.005, ***p<0.0005.

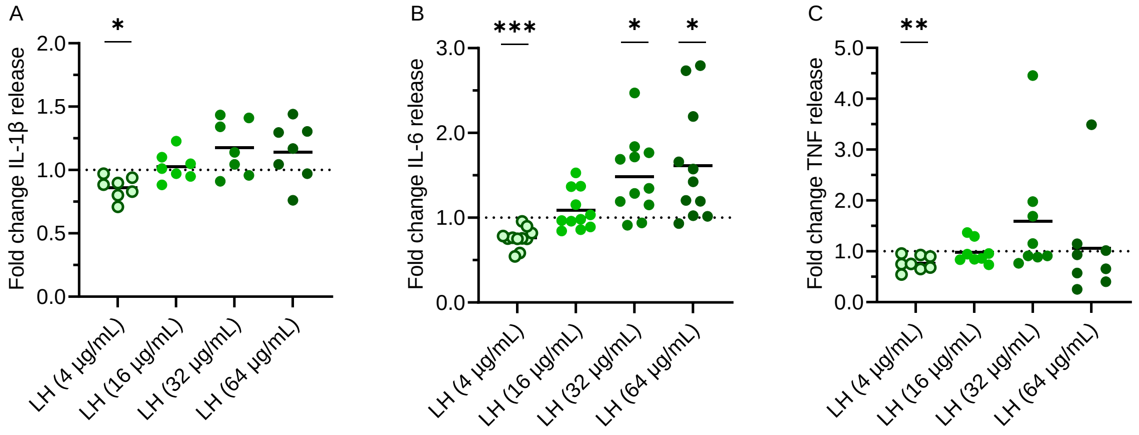

**Figure S22.** Pro-inflammatory cytokine release (A: IL-1β, B: IL-6, C: TNF) by human PBMCs after exposure to *C. albicans*, pre-treated with LH at different concentrations. Statistical analysis was performed by Dunnett's repeated measures ANOVA multiple comparison analysis and compared to untreated (fold change 1). *p<0.05, **p<0.005, ***p<0.0005.

### Hypha formation of *C. albicans* after polymer exposure

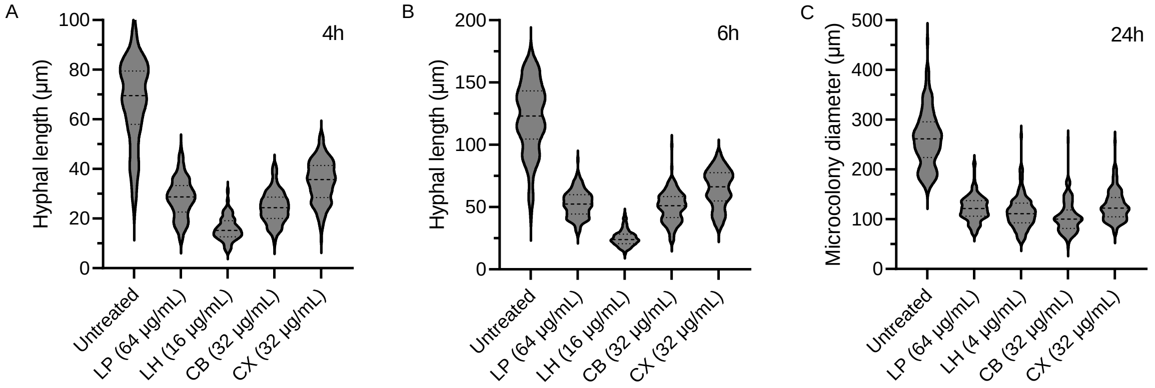

**Figure S23** – Effect of polymers at subinhibitory concentrations on hypha and microcolony formation of *C. albicans* SC5314. Hyphal growth and microcolonies were induced by incubating the fungal cells in RPMI medium at 37 °C with 5% $CO_2$. (A) Lengths of the hyphae after 4 and (B) 6 h, and (C) microcolony diameter after 24 h were measured microscopically. All polymer treatments significantly reduced hyphal length and microcolony diameter compared to the untreated control (p-value < 0.0001) according to ordinary one-way ANOVA (n=150).

#### Effect of polymer LH and established antifungal drugs on infection of vaginal epithelial cells by *C. albicans*

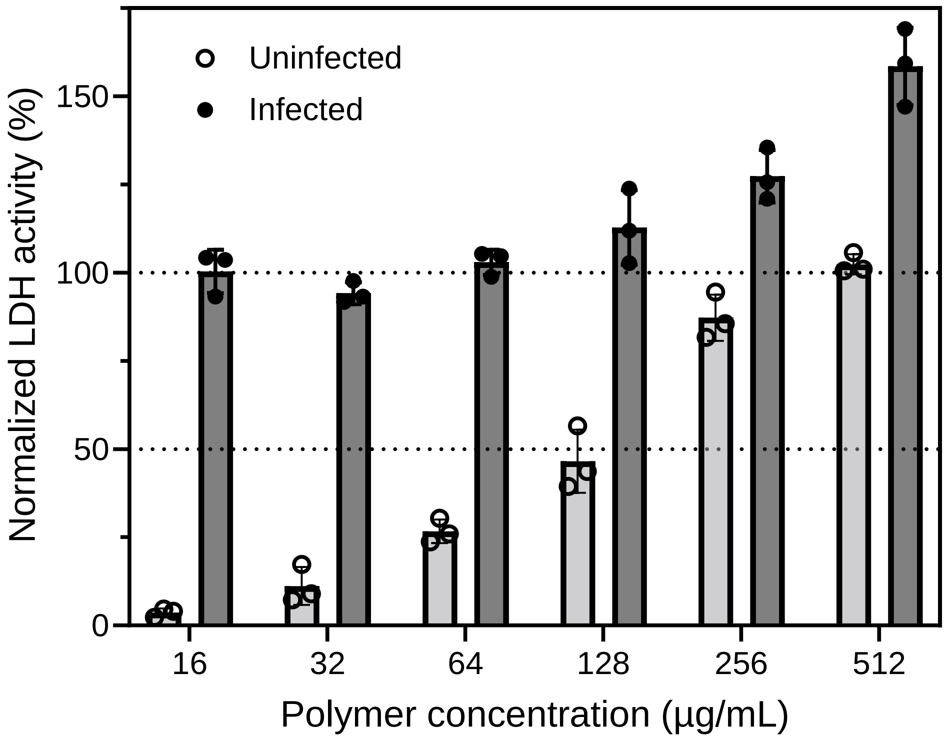

**Figure S24** – LDH activity as a measure of damage to A-431 vaginal epithelial cells after 24 h caused by *C. albicans* (infected) and the polymer LH (both infected and uninfected). For infected samples (dark grey, black circles), the LDH signal of infected controls without addition of antifungals was set to 100% and for uninfected samples (light grey, clear circles), the LDH signal of Triton-X treated vaginal epithelial cells was set to 100%.

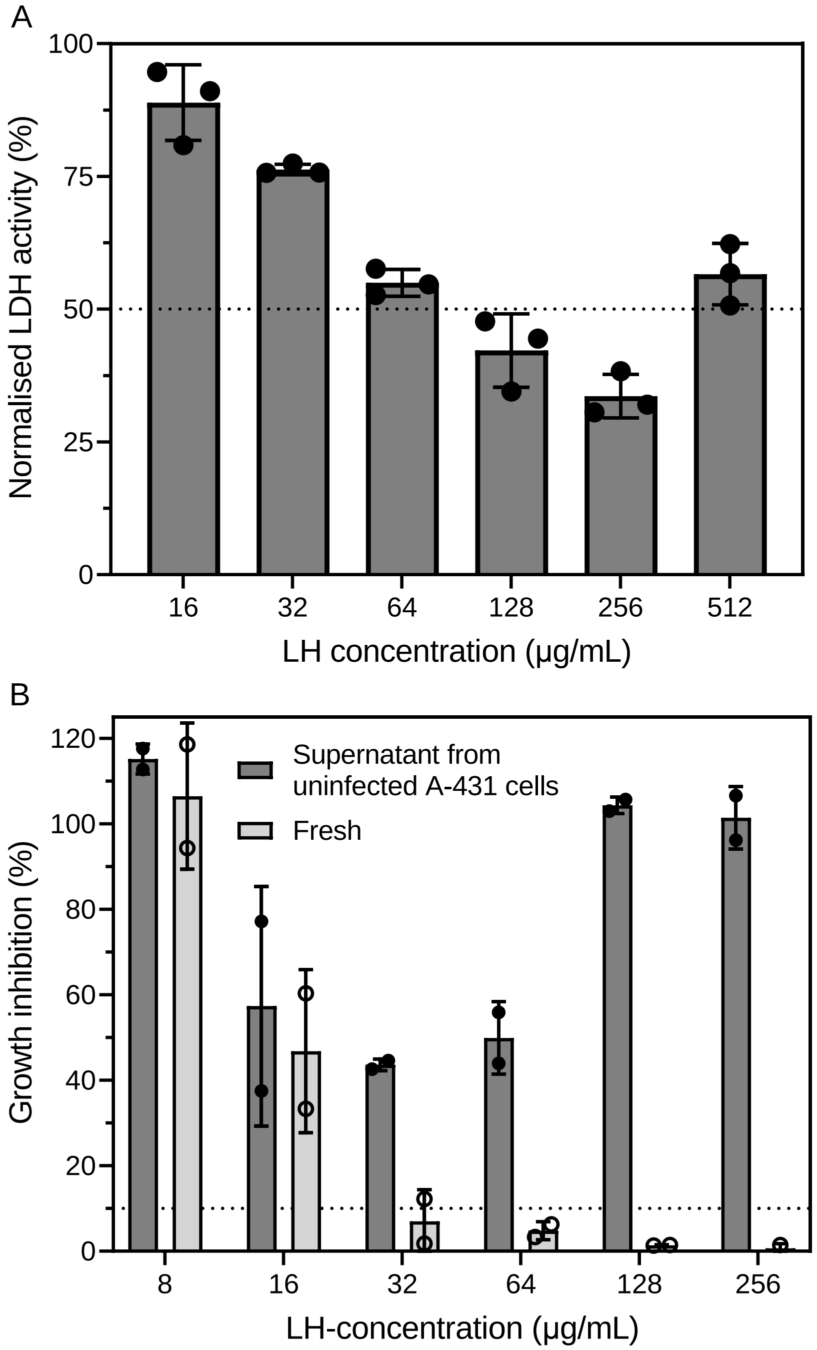

**Figure S25** – (A) LDH activity as a measure of damage to A-431 vaginal epithelial cells after 24 h caused by *C. albicans* preincubated for 60 min with polymer LH (16-512 µg/mL). The LDH signal of infected controls without addition of LH was set to 100%. (B) *C. albicans* growth inhibition after treatment with polymer LH in an MIC assay (n=2). Dark grey bars represent treatment with supernatant taken from uninfected A-431 cells after 24 h, treated with 2× displayed polymer concentrations, and light grey bars represent treatment with freshly prepared polymer solutions. Growth inhibition was normalised to untreated controls.

**Table S3** – Effective concentrations (EC) of antifungals to cause minimum 50% damage to vaginal epithelial cells A-431 after 24 h compared to lysis control (100%) and minimum concentration to decrease damage to the vaginal epithelial cells by at least 90% compared to the infection control in the *in vitro* human epithelial cell model (MIC).

| **Antifungal** | **$EC_{50\%, 24\ h}$ (µg/mL)** | **$MIC_{90\%, 24\ h}$ (µg/mL)** |
|---|---:|---:|
| Amphotericin B | 16-32 | 8 |
| Caspofungin | >2 | 0.25 |
| Cycloheximide | 1024 | 512 |
| FK506 | 64-128 | >128 |
| Fluconazole | >2 | 0.25 |
| Geldanamycin | >8 | >8 |
| Nikkomycin Z | >32 | >32 |
| Tunicamycin | 64-128 | 8-16 |

#### Synergistic effects of polymer LH and established antifungal compounds on *Candida albicans* alone

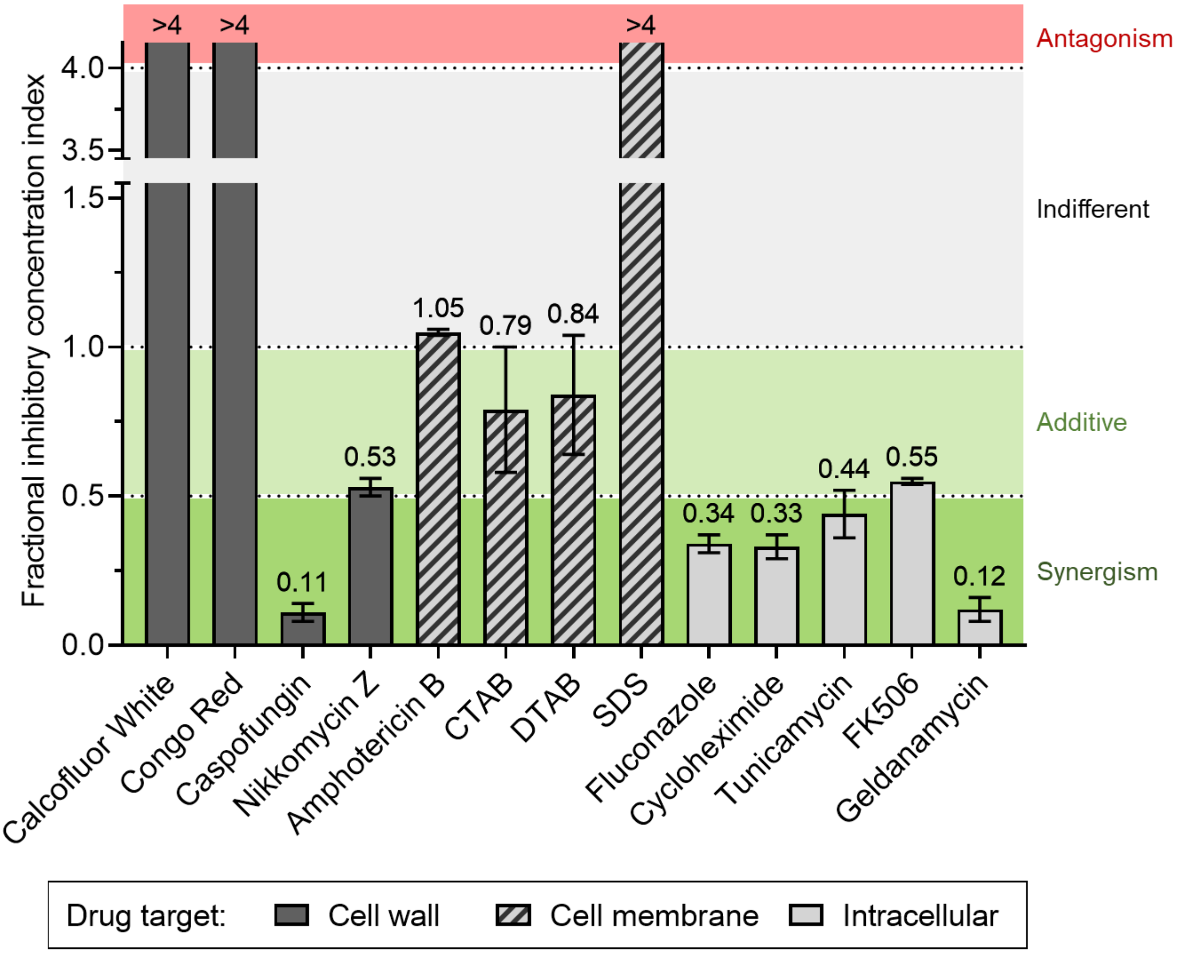

**Figure S26** – Fractional inhibitory concentration (FIC) index as a measure of synergistic (green, FIC < 0.5), additive (light green, FIC 0.5-1.0), indifferent (grey, FIC 1-4) or antagonistic (red, FIC >4) activity of the combination of polymer LH and another antifungal against *C. albicans*. The antifungals are ordered by their primary drug target, being the fungal cell wall (dark grey bars), cell membrane (striped bars), or intracellular biosynthetic pathways (light grey bars).

*For the calculation of the FIC index and individual MIC values, consult **Supplementary Table S4**.

**Table S4** – Minimum concentrations to inhibit *C. albicans* growth by at least 90% (MIC) after 24 h of the displayed antifungal compounds alone and in combination with LH. Fractional inhibitory concentration (FIC) indices are displayed including standard deviation (SD).

| **Antifungal compound** | **MIC antifungal only (µg/mL)** | **MIC in combination (antifungal, µg/mL)** | **MIC in combination (LH, µg/mL)** | **FIC index ± SD** |
|---|---|---|---|---|
| LH | 16-32 | - | - | - |
| Calcufluor White* | >512 | >512 | >128 | >4 |
| Congo Red* | 128 | >512 | >128 | >4 |
| Caspofungin | 0.25-0.5 | 0.03-0.06 | 1-2 | 0.11 ± 0.03 |
| Nikkomycin Z | 0.25 | 0.06-0.125 | 2-8 | 0.53 ± 0.03 |
| Amphotericin B | 1 | 0.5-1 | 16-32 | 1.05 ± 0.01 |
| CTAB | 2 | 1 | 4-16 | 0.79 ± 0.21 |
| DTAB | 32 | 2-16 | 16 | 0.84 ± 0.20 |
| SDS | 128 | >1024 | >128 | >4 |
| Fluconazole | 0.25-0.5 | 0.06-0.125 | 2-4 | 0.34 ± 0.03 |
| Cycloheximide | 256-512 | 32-64 | 4-8 | 0.33 ± 0.04 |
| Tunicamycin | 4 | 1-2 | 2-8 | 0.44 ± 0.08 |
| FK506 | 128-256 | 4-64 | 4-8 | 0.55 ± 0.01 |
| Geldanamycin | 4-8 | 0.125-0.25 | 2-4 | 0.12 ± 0.04 |

*determined visually

The FIC index was calculated as described in the following equation, where $c_{A/B}$ are the concentrations of compounds A or B, respectively, in combination resulting in growth inhibition >90%, and $MIC_{A/B}$ are the MICs of compound A or B, respectively, alone:

$$FIC\ index = \frac{c_A}{MIC_A} + \frac{c_B}{MIC_B}$$

The anionic, fluorescent dyes Calcofluor White and Congo Red are used in microscopy to stain the cell wall of fungi due to their polysaccharide-binding activity. At relatively high concentrations, they inhibit the growth of *C. albicans* (**Supplementary Table S4**). A combination of Calcofluor White or Congo Red with LH caused antagonistic behaviour which could be a consequence of the opposite charges of the molecules and putatively similar drug targets. Caspofungin and nikkomycin Z interfere with cell wall integrity by targeting the glucan- and chitin-synthase, respectively. While nikkomycin Z showed an additive effect on *C. albicans* growth inhibition in combination with LH (FIC index 0.53), caspofungin showed strong synergism (FIC index 0.11).

The polyene antifungal AmpB and the positively charged surfactants cetyltrimethyl ammonium bromide (CTAB) and dodecyltrimethylammonium bromide (DTAB) showed indifferent (AmpB, FIC index 1.05) or slightly additive effects (CTAB – 0.79; DTAB – 0.84) in combination with LH. In contrast to the positively charged surfactants, the negatively charged surfactant sodium dodecyl sulfate (SDS) caused antagonistic behaviour in combination with LH, possibly due to the opposite charges of the molecules.

Combining LH with antifungals with an intracellular target shows at least strong additive (FK506, FIC index 0.55) or even synergistic effects (fluconazole, FIC index 0.34; cycloheximide, FIC index 0.33; tunicamycin, FIC index 0.44; geldanamycin, FIC index 0.12). The synergistic effects of LH with antifungals targeting intracellular processes and caspofungin which targets the glucan synthase supports the observation of LH to interfere with the fungal cell wall primarily, which could facilitate the delivery of the antifungal drugs to their target.

#### Testing polymer LH-drug combinations for synergistic effects on *C. albicans* infection of human epithelial cells *in vitro*

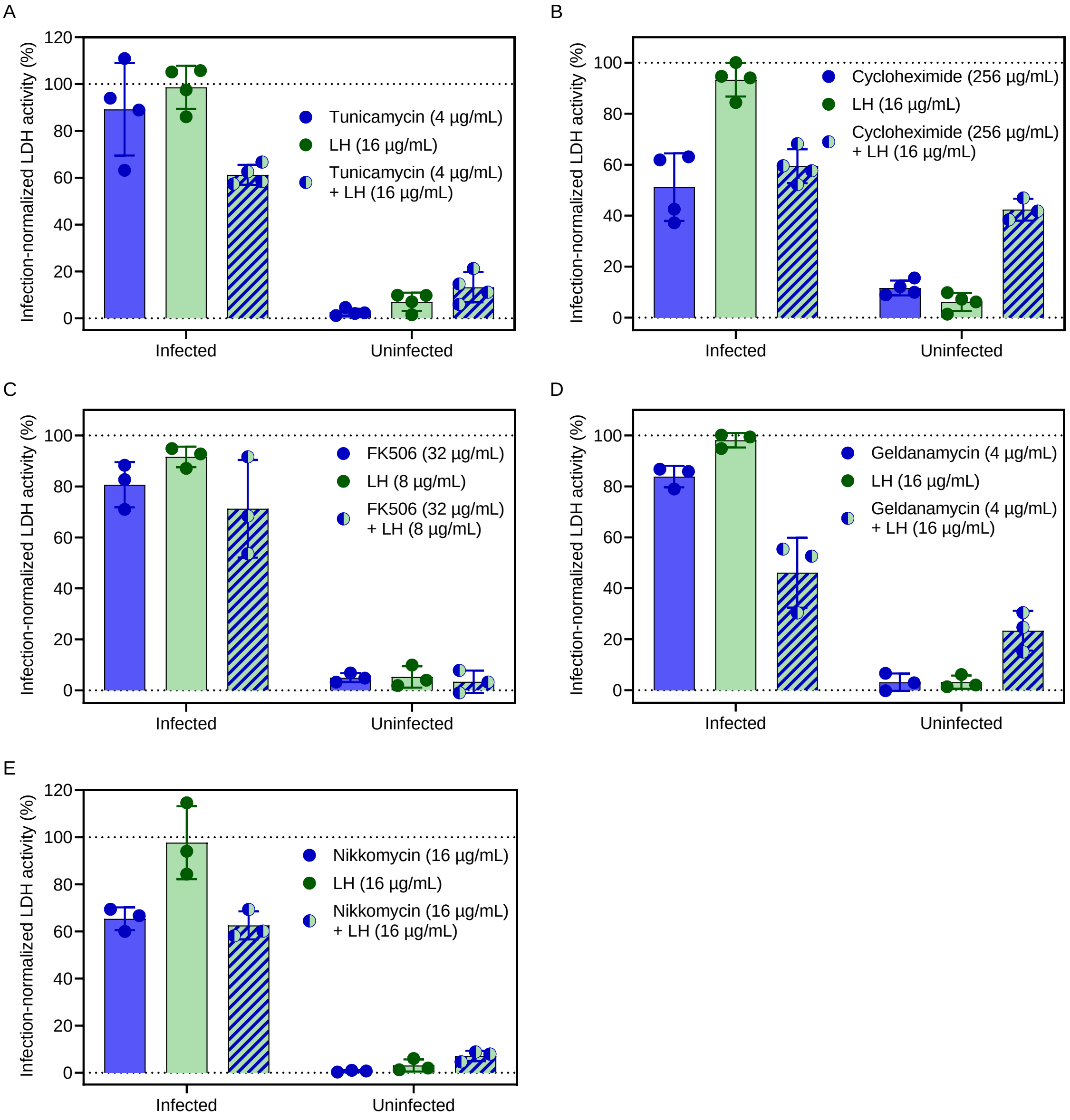

**Figure S27** – Damage to vaginal epithelial cells, co-incubated with *C. albicans* (infected) or uninfected, after 24h treatment with the polymer LH, the antifungals (A) tunicamycin, (B) cycloheximide, (C) FK506, (D) geldanamycin, (E) and nikkomycin Z and their respective combinations to investigate synergistic effects.

#### *In vivo Galleria-C. albicans* infection model

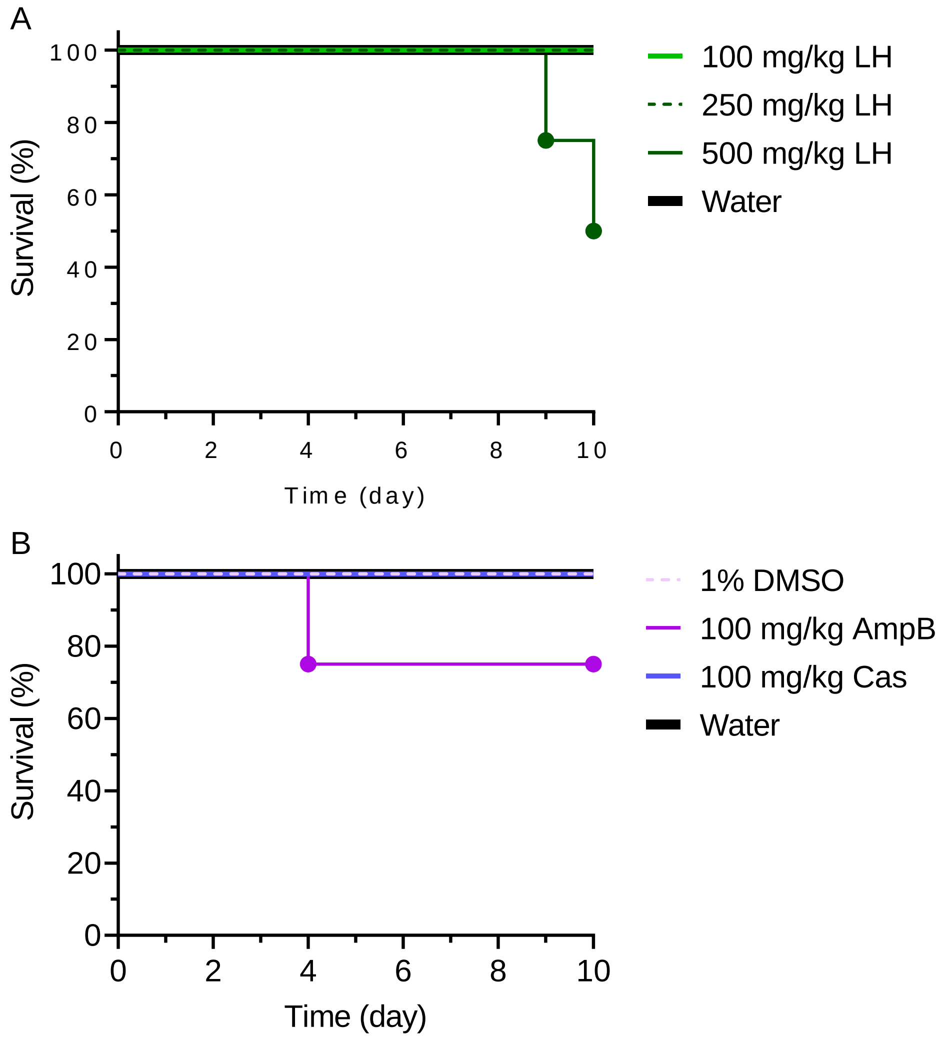

**Figure S28** – Acute toxicity of (A) polymer LH at different concentrations (100 mg/kg light green, 250 mg/kg green dashed, 500 mg/kg dark green), and (B) the antifungal drug controls caspofungin (Cas, blue) and amphotericin B (AmpB, purple) against uninfected *Galleria mellonella* larvae (n = 4 larvae per group).

**Table S5** – Visual MIC of polymer LH, fluconazole (Flu), and their combination against *C. albicans* in modified RPMI medium, supplemented with different ratios of FBS (0-100% (v/v)) (n=2). n. d. indicates no detected synergistic action.

| FBS (%) | LH (µg/mL) | Flu (µg/mL) | LH + Flu (µg/mL) |
|---|---|---|---|
| **0** | 16 | 0.5 | 4 + 0.125 |
| **5** | 128-256 | 1-2 | 8 + 0.125 |
| **50** | >256 | 1-2 | n. d. |
| **100** | >256 | 0.25-0.5 | n. d. |

### *C. albicans* evolution assay

**Table S6** – Minimum concentrations to inhibit 90% growth of *C. albicans* strains after evolution assay.

| ***C. albicans* strain** | **MIC$_{24\,h,\,90\%}$ (µg/mL)** | | | | | | | |
|---|---|---|---|---|---|---|---|---|
| | LP | LH | CB | CX | AmpB | Flu | Cas | TM |
| SC5314 (t$_0$) | 128-256 | 16-32 | 64 | 64-128 | 1 | 0.5 | 0.25-0.5 | 2 |
| evo-SC5314 (14 days in RPMI, pH 4) | 256 | 16-32 | 64 | 64-128 | 1 | 0.5 | 0.25-0.5 | 2-4 |
| evo-LH1 | >512 | 64-128 | 256-512 | 256-512 | 2 | 0.5-1 | 0.25-0.5 | 2 |
| evo-LH2 | 256-512 | 64 | 128 | 128-256 | 1-2 | 0.5 | 0.25 | 2 |
| evo-LH3 | 256-512 | 32 | 128 | 128-256 | 1-2 | 0.5 | 0.25 | 2 |
| evo-LH+Cas1 | 128 | 16 | 64-128 | 128 | 1 | 0.5 | 0.5 | 2 |
| evo-LH+Cas2 | 128-256 | 16 | 64 | 128 | 1 | 0.5 | 0.5 | 2 |
| evo-LH+Flu1 | 128 | 8-16 | 32-64 | 32-64 | 0.5-1 | 1 | 0.5 | 1-2 |
| evo-LH+Flu2 | 128 | 16 | 32 | 32-64 | 0.5-1 | 1 | 0.5 | 2 |
| evo-Cas1 | 128-256 | 16 | 64 | 64-128 | 1 | 0.5 | 2 | 2 |
| evo-Flu1* | >512 | 256 | 256-512 | 512 | 2 | 2 | 2-4 | 1-2 |

*Note:* Yellow reflects an increase in MIC by twofold, and orange reflects a fourfold or higher increase in MIC. AmpB - amphotericin B, Flu – fluconazole, Cas – caspofungin, TM – tunicamycin, * Flu-treated strains needed 48 h to reach visible growth even at the untreated condition wherefor the MIC was measured after 48 h instead of 24 h for this evolved strain.

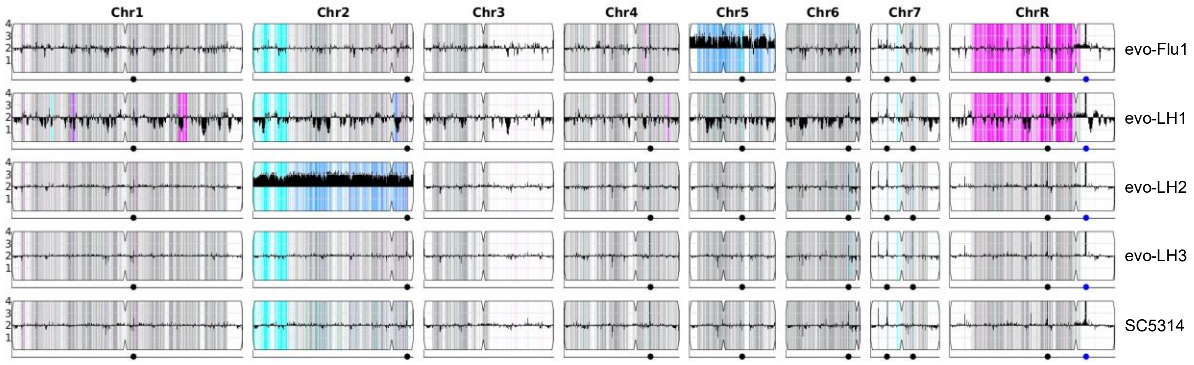

**Figure S29** – Genomes of selected *C. albicans* isolates with increased LH tolerance in comparison to wild-type SC5314, based on YMAP[1]. The black graph displays the relative gene count compared to SC5314 for a gene (Y-axis) on its chromosome position (X-axis). Cyan shading represents loss of heterozygosity (LOH) of allele A, pink represents LOH of allele B, grey presence of both alleles A and B, dark blue shading represents trisomy.

### References

1. Abbey, D.A. et al. YMAP: A pipeline for visualization of copy number variation and loss of heterozygosity in eukaryotic pathogens. *Genome Medicine* **6**, 100 (2014).
